# A Cattle BodyMap of Transcriptome across 52 Tissues and 3 Developmental Stages Reveals New Genetic Insights into Beef Production Traits

**DOI:** 10.1101/2025.07.21.665934

**Authors:** Wentao Cai, Yapeng Zhang, Lei Xu, Yahui Wang, Xin Hu, Qian Li, Linxi Zhu, Zezhao Wang, Huijiang Gao, Lingyang Xu, Junya Li, Lupei Zhang

## Abstract

Cattle are important livestock that provide essential meat and milk resources. However, a comprehensive analysis of gene expression, alternative splicing (AS), and RNA editing across various organs and developmental stages in cattle has not been reported. This study aims to create a comprehensive transcriptomic BodyMap across various tissues and developmental stages, integrating this information into genomic predictions of beef production traits. We created a comprehensive transcriptomic BodyMap using 400 samples collected from 52 organs of newborn, young, and adult cattle, and estimated their contributions to genetic variance and genomic predictions for 23 beef production traits in 1476 beef cattle. We cataloged the expression of 25,530 annotated genes, 28,533 novel long non-coding RNAs (lncRNAs), 215,754 AS events, and 3,093,058 A-to-I RNA-editing sites. Integrating transcriptome BodyMap with 23 beef production traits, we found lncRNAs influenced traits like rib-eye area and carcass length, while RNA editing associated with chunk roll weight and daily gain. We observed trait-relevant tissues between different stages, including the differential expressed genes of cerebellum, longissimus muscle, and testis between newborn and adult stages are more relevant to beef production traits. The tissue-specific genes and development-associated genes in several tissues could improve the reliability of genomic prediction in beef production traits. We developed Cattle BodyMap Transcriptome Database (https://cattlegenomics.online/cattle_bodymap) to retrieve, analyze, and visualize gene expression, lncRNA, splicing and RNA editing data. Our results demonstrated the potential of using transcriptome data as a valuable resource for genomic selection and breeding programs in beef cattle. Additionally, our transcriptome BodyMap serves as a valuable resource for biological interpretation, functional validation, and genomic improvement in livestock.

## Introduction

*Bos taurus*, specifically cattle, are essential livestock that provide vital meat and milk resources for humans. Systematic analysis of the transcriptome plays a central and crucial role in the research community focused on cattle. The comprehensive characterization of the mammalian transcriptome has significantly advanced our understanding of regulatory mechanisms and genome complexity. This has been achieved through the collaborative efforts of various projects such as the ENCODE [1], FANTOM5 [2], and GTEx [3] projects in humans, BodyMap database in rats [4] and pigs [5], as well as the FAANG [6] and FarmGTEx [7] project in livestock.

Although some progress has been made in multi-tissue transcriptome studies in cattle [8, 9], the majority have focused on the adult stage, which limited exploration of the transcriptomic BodyMap during other developmental stages and its changes across development. In addition to protein-coding genes (PCGs) that affect phenotypes, mammalian genomes also contain long noncoding RNAs (lncRNAs) [10, 11], alternative splicing (AS) [12, 13], RNA editing [14, 15], which play crucial regulatory roles through various mechanisms, contributing to the complexity and diversity of the transcriptome and influencing tissue physiology [16]. Tissue-specific and stage-specific characteristics of AS contribute to the generation of significant regulatory and proteomic complexity in mammals, thereby introducing a crucial layer of gene regulation that influences numerous biological pathways. Misregulation of AS often causes or contributes to complex human traits and diseases [17, 18]. Therefore, there is a significant demand in the research community for easily accessible AS profiling data across the most extensively studied tissue types and developmental stages. RNA editing can lead to alterations in missense codons [19], influence splicing activity or other regulatory RNA [20–22], and modify seed sequences or targeting sites of miRNAs [23]. However, the global pattern of AS and RNA editing across tissues and developmental stages remains largely unknown in cattle. Using RNA-Seq to catalog the variations in the transcriptome among tissues and over development of the cattle, from birth to adult, can not only enhance cattle genetic improvement program by leveraging prior information of tissue, but also provide insights into establishing cattle as a potential biomedical model to human research [24].

In this study, to create a comprehensive transcriptomic BodyMap of beef cattle, we employed RNA-Seq to extensively document transcriptomic profiles across 52 diverse tissues and three developmental stages (newborn, young, and adult). These tissues were categorized into 13 main organ/tissue types based on common developmental, functional, and anatomical properties (Figure 1). We sequenced 400 pair-end RNA-Seq libraries (150 bp) and generated ∼ 20.4 billion reads. We summarized the details of sample information in Table S1.

**Figure 1.**
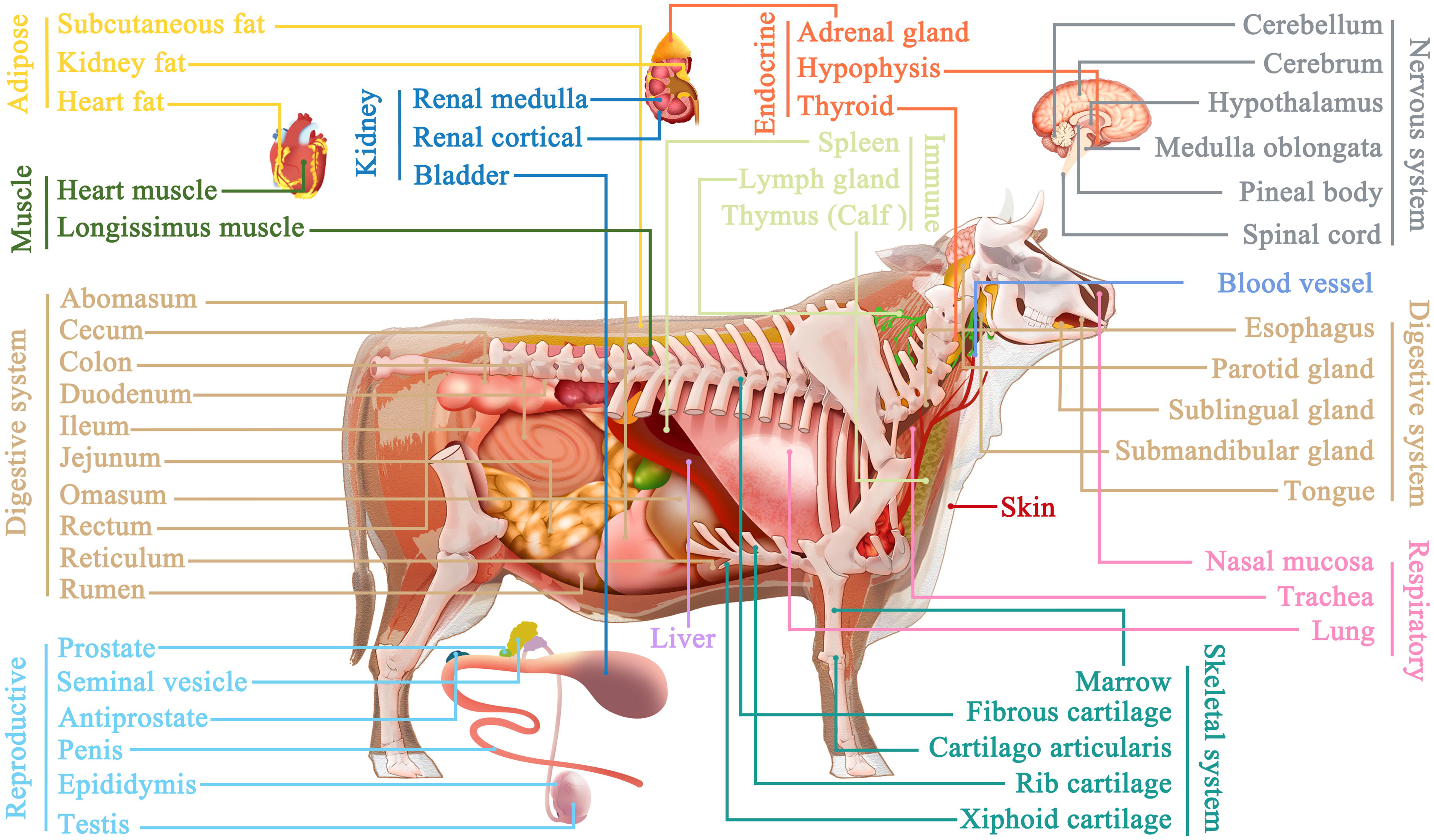
Samples derived from 52 tissues across three developmental stages were used for ab initio cattle BodyMap transcriptome construction.

## Results

### Landscape of cattle genes

Considering all tissues across three stages, we discovered 20,826 PCGs, 1474 lncRNAs, and 3230 other genes, accounting for 95.3%, 99.6%, and 80.3% of all known PCGs, lncRNAs and other genes, respectively, with the fragments per kilobase per million mapped reads (FPKM) > 0.1. We identified 28,533 novel lncRNAs (144,363 transcripts), including 78.7% intergenic lncRNAs (lincRNAs) and 21.3% intron lncRNAs. Compared with PCGs, the lncRNAs were shorter in length (mean: 1.10 kb vs 2.47 kb, Figure S1a), had fewer exons (mean: 2.59 vs 11.25 exons per transcript, Figure S1b), exhibited relatively frequent events of AS (mean: 4.88 vs 1.72 isoforms, Figure S1c), lower evolutionarily conserved (mean PhastCons score: 0.15 vs ∼0.65, Figure 2a), low expression (mean FPKM: 10.61 vs 25.38, Figure S1d), and high tissue specificity in their expression (95.28% vs 70.26% with τ ≥ 0.75, Figure 2b).

**Figure 2.**
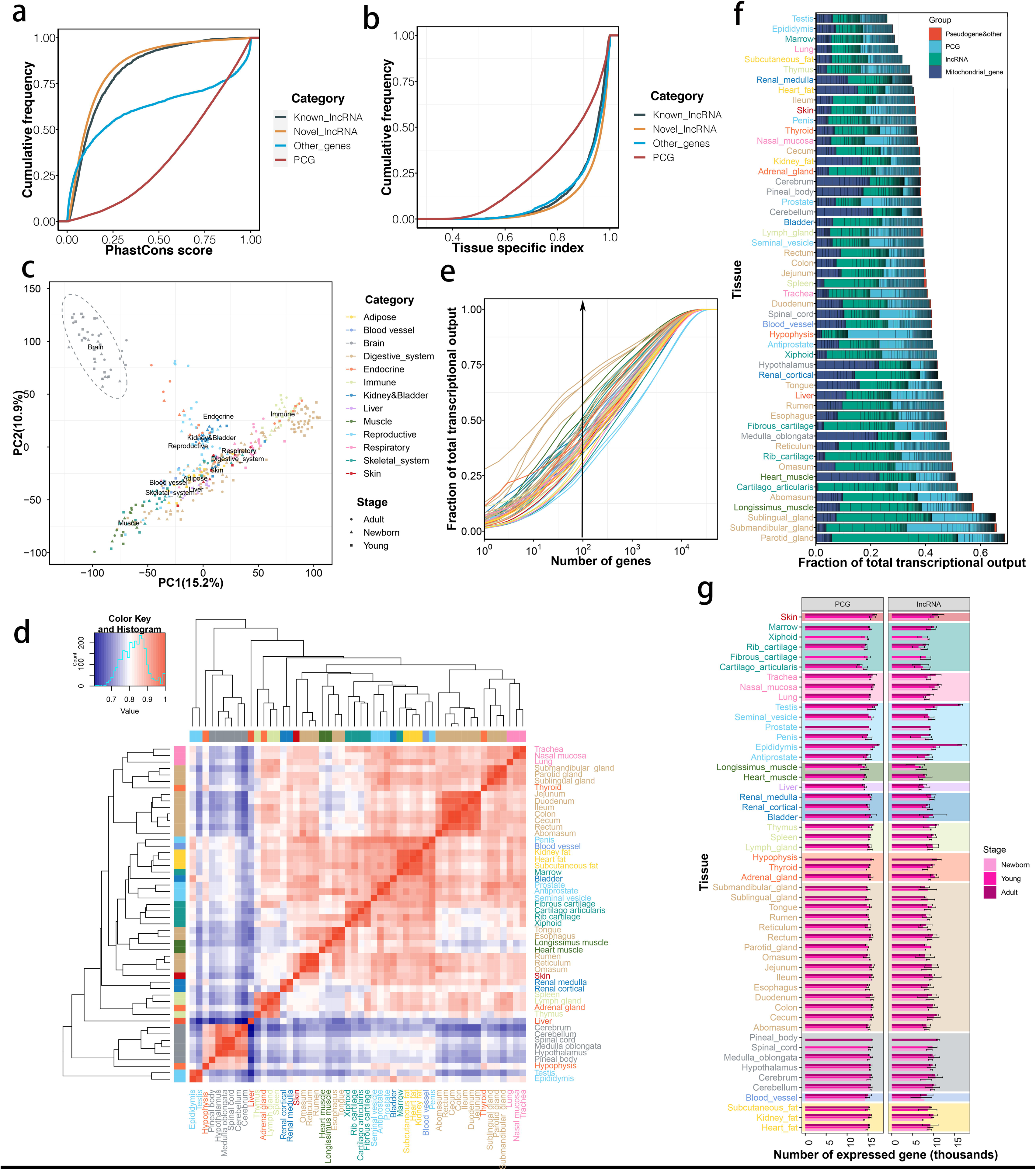
Characteristics of the cattle BodyMap transcriptome. (a) Sequence conservation of four transcript types in the cattle transcriptome. The base (nucleotide resolution) phastCons scores were collected from the UCSC Genome Browser. The transcript level phastCons scores were calculated as their average value among exonic sequences. (b) Tissue specificity of four transcript types reflected by tau (τ) score. (c) Sample (n = 400) clustering using principal component analysis (PCA) based on gene expression levels. (d) Pearson correlation (heatmap) and hierarchical clustering (tree) of transcriptome profiles across 400 samples (rows/columns) show tissue-specific clustering (colors). (e) Numbers of the expressed genes in different cattle tissues across stages. The error bars are data standard deviations. (f) Cumulative distribution of the average fraction of total transcription contributed by genes when sorted from most to least expressed in each tissue (x-axis). (g) Biological type and relative contribution to total transcription of the hundred most expressed genes. The height of the bars is proportional to the fraction that these genes contribute to total transcription.

Samples of similar tissues clustered together based on both PCG and lncRNA expression profiles, the brain was the most distinct (Figure 2c). The Spearman correlation was utilized in a pairwise correlation heatmap for the 52 tissues to delve deeper into similarities in global transcriptome profiles across tissues (Figure 2d and Figure S2). The accompanying dendrogram revealed that related tissues tended to cluster together, encompassing tissues from the immune, respiratory, digestive, and brain systems (Figure S3). The testis and epididymis, with a large enrichment of germ cells, were clustered separately. We observed that in most tissues, the transcription of only a few hundred genes accounted for 50% of the total transcription (Figure 2e). We detected the majority of transcription was mitochondrial origin in the brain, heart, and fat tissues (Figure 2f). In general, we found that an average of 15,306 (70%) PCGs and 9110 lncRNAs (30.4%) were defined as expressed (FPKM > 0.1) per organ. The adult male reproductive tissue, such as epididymis and testis had the highest numbers of expressed genes, while the liver and muscle had the lowest numbers of expressed genes (Figure 2g). We detected 9,113 PCGs (41.7%) and 1,435 lncRNAs (4.8%) were expressed in all the 52 tissues at all developmental stages. These commonly expressed PCGs appear to be primarily involved in basic biological functions, such as metabolic pathways, spliceosome, oxidative phosphorylation, and cell cycle (Figure S4 and Table S2).

### Tissue- and development-dependent genes

We detected thousands of genes with differential expression between tissues at each developmental stage (Figure S5-S10). Compared with other tissues, brains had more differentially expressed genes (DEGs) in all three stages, while the testis was different in the adult stage with an average of 62.1% PCGs differentially expressed (Figure 3a). Preferentially expressed analysis revealed that many genes were preferentially highly expressed in the brain tissues of newborn and young stages, while this preference was decreased at adult stage (Figure S11). Conversely, the number of testis genes with preferentially high expression increased with age. Nearly two thousand genes were preferentially expressed at low level in newborn thyroid. The thymus and hypothalamus had more genes preferentially expressed at low levels in young bulls, while the cartilago articularis had more genes preferentially expressed at low levels in adult bulls. More genes were exclusively expressed in epididymis and testis with relatively high φ (Phi) values (Figure 3b and Figure S13), and 27 genes were expressed exclusively in skin, pineal body, liver, and heart muscle with *φ* >= 0.95 (Figure S12). Hundreds of cartilage genes exclusively disappeared during the adult stage (*φ* <= -0.95). Focused on each stage, we detected several genes of the cerebrum and cerebellum that were exclusively expressed during the early stage, while several testis genes were exclusively expressed after the young stage (Figure S12).

**Figure 3.**
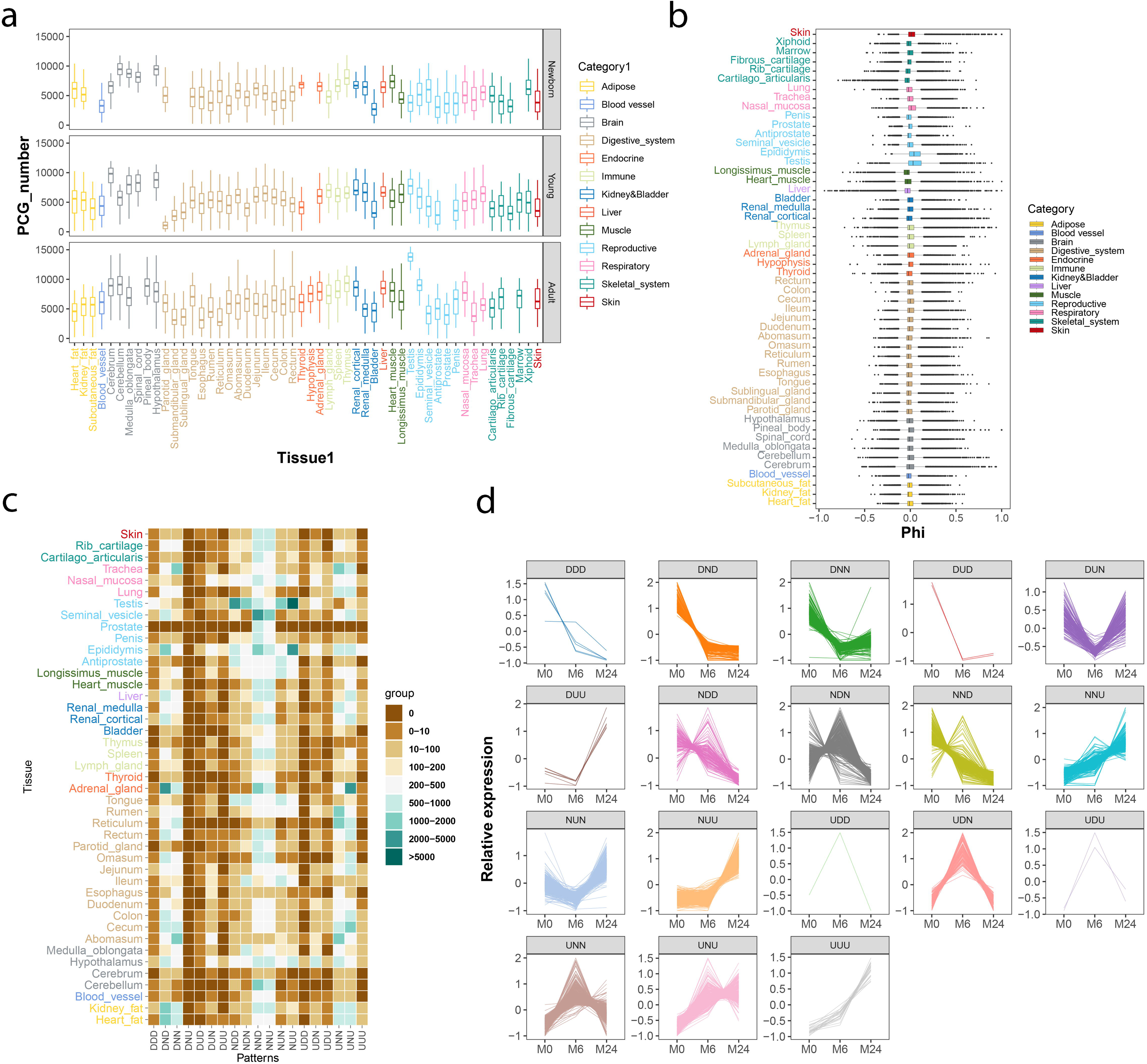
Tissue- and development-dependent genes. (a) The number of protein-coding genes (PCGs) in differentially expressed genes (DEGs) for across tissues at three development stages. (b) Genes with tissue-exclusive expression. The distribution of the phi correlation is given for all genes within each tissue. (c) The patterns of differentially expressed PCGs within each tissue. (d) The patterns of differentially expressed PCGs in muscle.

We identified 18,897 PCGs, 13,967 lncRNAs, and 1764 other genes differentially expressed across ages in at least one tissue. The number of development-dependent genes varied by organ, from 274 in the cerebrum to 24,293 in the testis (Table S3). As expected, a significant number of genes were differentially expressed in the testis with ages. The smallest number of DEGs was observed in the renal medulla of young 6-month-old cattle compared with adult cattle. The expression of the digestive tract became more distinct from other tissues from newborn to adult. To evaluate the transcriptomic activities that were dependent on development, we conducted a time course analysis based on DEGs by comparing any two developmental stages for each tissue (Young vs. Newborn, Adult vs. Young, and Adult vs. Newborn). These potential patterns included those that consistently increased across all developmental stages, referred to as ‘up-up-up’ (UUU), and those that consistently decreased across all developmental stages, referred to as ‘decrease-decrease-decrease’ (DDD). Additionally, there were genes with expression levels that remained stable across developmental boundaries, termed ‘non-change’ (N). The overall development-dependent patterns across all organs were shown in Figure 3c. Relatively few genes continuously increased (UUU) or decreased (DDD) in expression with aging. The NND and NNU patterns more frequently occurred meaning that most DEGs were only differentially expressed between newborns and adults. The prostate was relatively stable during development, except for hundreds of gene changes from newborn to adult. The hypothalamus had a variety of patterns, which implied many biological activities produced in the hypothalamus during development. The male reproductive were dramatically changed between young and adult. Interestingly, the adrenal gland only dynamically changed before young, which implied its specific function in the newborn stage. We found similar patterns in lncRNAs (Figure S14). The example patterns of differentially expressed PCGs in muscle were shown in Figure 3d.

### Landscapes of AS in cattle

We uncovered 215,754 AS events across all 400 samples, including skipping exon (SE), alternative 5’ splice sites (A5SS), alternative 3’ splice sites (A3SS), intron retention (IR), and mutually exclusive exons (MXE). SE was the most prevalent type of AS event in cattle transcriptomes (Figure S15). We found the respiratory tissues had more AS events, while AS events were relatively limited in brains (Figure 4a). Comparison to Ensembl annotations, 96.3% of AS events were newly discovered. Most SE were found in only a small proportion of the samples, while MXE was more common in a large proportion of the samples (Figure 4b). When alternative exons were binned according to the percentage of samples in which they were alternatively spliced, we found a bimodal distribution in 15% ∼ 40% of the samples progressively converted into a unimodal distribution centering on a median percent spliced-in (PSI) 0.5 in > 80% of the samples. The widely occurred AS events across samples (called PanAS events) were enriched in genes that were involved in metabolic, endocytosis, focal adhesion, and autophagy pathways (Table S4a). In contrast, switchlike AS events (events with a PSI > 0.9 or PSI < 0.1 in > 80% of the samples, but with a range of PSIs ≥ 0.8 across samples) were enriched in genes that were involved in the Rap1 signaling, phosphatidylinositol signaling system, axon guidance, inositol phosphate metabolism, and Fc gamma R-mediated phagocytosis pathways (Table S4b).

**Figure 4.**
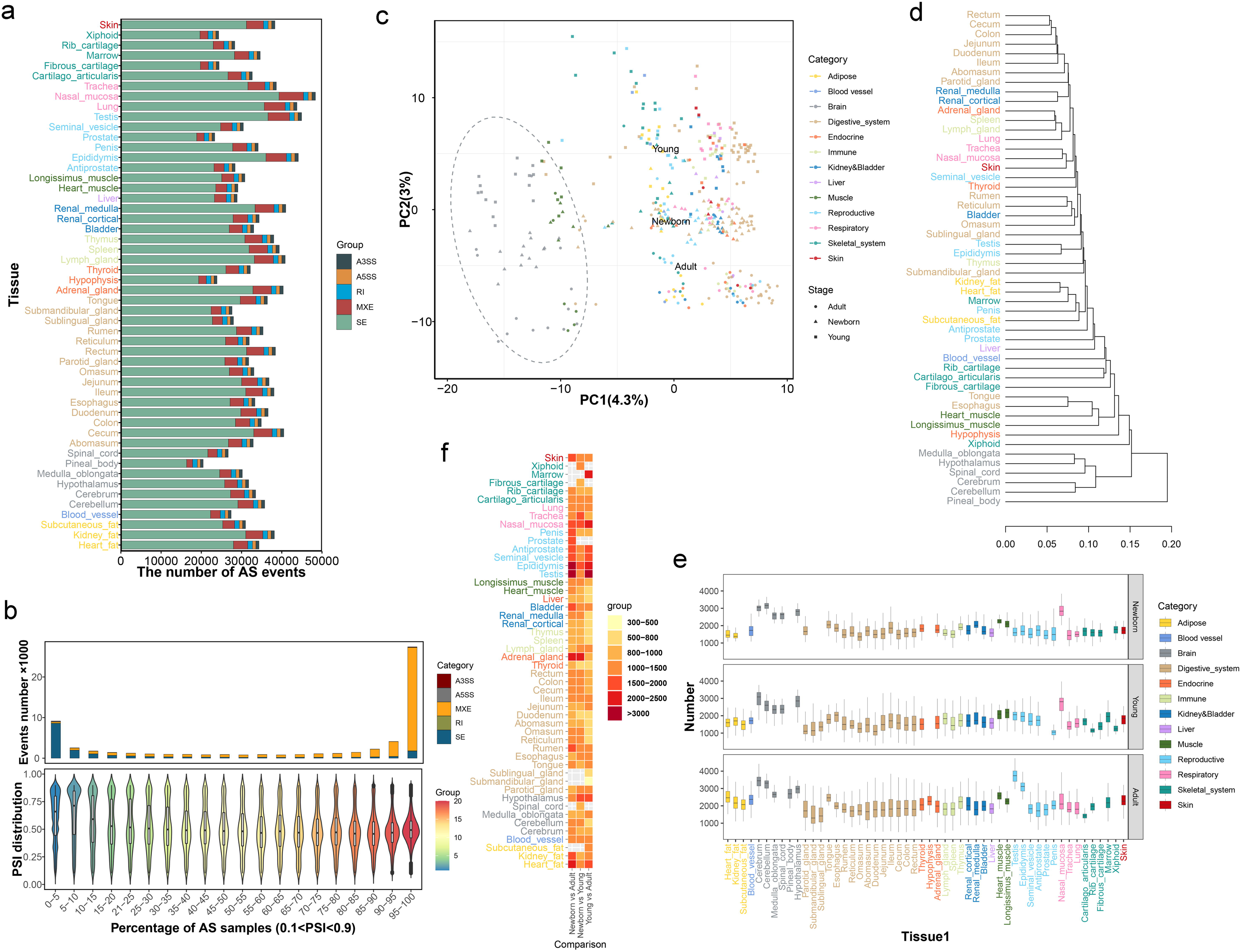
The character of splicing events in multi-tissues across stages. (a) The number of splicing events in 52 tissues at three development stages. (b) Distribution of PSI values across 400 samples with five splicing types (violin plots, bottom) and the total number of exons (histograms, top) for bins of alternative exons that are alternatively spliced (0.1< PSI < 0.9) in an increasing fraction of samples. (c) Sample (n = 400) clustering using principal component analysis based on percent spliced-in (PSI) values. (d) Tissue hierarchical clustering. Dendrogram based on pairwise Spearman correlation between tissue types. The label colors are annotated based on common biological features. (e) The differentially expressed splicing across tissues at three development stages. (f) The heatmap of numbers of the differentially expressed splicing events between each stage.

PCA based on PSI again largely recapitulated tissue types and development stages. The brain samples were primarily clustered out-group, indicating the existence of a distinct splicing program in cattle brains (Figure 4c and Figure S16). Samples from similar tissues clustered together (Figure 4d). Different organs could be distinct by development stages, suggesting strong commonalities for organ splicome at each stage. We found thousands of significantly differential AS events between tissues (Figure S17-S19). Consistent with previous human studies, the brains were distinctive in all development (Figure 4e). The testis and epididymis were distinctive at the adult stage, while the nasal mucosa was more distinctive at the newborn and young stages. We observed brain-specific splicing patterns might result from the specific up-regulation of several RNA-binding proteins, such as ELAV-like protein (ELAVL) family, A2BP1, and NOVA family, while Y box binding protein 1 (YBX1) was specifically up-regulated in the muscle organs, which might cause their exclusively splicing patterns (Figure S20). We found 227 splicing events that were exclusively included or excluded in a particular tissue, 89.4% of them being mutually exclusive splicing (Table S5). These exclusive splicing were mainly concentrated in the cartilago articularis of newborn and young stages, newborn brain, and young digestive tissues. When we focused on development-dependent splicing activities, we found that splicing activities in male reproductive tissues were more active during development (Figure 4f). The skin, adrenal gland, and fat tissues were dynamically changed between newborn and young stages. The number of genes with differential splicing events ranged from 1159 in the submandibular gland to 4850 in the testis in the Adult vs. Young comparison group (Figure S21). The number of genes with differential splicing events between stages in each tissue was highly correlated with their corresponding DEGs (Figure S22).

### Landscape of A-to-I RNA-editing

A-to-I (G, guanosine) editing was the predominant type of RNA editing in cattle, and occupied 98.9% of total editing sites (Figure S23). Our analysis identified 3,093,058 A-to-I RNA-editing sites within the cattle genome, 99% of A-to-I RNA-editing sites were clustered together in 164,214 cluster regions. The average length of the RNA editing cluster regions was 146.6 bp and contained 18.7 editing sites (Figure S24b). We found a notable depletion of G in the −1 base position and an enrichment of G in the +1 base position, which agreed with the Adenosine Deaminase Acting on RNA (ADAR) preference target sequences (Figure 5a). Most sites were in introns (43.4%) and intergenic regions (33%), while only 0.6% in protein-coding regions (Figure 5b). We detected that 93.7% edited sites were within repetitive elements. Of which, 87.4% of all A-to-I editing sites were in BovB LINEs and BovB-derived SINEs (Bov-tA2, Bov-tA1, BOV-A2, Bov-tA3), followed by 4.3% in the LINE1 elements (Figure 5c). The RNA editing cluster regions harbored dsRNA structures: 56.5% of clusters showed the reverse-complement alignment within their flanking (± 2 kb) genomic sequence (≥ 70% identities, and 60% length of the region; see Methods), compared with only 8.4% (P-value < 2.2e-16, χ^2^-test) for the random regions of similar length. The dsRNA structures were more common in BovB-derived SINEs (Figure 5c).

**Figure 5.**
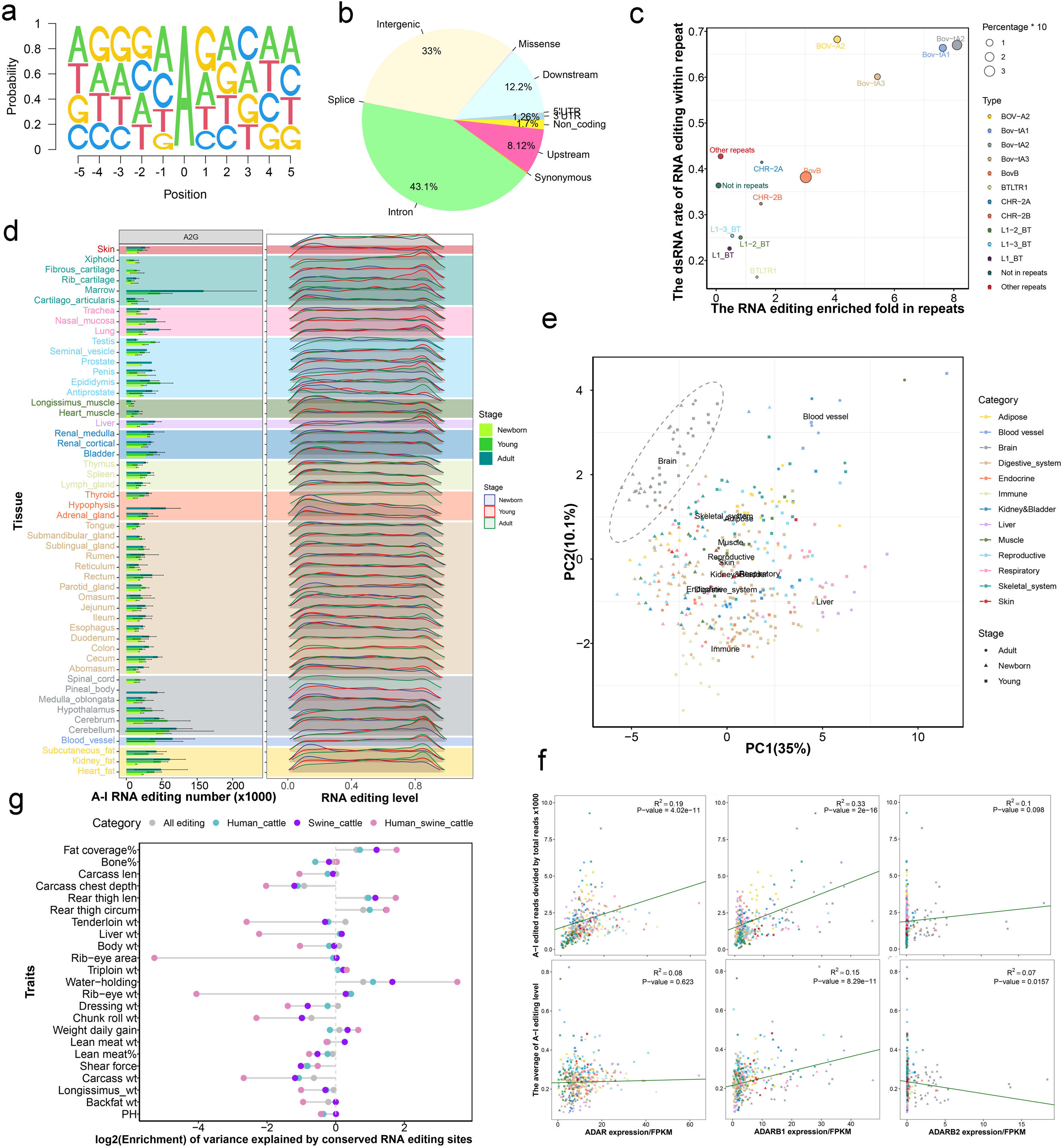
The character of RNA editing events in multi-tissues across stages. (a) The preference of nucleotides around A-to-I RNA editing sites. (b) The genome distribution of RNA editing sites. (c) RNA editing enriched in repeat elements and harbored double-strand RNA. (d) The number of the identified A-to-I editing sites (left). The distribution of RNA editing levels in 52 tissues across three stages (e) Sample (n = 400) clustering using principal component analysis based on common RNA editing level (Sample>100). (f) Correlations between the expression level of ADAR family genes and the edited reads (corrected by sequence depth, the top three figures). Relationships between the expression level of ADAR family genes and overall editing level (the bottom three figures). (g) The enrichment of genetic variance was evaluated for all RNA editing sites (3,093,058), as well as for those conserved between human and cattle (17,955), swine and cattle (1,034), and among all three species (87). Enrichment was defined as the ratio of heritability captured by RNA editing sites to the mean heritability captured by 20 randomly shuffled sets of editing site positions.

The cerebellum, cerebrum, blood vessels, and marrow displayed the highest number of A-to-I editing sites (Figure 5d), while muscle tissue and cartilage exhibited the lowest number. In most tissues, the RNA editing numbers were limited in the newborn stage, indicating RNA editing in newborn was less active than in adults. For example, plenty of low levels of RNA editing were detected in adipose tissue of newborn stage, while the levels turned to high in the young and adult stages (Figure 5d). We conducted PCA on 1,690 high-quality editing sites that were presented in at least 100 samples. The brain tissues were separated, and other tissues, like the immune, blood vessels, liver, and muscle could also be distinct (Figure 4e and Figure S25). It had been reported that ADAR enzymes catalyzed A-to-I editing in mammals [25]. We observed dynamic expression of *ADAR* and *ADARB1* across 52 tissues, whereas *ADARB2* was predominantly expressed in the brains and newborn adrenal glands (Figure S26). Using standardized regression analysis, we determined that *ADARB1* and *ADAR* expressions contributed 33% and 19% of the variations in edited reads ratio (Figure 5f). For the global RNA editing level, *ADARB1* could explain 15% variations, while *ADAR* was not significant, suggesting that *ADARB1* expression might play a crucial role in determining global editing levels. *ADARB2*, which was exclusively present in brain tissue, showed a weaker association with RNA editing events across tissues. We detected that *ADARB1* and *ADARB2* in the hypothalamus were significantly up-regulated at the young stage, while downregulated at the adult (Figure S27). For the adrenal gland, all the three ADAR genes were significantly downregulated after birth. Interestingly, we found that nine RNA editing sites located downstream of *TMED10* (Transmembrane P24 Trafficking Protein 10) and *CCNYL1* (Cyclin Y Like 1) genes shared by all 400 samples (Table S6).

We further examined the relationship between evolutionary conservation of RNA editing sites (Table S7) and their enrichment for trait-associated genetic variance. For most traits, RNA editing sites conserved across three species (human, swine, and cattle) exhibited higher heritability enrichment than those conserved in only two species (human–cattle or swine–cattle). Both categories showed greater enrichment compared to all detected RNA editing sites. Notably, this trend remained consistent even when the enrichment was negative, suggesting a general pattern where higher evolutionary conservation corresponded to greater functional relevance in relation to complex traits (Figure 5g).

### Integrate transcriptomic BodyMap with cattle agronomic traits

We estimate that 43.1% ∼ 89.6% of phenotypic variation for beef carcass and quality traits was tagged by autosomal SNPs (Table S8). To assess the contribution of genetic variance by novel lincRNA to 23 cattle agronomic traits, we captured proximal SNPs within them, including their flankingLJ5kb regions. The novel lincRNAs explained more variance than expected by chance for most traits (paired t-test *P* = 0.042), especially for rib-eye area and carcass length (Figure S28). Using weighted gene co-expression network analysis (WGCNA), we identified that 55 co-expression modules involving 17,301 PCGs and 12,331 lncRNAs expressed in at least 20% of the samples (Figure S29). Based on module membership (|k_ME_| ≥ 0.9), a total of 2167 hub genes were identified, including 1651 PCGs and 516 lncRNAs (Table S9). Then we performed genetic variance analysis using SNPs located within hub genes and compared them with SNPs from all genes expressed in at least 20% of the samples. For most traits, hub genes captured the higher or more depleted proportion of genetic variance (Figure 6a). In addition, 81,415 variants located in the editing clusters explained more variance than expected by chance for 12 traits, especially for fat coverage rate and chunk roll weight, which implied the causative variants might be enriched in these edited regions (Table S8).

**Figure 6.**
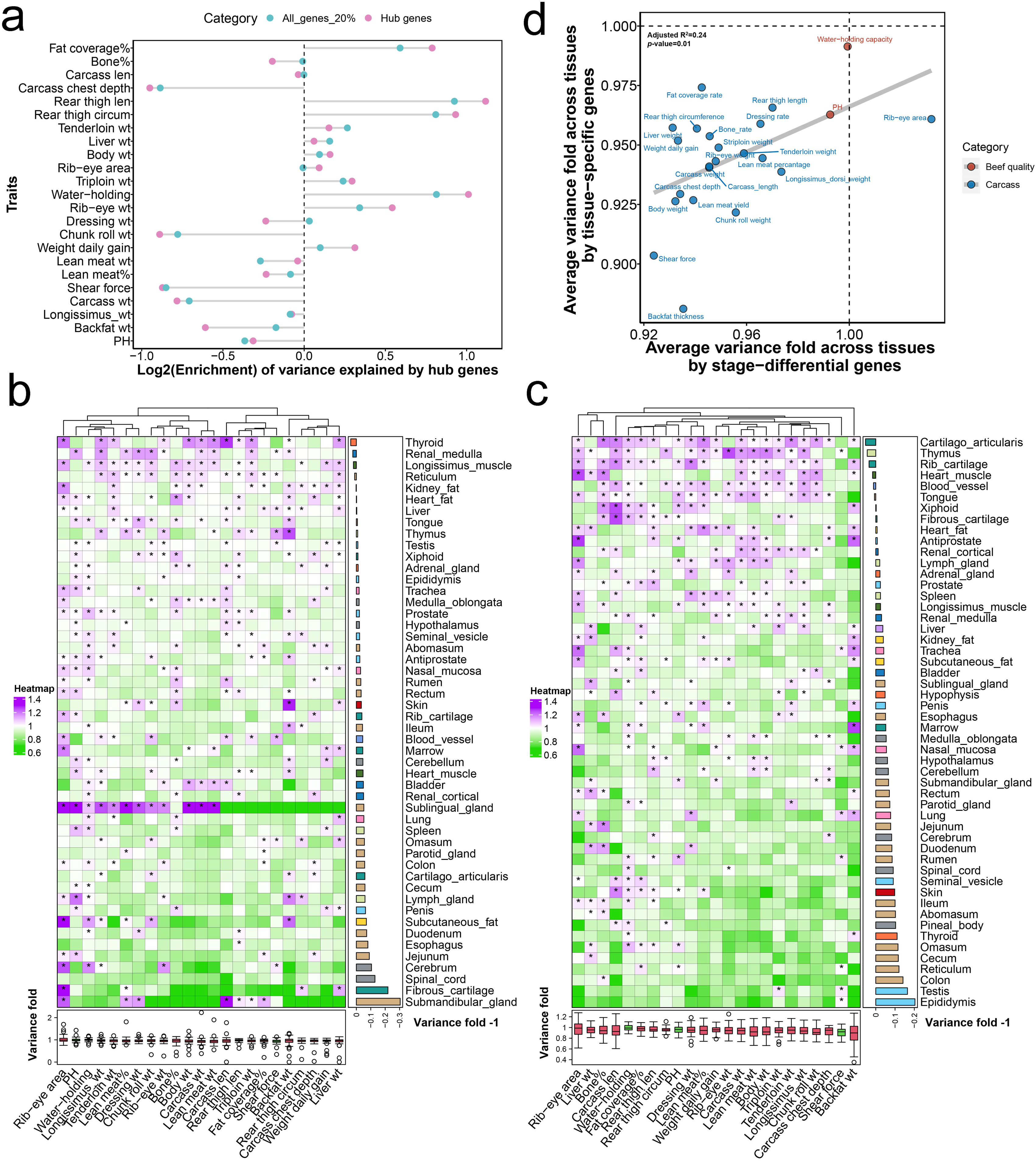
Genetic variance explained by development-associated and tissue-specific genes in 23 beef production traits. (a) The enrichment of genetic variance was evaluated for PCGs and lncRNAs expressed in at least 20% of the samples, as well as hub genes. Enrichment was defined as the ratio of heritability captured by genes to the mean heritability captured by 20 randomly shuffled sets of gene positions. (b) The variance enrichment was analyzed through development-associated genes across three comparison groups for each tissue (row), in relation to 23 beef production traits (column). Enrichment folds were determined as the ratio between the SNP heritability of development-associated genes and the average derived from 20 permutations with an equivalent number of SNPs. Enrichment ratios exceeding 1.0 are indicated with star symbols. The bottom boxplot displays the distribution of enrichment ratios across tissues for each trait, while the right barplot illustrates the average enrichment ratio across traits for each tissue, adjusted by subtracting 1. (c) The enrichment of genetic variance for each tissue (row) was analyzed through tissue-specific genes, in relation to 23 beef production traits (column). Enrichment folds were determined as the ratio between the SNP heritability of development-associated genes and the average derived from 20 permutations with an equivalent number of SNPs. (d) Comparing the enrichment fold of genetic variance for each trait between development-associated genes and tissue-specific genes across tissues.

DEGs identified between newborn and young stages in the cerebellum, longissimus muscle, testis, cartilago articularis, reticulum, renal medulla, epididymis, and tongue were found to explain a greater average variance for all traits compared to randomly selected genes (Figure S30). Similarly, the DEGs between newborn and adult stages in the renal medulla, thymus, thyroid, medulla oblongata, and heart fat were also able to explain more average variance for all traits compared to randomly selected genes (Figure S31). Additionally, the DEGs between young and adult stages in the omasum, renal medulla, heart muscle, spleen, liver, blood vessel, adrenal gland, esophagus, and cerebellum were found to explain more average variance for all traits compared to randomly selected genes (Figure S32). When we merged all DEGs across stages together, we found DEGs of the thyroid, renal medulla, longissimus muscle, and reticulum captured more variance of traits (Figure 6b).

We also investigated whether tissue-specific genes could capture a greater amount of genetic variance for economic traits. The tissue-specific genes were defined based on the top 5% rank of t-statistics. We found that specific genes in various tissues, such as cartilago articularis, thymus, rib cartilage, heart muscle, blood vessel, and tongue, explained more average variance for all traits compared to randomly selected genes (Figure 6c). Xiphoid and fibrous cartilage captured more variance for carcass length. In the newborn stage, specific genes in rib cartilage, longissimus muscle, renal cortical, and tongue exhibited a higher average variance for all traits than randomly selected genes (Figure S33). Additionally, heart fat and kidney fat were the most enriched tissues for liver weight. In the young stage, specific genes in rib cartilage, heart muscle, esophagus, and cartilago articularis displayed a higher average variance for all traits compared to randomly selected genes (Figure S34). In the adult stage, specific genes in cartilago articularis, blood vessels, subcutaneous fat, kidney fat, and thymus demonstrated a higher average variance for all traits compared to randomly selected genes (Figure S35). Similarly, the subcutaneous fat, heart fat, marrow, and penis were the most enriched tissues for backfat, and omasum was associated with pH. We observed a weak correlation between development-associated and tissue-specific genes across tissues (Figure S36). However, we found a significant correlation between stage-different genes and tissue-specific genes for genetic variance across traits (Figure 6d).

### Genomic prediction of beef production traits using prior information

These DEGs in each tissue with the ratio of genetic variance divided by that calculated by permutation test larger than 1.01 were defined as prior biological information. The tissues used as prior biological information in each stage comparison were shown in Table S10. We divided the original 675K genotype panel into two groups based on whether the SNPs were in prior biological information. We fitted these two SNP sets simultaneously with BayesRC models to conduct two-component genomic predictions. When we treated the DEGs of each comparison group as separate prior biological information, the predictive reliability for beef carcass and quality traits was higher compared to the BayesR model. Specifically, we observed increases of 0.19%, 0.17%, and 0.33% for Young vs. Newborn, Adult vs. Newborn, and Adult vs. Young (Table 1), respectively. When incorporating prior biological information, a more noticeable increase (> 1% increment) in reliability was observed for WHC and rear thigh length for Young vs. Newborn, Rear thigh circumference and RTL, and carcass chest depth (CCD) for Adult vs. Newborn, as well as lean meat percentage, chunk roll weight, WHC, RTL and CCD for Adult vs. Young. Compared to the BayesR model, prior biological information merged from all three comparison groups resulted in higher predictive reliability with BayesRC for beef carcass and quality traits (average increase of 0.42%, Table 1 and Figure 7a). For the detailed traits, we found the predictive reliabilities of most traits (17 of 23) were increased. The reliability increment obtained using prior biological information was more noticeable for water-holding capacity (WHC) and rear thigh length (RTL) with 4.6% and 1.4% increments.

**Figure 7.**
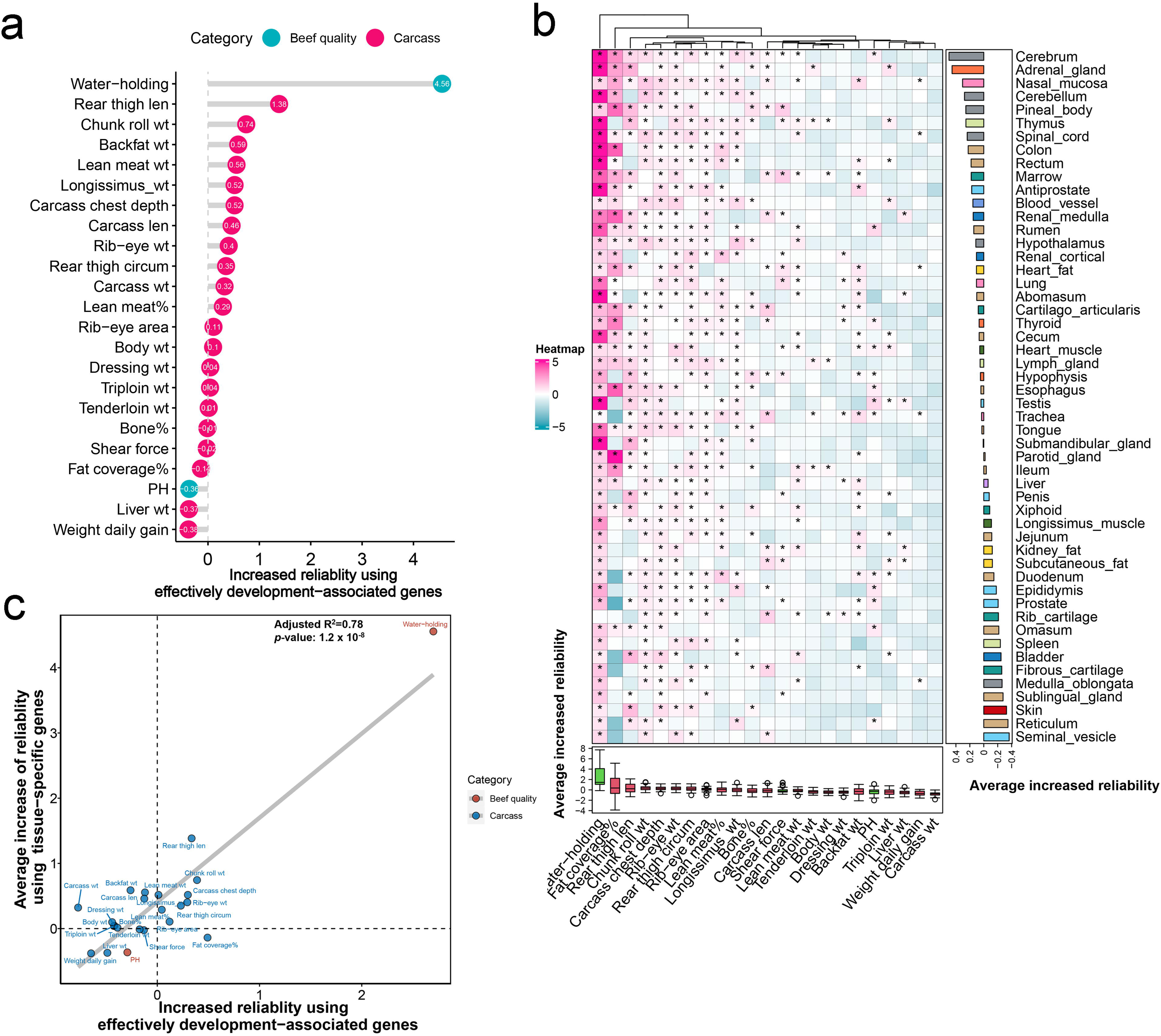
Development-associated and tissue-specific genes improve genomic predictions of 23 beef production traits. (a) The increased reliabilities of genomic predictions in beef production traits using priors of effectively development-associated genes. (b) The increased reliabilities of genomic predictions in beef production traits using priors of tissue-specific genes for each tissue. Increased reliabilities exceeding 0 are indicated with star symbols. The bottom boxplot displays the distribution of increased reliabilities across tissues for each trait, while the right barplot illustrates the average increased reliability across traits for each tissue. (c) Comparing the increased reliabilities of genomic predictions using prior information for 23 beef production traits between effectively development-associated genes and tissue-specific genes.

**Table 1.** The genomic predictive reliability of 23 traits using prior information of development-associated genes.

| Traits | Non-Prior | Adult vs. Newborn priors | Young vs. Newborn priors | Adult vs. Young priors | All Priors |
| --- | --- | --- | --- | --- | --- |
| PH | 37.93% | 37.56% | 36.76% | 37.88% | 37.56% |
| Backfat thickness | 31.88% | 32.19% | 31.45% | 32.12% | 32.46% |
| Longissimus dorsi weight | 44.26% | 44.04% | 44.09% | 45.07% | 44.77% |
| Carcass weight | 40.56% | 40.54% | 39.66% | 40.88% | 40.88% |
| Shear force | 36.63% | 36.40% | 36.86% | 36.02% | 36.61% |
| Lean meat percentage | 37.05% | 36.92% | 36.89% | 38.13% | 37.34% |
| Lean meat yield | 39.74% | 40.10% | 40.21% | 39.87% | 40.30% |
| Weight daily gain | 35.35% | 35.34% | 34.88% | 35.04% | 34.97% |
| Chunk roll weight | 41.78% | 42.15% | 42.29% | 42.81% | 42.52% |
| Dressing rate | 39.99% | 39.96% | 40.17% | 39.96% | 40.03% |
| Rib-eye weight | 43.03% | 43.64% | 43.56% | 43.14% | 43.43% |
| Water-holding capacity | 62.28% | 63.19% | 63.48% | 64.06% | 66.83% |
| Striploin weight | 43.20% | 43.02% | 43.28% | 43.13% | 43.24% |
| Rib-eye area | 44.38% | 44.49% | 44.46% | 44.47% | 44.48% |
| Body weight | 38.88% | 38.29% | 39.01% | 38.77% | 38.98% |
| Liver weight | 35.49% | 35.58% | 35.41% | 35.67% | 35.12% |
| Tenderloin weight | 41.87% | 41.94% | 41.80% | 41.58% | 41.88% |
| Rear thigh circumference | 51.95% | 53.02% | 52.11% | 51.68% | 52.30% |
| Rear thigh length | 56.00% | 57.87% | 57.68% | 57.04% | 57.39% |
| Carcass chest depth | 42.88% | 43.93% | 43.44% | 44.32% | 43.40% |
| Carcass length | 30.45% | 30.64% | 30.78% | 30.93% | 30.91% |
| Bone rate | 33.69% | 33.21% | 34.04% | 33.65% | 33.68% |
| Fat coverage rate | 52.05% | 51.58% | 52.82% | 52.74% | 51.91% |
| Average | 41.80% | 41.98% | 41.96% | 42.13% | 42.22% |

**Note:** The increased reliabilities were marked with red color.

Using tissue-specific genes as prior biological information, we observed that tissue-specific genes of most brain and digestive tissues helped improve the predictive reliability of most carcass and quality traits (Figure 7b). Specific genes in the cerebellum demonstrated higher predictive reliability, with an average of 0.49% across all traits, compared to the model without prior information. The reliability increment obtained using tissue-specific genes was more noticeable in WHC and fat coverage rate, chunk roll weight, rear thigh length and circumference, carcass chest depth, rib-eye weight, and area. Specifically, the predictive reliability of WHC was improved by an average of 2.7% using different tissue-specific genes. When comparing the increased reliability in different traits using development-associated genes and tissue-specific genes, we found a strong correlation in the increased predictive reliability across traits between development-related genes and tissue-specific genes (Figure 7c).

### Cattle Bodymap Transcriptome (CBT) Database

To visualize and explore gene expression, lncRNAs, RNA editing and splicing across multiple tissues and stages based on cattle bodymap of transcriptome data, we developed the CBT Database (http://cattlegenomics.online/cattle_bodymap/), an interactive web portal featuring a user-friendly interface constructed using the R/Shiny framework (Figure 8a). It offers uniform and flexible interfaces for manipulating query parameters and provides six menus (Home, Gene expression, LncRNA, RNA editing, Splicing and About). The “Home” menu introduces the datasets and function of CBT Database, and the “About” menu displays information about technologies and the installation procedures of CBT Database. The remaining four menus show versatile retrieval and visualization functions. These functional menus provide a search box that users can utilize a specific genomic region, gene name or Ensembl ID based on their defined tissues and stages (Figure 8b). The graphics and table produced by these functions are available for download (Figure 8c and 8d).

**Figure 8.**
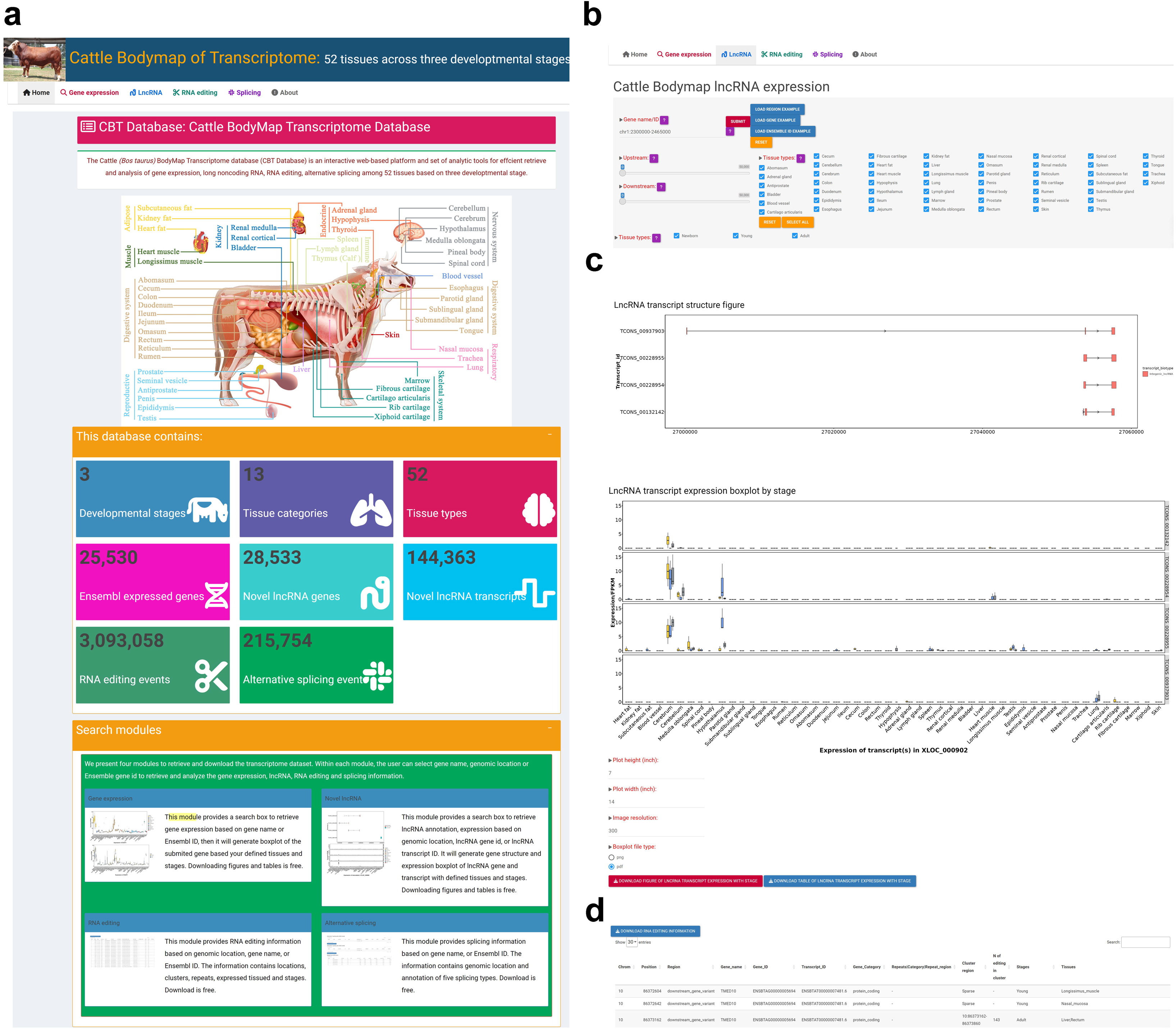
The graphical interface of the Cattle BodyMap of Transcriptome (CBT) Database. (a) The home page of CBT Database. A total of six menus are implemented in the database. (b) The search function of CBT Database. Users can search information using a specific genomic region, gene name or Ensembl ID with their defined tissues and stages. (c) An example of figure generated based on long non-coding RNAs (lncRNA) searching. (d) An example of table generated based on RNA editing searching.

For instance, a researcher interested in muscle development can query the expression profile of *MYF5*, a key myogenic regulatory gene, across multiple tissues and developmental stages. The CBT Database will visualize its spatiotemporal expression patterns, which helps infer its regulatory roles during myogenesis. Similarly, researchers interested in exploring regulatory mechanisms can use the CBT Database to retrieve and analyze lncRNAs in specific genomic regions. For example, querying the region chr1:2300000–2465000 will return all annotated lncRNAs within this interval, along with their transcript structures and expression profiles across different tissues and developmental stages. Such information can help prioritize candidate lncRNAs potentially associated with economically important traits, paving the way for further functional studies or marker development in genomic selection programs. Moreover, by exploring tissue-specific splicing or RNA editing events in reproductive organs, the database may aid in pinpointing regulatory elements associated with fertility traits. The CBT Database is compatible, extendable, portable, and easy to be established in different operating systems (Linux, macOS, and Windows) with an installation guide at https://github.com/WentaoCai/CBTDatabase.

## Discussion

The emergence of RNA-seq has revolutionized our comprehension of gene expression, AS, and RNA editing, shedding light on their involvement in normal physiological processes and their potential impact on economically complex traits in cattle. Nevertheless, despite the increasing number of transcriptome studies, there has been a lack of a comprehensive and centralized resource that consolidates crucial and easily accessible information on the regulation and functional associations of AS and RNA editing events, hindering further explorations in the field. In this study, we have presented and described such a resource.

Our study significantly enhances the annotation of transcripts in the cattle genome and provides a comprehensive landscape of transcription, splicing, and RNA editing sites across tissues with varying developmental stages. Cattle tissue transcription was generally dominated by the expression of a relatively small number of genes, which was the same with humans [16] and pigs [5]. The transcription was mitochondrial origin in the brain, heart, and fat tissues, while the kidney exhibited the highest enrichment of mitochondrial genes in humans [16]. While numerous genes displayed organ-specific differential expression throughout developmental stages, less than half of the genes were expressed in all tissues and developmental stages. These frequently expressed PCGs seemed to primarily participate in fundamentally biological processes, which implied they were essential to cell life. The cellular composition of the testis underwent the substantial change during sexual maturation, with spermatogenic cells becoming predominant [26, 27].

AS of genes plays a critical role in mechanism in organ development during the intricate process of forming complex organisms [13]. Aberrant organ-specific expression of isoforms may contribute to human diseases [18]. We cataloged and assessed the expression of 215,754 alternatively spliced transcripts in cattle. Like other mammalians [28, 29], SE was the most prevalent type of AS event in cattle. Nasal epithelial cells are a reliable source to study splicing variants [30]. We detected that nasal mucosa had more AS events than other tissues in cattle. These genes with PanAS were involved in essential activities of cells, such as metabolic pathways. While genes with switchlike AS were associated with signaling pathways, such as Rap1 signaling pathway and phosphatidylinositol signaling system. Sample from similar tissues clustered together based on PSI, indicating distinct splicing programs in each tissue category. Different organs could be distinct by development stage, suggesting strong commonalities for organ splicome at each stage. ELAVL family, shown to regulate AS in neurons [31], is involved in brain development and disease [32]. YBX1 is essential for mRNA stability during muscle-fiber formation [33]. Here, we identified that *YBX1* was specifically highly expressed in muscle tissues. AS events were more active in male reproductive, respiratory, skin, and fat tissues during development, indicating AS was essential for organ development. We acknowledged the limitations of using short-read RNA sequencing for analyzing alternative splicing events. Specifically, its shorter read lengths might hinder the accurate distinction of closely related splice variants and the identification of low-abundance transcripts, which could impact the comprehensive understanding of the splicing landscape in cattle.

A-to-I RNA editing sites frequently occur in clusters, although some sites, particularly those within the coding sequence, are often isolated. Therefore, we employed two complementary methods. This study had identified more than 3 million RNA editing sites, significantly broadening the RNA editing dataset and enabling further exploration of RNA editing function. Similar to other mammals [34], A-to-I editing was the predominant type of RNA editing in cattle. Almost all these A-to-I editing sites were within clusters, indicating the hyper-edited mechanism commonly existed [35]. While the majority of identified A-to-I RNA editing sites resided in non-coding regions, they might exert crucial regulatory functions by modulating RNA secondary structures [36], splicing efficiency [37], miRNA interactions [38], lncRNA activity [39], and transcriptional regulation through promoter/enhancer-associated RNAs [40]. In mammals, most A-to-I editing sites are located in repetitive elements, especially SINEs, such as Alu in humans [41], PRE in pigs [42], B1 in mice [43], and Bov in cattle (this study). The primary role of the broadly expressed ADAR in vertebrates is to edit endogenous long dsRNAs, thereby preventing an innate immune response triggered by these endogenous transcripts [14]. The RNA editing cluster regions identified in this study harbored plenty of dsRNA structures. Unlike *ADAR*, which predominated in humans [44], our research confirmed that *ADARB1* was the primary contributor to the A-to-I changes across various tissues. In addition to the brain, RNA editing was also quite active in the bone marrow and blood vessels. Previous studies also detected RNA editing was abundant in arteries and was elevated in several critical cardiovascular conditions [45]. The RNA editing numbers were limited in most tissue of the newborn stage, indicating RNA editing in infancy was less active. Only *TMED10* and *CCNYL1* genes were edited in their downstream regions in all samples. *TMED10* is involved in intracellular protein transport, which is a candidate gene for hip width or rump width in cattle [46]. *CCNYL1* plays a crucial role as a regulator of cell cycle transitions and positive regulation of cyclin-dependent protein serine/threonine kinase activity [47]. *CCNYL1*, which is located within binding regions of miRNAs, can be edited in prostate cancer [48]. Although RNA editing sites contributed modestly to trait variance, those conserved across multiple species tended to show higher enrichment, suggesting evolutionary conservation as a useful indicator of functional importance. A-to-I RNA editing affects skeletal muscle differentiation and function by altering selenoprotein expression [49]. To our knowledge, no previous publication had investigated the associations between tissue-specific or development-related gene and beef carcass traits through an integrative analysis of genomic and transcriptomic data. Hub genes identified from co-expression modules captured more or less genetic variance than all expressed genes in most traits, indicating their potential functional relevance or selective constraints. The RNA editing clusters explained more variance in chunk roll weight and weight daily gain, suggesting RNA editing might play a role in skeletal muscle development. LncRNAs might be important regulators for beef production and quality [50, 51] and related to skeletal muscle development [52]and muscle atrophy [53]. In this study, the novel lincRNAs captured more genetic variance than randomly genomic regions for most beef production traits, implying these novel lincRNAs held potential for applications in the treatment of human muscle diseases and the improvement of production traits in beef cattle.

We identified relevant tissues for 23 beef production traits of beef cattle and further applied them in genomic prediction. Of interest, we found that DEGs of the cerebellum, longissimus muscle, testis, cartilago articularis, and reticulum between newborn and young stage were associated with body composition weight traits. *MFAP5* and *FRZB* were some of the most significant genes in the cerebellum and longissimus muscle, respectively. *MFAP5* is positively correlated with body mass index [54] and obesity [55]. *FRZB* is a famous gene involved in muscle atrophy and dystrophy [56]. The omasum DEGs between young and adult stages were associated with most beef production and quality traits. *LOC100848132*, *CPA1*, and *PGA5* of omasum are involved in protein digestion and absorption [57]. The tissue-specific genes for cartilago articularis, thymus, and rib cartilage were associated with most beef production traits. *ZNF648* (Zinc Finger Protein 648) is involved in DNA-binding transcription factor activity and related to fat body mass [58]. *ZNF648* was specifically highly expressed in both cartilago articularis and rib cartilage, implying its specific function in the skeletal system.

The genomic variants of beef production traits seem to be enriched in certain genome regions. This means that not all markers contribute equally to the variability of these traits, which challenges the assumption made by the GBLUP approach [59]. One possible solution to address this issue is to incorporate prior genomic information into the model in a more sophisticated way, such as using Bayesian methods [60] or weighted marker approaches [61]. The key of these methods is how to identify and utilize effective prior information. Our study introduced two strategies for identifying effectively prior information based on transcriptome data that improved the reliability of genomic prediction. When we defined the prior information using a high variance of development-related genes, most traits achieved higher reliabilities of genomic predictions. When incorporating tissue-specific genes into the Bayesian prediction models, we thus observed an increase in predictive reliability compared to the non-prior model. The observed inconsistencies in predictive improvements across traits were likely due to significant variations in the genetic variance explained by tissue-specific and developmental stage-specific genes, as shown by genetic variance analyses. Additionally, the diverse genetic architectures and underlying regulatory mechanisms of different traits influenced their responsiveness to tissue-specific genetic effects. In some cases, predictive improvement might be limited due to mismatches between causal variants and context-specific gene expression, lower heritability, or the involvement of cross-tissue regulatory networks [62, 63].

We developed a novel database designed to store, retrieve, and analyze transcriptome data using exclusively R code, a widely-adopted programming language renowned in statistical and biological research. This method of website development was particularly beneficial for R users, obviating the need for expertise in SQL or other server-side programming languages. The source code for the CBT Database had been publicly accessible, enabling its straightforward adaptation for the creation of genomic databases in other organisms. Distinguishing CBT from other cattle genomic databases, such as the Bovine Genome Database (BGD) [64] and Bovine Genome Variation Database (BGVD) [65], CBT was pioneering in its integration of lncRNAs, RNA editing, and splicing information. Additionally, the database incorporated expression data from 52 different tissues at three distinct developmental stages, providing a unique resource for cattle genomic research. By offering swift and efficient access to extensive transcriptome data, the CBT Database stands to greatly facilitate future investigations in functional and comparative genomics, as well as other related disciplines, without the requirement for programming abilities.

This study provided a comprehensive transcriptomic BodyMap across multiple tissues and developmental stages in cattle, while it had several limitations. Although we identified numerous lncRNAs, AS events, and RNA editing sites across tissues and stages, the current study did not focus on their trait-specific functional characterization or molecular validation. In the future, this would be an important direction.

## Conclusions

Our study established the most comprehensive cattle transcriptomic BodyMap, covering 52 organs across 3 developmental stages. We generated large-scale profiles for gene expression, AS, and RNA editing across multiple tissues in cattle, providing valuable information for enhancing genome annotation and understanding biological functions. Through the integration of these profiles with a large population of genotypes and phenotypes, we uncovered a comprehensive genetic relationship between tissues and 23 economical traits in beef cattle, offering novel perspectives into the molecular mechanisms underlying these economically important traits. This information can be leveraged for developing more efficient breeding strategies and enhance the selection of superior animals for improved beef production. The CBT Database, an interactive web portal boasting a user-friendly interface, hosts the transcriptome data of the cattle BodyMap. This resource is poised to be of significant value for future functional genomics research on cattle.

## Methods

### Sample collection and processing

Nine half-sibling male Simmental beef cattle from Shayang (Hubei province of China, Hanjiang Cattle Development Co., Ltd.) were selected for sampling. Three calves were slaughtered at birth. Three calves were slaughtered after weaning at 6 months and the remaining three calves were fattened under the identical feeding and management conditions until they reached 24 months of age. After slaughtering, we collected a total of 400 tissue samples from 52 different tissue types, comprising 124 samples from newborns, 145 samples from young individuals, and 131 samples from adults. All samples were promptly frozen in liquid nitrogen for total RNA extraction. The nine blood samples were collected prior to transportation to the slaughter room. All experiments and procedures were conducted in compliance with the regulations outlined by the Animal Care and Ethics Committee for Animal Experiments at the Institute of Animal Science, Chinese Academy of Agricultural Sciences (Permit Number: IAS2021-43).

### RNA sequencing, expression quantification, and quality control

RNA samples that met the quality control criteria were utilized to create 400 transcriptome libraries employing a standard non-strand specific protocol with poly-A selection of mRNA. These libraries were sequenced on the Illumina HiSeq-PE150 platform. The 400 RNA-seq data supporting this study’s findings is available from the National Genomics Data Center (NGDC) database with accession numbers listed in Table S1. Quality trimming and adaptor removal of reads were performed using Cutadapt v2.8 [66] and Trimmomatic v0.39 [67]. The clean data were aligned with Hisat2 v2.1.0 to the ensemble genome release version ARS UCD1.2 [68]. Ensembl version 105 served as the reference for transcript model alignment and gene / isoform quantification, encompassing a total of 21,861 genes including 20,110 PCGs, 1480 lncRNAs, 393 pseudogenes, and 3873 other genes (except the above three types).

Gene expression were quantified in FPKM using Stringtie -e -B [69]. The expression threshold for a gene to be considered as expressed was set at FPKM >= 0.1 in at least one sample. To explore gene expression similarity between tissues and stages, we conducted principal component analysis (PCA) and hierarchical clustering (HC) using a log2-transformed scale of FPKM. The distance between samples was calculated as 1 – Spearman correlation.

### Novel lncRNA identification

The novel transcripts were assembled by StringTie (v1.3.5) [69] and Scripture (beta2) [70]. Transcripts that were present in at least two samples or supported by two assembly software programs were selected. Additionally, we preserved novel intergenic and intronic transcripts by focusing on class codes ‘u’ and ‘i’ as indicated in the gffcompare annotation (v0.11.6) [71]. Transcripts with length ≥ 200 nt, exon ≥ 2, and lengths of maximum open reading frame ≥ 120 amino acids were obtained for protein-coding potential prediction. Transcripts lacking protein-coding potential (CPC score < 0, PLEK score < 0, and CNCI score < 0) were kept. To eliminate transcripts containing known protein domains, we translated transcripts into amino acid sequences and conducted a search against the Pfam database (Pfam 30.0) using HMMER [72]. Transcripts demonstrating significant Pfam hits were disregarded.

### Differential, preferential, exclusive, and specific gene expression analysis

Differential expression analysis was conducted using DEseq2 with FDR < 0.05 as the cut-off for statistical significance [73]. We used as input read counts obtained by FeatureCount [74]. All pairwise combinations between all tissues or stages were tested. We used |log_2_fold change| >= 4 to call tissue preferential gene expression after comparing the samples from a given tissue to those samples that did not belong to the tissue [16]. To assess tissue exclusivity, we calculated the phi correlation coefficient on the contingency table generated by dividing the samples based on two conditions. The first condition involved selecting samples based on whether they were expressed (FPKM ≥ 0.1) or not expressed (FPKM < 0.1). The second condition involved selecting samples coming from either the tissue being analyzed or all other tissues. We used the function “phi” from the R package psych to calculate the phi-correlation coefficient (http://CRAN.Rproject.org/package=psych). We defined those genes with phi >= 0.95 or =< -0.95 as tissue-exclusive genes.

The *t*-statistic of specific gene expression within a given tissue was calculated from 400 samples across 52 tissues with 13 tissue categories. Samples from the same tissue category were excluded during analysis while considering known covariates. For example, when computing the *t*-statistic of specific gene expression in kidney fat, expression levels in kidney fat samples were compared against expression levels in all other non-adipose tissue samples, excluding subcutaneous fat and heart fat. The *t*-statistics were determined using a general linear model:

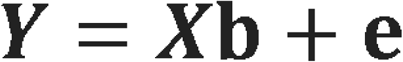

where ***Y*** was log2-transformed gene expression and normalized using Z-score normalization within each tissue. ***X*** was a design matrix, with each row corresponding to a sample. The first column of ***X*** had a ‘1’ for each kidney fat sample and a ‘–1’ for each non-adipose tissue sample. **b** is the corresponding tissue effect. **e** was the residual effect. We calculated *t*-statistics using ordinary least-squares according to Finucane’s formula [75]. We constructed integrative transcriptome co-expression networks using WGCNA based on PCGs and lncRNAs expressed in at least 20% of the samples. Co-expression modules were detected, and hub genes or lncRNAs were defined as those with high module membership (|k_ME_| ≥ 0.9), where k_ME_ represented the correlation between each gene’s expression and the corresponding module eigengene.

### AS identification

We identified AS and estimated exon splicing levels (percent spliced in, or PSI) using rMATs turbo v4.2.0 [76]. The AS events with 0.1 < PSI < 0.9 in >80% of the samples were defined as PanAS, while AS events with PSI > 0.9 or PSI <0.1 in >80% of the samples were defined as SwitchAS. Subsequently, we employed rMATS turbo v4.2.0 with a threshold set at FDRLJ<LJ5% and |deltaPSI|LJ≥5% to detect differential AS events between tissues or developmental stages.

### Identifying isolated RNA editing using whole genome sequencing (WGS) and RNA-seq

Genomic DNA was extracted from 9 blood samples utilizing the Qiagen DNeasy Kit. Paired-end libraries with an average 500 bp insert size were generated using standard procedures. All libraries were sequenced on the MGISEQ-T7 platform in PE150 mode (BGI, Shenzhen, China)) to an average raw read sequence coverage of 30 ×. The raw reads were quality-controlled by Fastp (V0.21.0, parameters: -n 10 -q 20 -u 40) [77]. To detect isolated editing events, we identified putative RNA editing sites by the combination of RNA-seq and WGS. We used BWA mem (v0.7.17) [78] and Hisat2 (v2.1.0) to align WGS and RNA-seq to the ARS-UCD 1.2 genome. Alignments were enhanced using Picard [79] and GATK tools [80]. To eliminate heterozygous SNPs in the genome and precisely identify RNA-editing sites, we compared RNA and DNA BAM files using REDItools [81]. The isolated editing events were defined when sites with DNA coverage ≥ 10, RNA coverage ≥ 5, and edited reads ≥ 3.

### Identification of hyper RNA editing sites

Adenosine (A) to inosine (I) editing sites are often clustered together, but certain sites, especially those located within the coding sequence, are typically solitary. [82]. Due to the large number of clustered editing sites causing heavily (hyper) edited reads, we prioritized the analysis of the unmapped reads. We substituted all adenines with guanines (Gs) in both the unmapped reads and the reference genome, followed by realigning the modified RNA reads to the transformed reference genome using BWA (v0.7.17) [78]. Subsequently, we extracted and restored the mapped reads to their initial sequences, categorizing them as candidate hyper-edited reads. To improve the accuracy of identifying hyper-editing reads, we set the threshold for the number of A- to-G mismatches at 5% of the read length and 80% of the total mismatches. Furthermore, we applied additional filters to the reads, requiring the average Phred quality score > 25, ambivalent nucleotides (N) < 10%, simple repeats < 10, and successive single nucleotides < 20. To eliminate heterozygous SNP, we compared hyper RNA editing sites to DNA BAM files using REDItools [81]. We followed the same methodology to detect all other possible types of mismatches, such as A-to-C, G-to-A, and others. The RNA-seq reads could originate from either the sense or antisense strand (in the case of a non-stranded library). As a result, although there are 12 potential single-nucleotide mismatches, our analysis specifically focused on six categories of editing events: A-to-G, G-to-A, A-to-T, A-to-C, G-to-C, and C-to-A. Each category accounted for both the forward and complementary mismatches.

### Defining editing level, clusters, and dsRNA structure

The hyper and isolated RNA editing sites were combined to calculate the editing level of each site, which was quantified as the number of Gs divided by the total of As + Gs at that specific site [83]. The overall editing level of each sample was computed by dividing the total number of Gs by the combined total of As + Gs across all editing sites. We defined the clusters as the distance between adjacent RNA editing sites being less than 100bp, where the number of editing clusters was linear with the distance (Figure S24b). To assess whether editing clusters harbored dsRNA structures, we aligned the DNA sequence of each editing cluster to its spanning 2 Kb region using bl2seq with parameters of -F F -W 7 -r 2 -D 1 [84]. Alignment with 90% identity along 80% of the editing cluster length was considered as a dsRNA structure. To assess the conservation of RNA editing sites across species, we analyzed 15.6 million human sites (from REDIportal [85]) and 727,775 swine sites from published studies [42, 86]. Using the UCSC LiftOver tool, we converted the genomic coordinates of human and swine RNA editing sites to their corresponding positions in the cattle genome.

### Variance component analysis

The phenotype of 23 traits was collected from 1476 individuals over the past decade. The summary of phenotype records can be found in Table S10. All individuals were genotyped using with the Illumina Bovine 770K Bead chip. Genotype data included SNP and insertion-deletion (InDel) was imputated from the 1000 Bull Genomes Project, described in detail previously [87]. After quality control, 8,849,824 autosome variants were available for variance component analysis. We used restricted maximum likelihood (REML) to calculate the proportion of variance explained by each functional class of SNP with the following model:

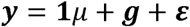

where the vector **y** represents phenotypic data that has been adjusted for year, sex, age, and the first two principal components (PCs) of the genotype. These PCs were computed using PLINK [88] and further normalized through rank-transformation using the transform function of GenABEL [89]. ***1****μ* is a vector of trait means. ***g*** is a vector of total additive genetic effects 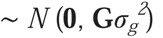, where **G** represents the genomic relationship matrix (GRM) and 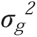 denotes additive genetic variance. *ε* denotes random residual errors 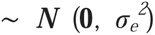, where 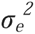 is the error variance. 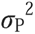 is the phenotypic variance. The proportion of variance explained was obtained using the GRM by a REML analysis in GCTA [90]. Then heritability 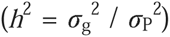 was the proportion of phenotypic variance explained by all variants together.

To determine whether SNPs in novel lincRNAs, RNA editing clusters, or DEGs explained more variance than the variance explained by the same number of randomly selected variants, we performed variance component analysis using a shifted permutation test. To perform the shifted permutation test, we first converted the genomic position of variants in the functional class into a continuous position. For a variant with genomic position B in chromosome N, its continuous position should be 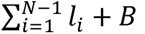. Then the observed variants set was shifted to the new continuous (*p*_1_, *p_2_*, *p*_3_, …, *p*_4_) based on random value R within 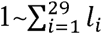 using the following formula:

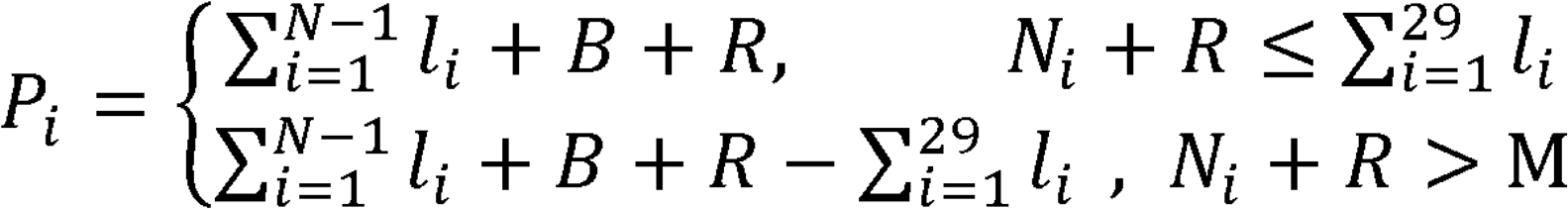

The new positions were recovered into genomic positions based on autosome length. This shifted permutation test using Perl scripts is available in an online code repository (https://github.com/WentaoCai/Permutation). We repeated this permutation 20 times to obtain stable results. The enrichment fold was equal to the heritability of functional class divided by the average of 20 permutations

### Genomic prediction

We randomly divided a total of 1476 genotyped animals with phenotypes into five subsets. Each subset was used for prediction while the other four subsets were used for training in rotation. The genotypes included 675,686 autosomal variants from the Illumina Bovine 770K Bead chip, commonly used in genomic selection for Simmental beef cattle. We split the original 675K genotype panel into two groups based on whether the SNPs fell into functional classes. These two sets of SNPs were simultaneously incorporated into the BayesRC model to perform two-component genomic predictions. The Bayesian module is

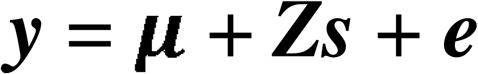

where ***y*** represents the vector of observation, ***Z*** is the incidence matrix allocating phenotypic observations to individuals, and ***e*** denotes random residual effects. ***s*** represents the sum of SNP effects obtained from diverse assumed distributions. BayesR assumes that SNP effects follow a mixture of four normal distributions 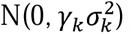, where the *γ_k_* are 0, 0.01, 0.1 and 1 with probabilities *π*_1_, *π*_2_, *π*_3_ and *π*_4_, respectively, and *π*_1_ + *π*_2_ + *π*_3_+ *π*_4_ =1 [91]. BayesRC was similar to BayesR except that a priori independent biological information was allocated to each variant [60]. Variant effects within functional classes are assumed to follow a mixture of four normal distributions with specific proportions, while variant effects outside functional classes follow an independent mixture of the four distributions.

### CBT Database building

The CBT Database features an intuitive and interactive web interface, developed using the R/Shiny framework, which provides powerful and accessible tools for the post-processing and visualization of transcriptome data. The database is structured around three R scripts: ui.R, server.R, and global.R. The ui.R defines the visual layout of the CBT Database interface and collects user inputs, which are then transmitted to the server side for processing. The server.R handles the calculations and generates the output, which is subsequently displayed on the interface. The global.R is responsible for loading the necessary R packages and functions, as well as importing the original data required for the database. The CBT Database is accessible at http://cattlegenomics.online/cattle_bodymap/. The complete set of R scripts used to create the interactive web portal is freely available at https://github.com/wentaocai/CBTDatabase.

## Declarations

### Ethics approval

All experiments and procedures were conducted in compliance with the regulations outlined by the Animal Care and Ethics Committee for Animal Experiments at the Institute of Animal Science, Chinese Academy of Agricultural Sciences (Permit Number: IAS2021-43).

### Availability of data and materials

The 400 RNA-seq data supporting the findings of this study are available in the National Genomics Data Center (NGDC) database (https://ngdc.cncb.ac.cn/bioproject/browse/), with accession numbers listed in Table S1. This dataset includes 124 samples from newborns (CRA016956), 145 samples from young individuals (CRA017003), and 131 samples from adults (CRA028008). Additionally, the whole genome sequencing data for the nine cattle generated in this study can be accessed under accession number CRA017018. The gene expression, lncRNA, RNA editing, and splicing data is accessible at CBT Database ((http://cattlegenomics.online/cattle_bodymap/). Each use of software programs has been clearly indicated and information on the options that were used is provided in the Methods section. The complete set of R scripts used to create the interactive web portal is freely available at https://github.com/wentaocai/CBTDatabase.

### Code availability

The computational pipelines and code for identifying splicing events, lncRNAs, and RNA editing sites, along with the implemented methods for genetic variance analysis and genomic prediction, had been deposited in the BioCode repository (Accession: BT007979) at the National Genomics Data Center (NGDC). These resources are publicly available through the official portal at https://ngdc.cncb.ac.cn/biocode.

### CRediT author statement

**Wentao Cai**: Conceptualization, Methodology, Software, Formal analysis, Writing - Original Draft. **Yapeng Zhang**: Investigation, Resources. **Lei Xu**: Investigation, Data Curation. **Yahui Wang**: Investigation. **Xin Hu**: Investigation. **Qian Li**: Data Curation. **Linxi Zhu**: Data Curation. **Zezhao Wang**: Data Curation. **Huijiang Gao**: Resources. **Lingyang Xu**: Methodology. **Junya Li**: Supervision, Funding acquisition, Project administration. **Lupei Zhang**: Conceptualization, Methodology, Writing - Review & Editing, Funding acquisition.

## Acknowledgements

This work was supported by grants from the Fund of the National Natural Science Foundation of China (32202652 and 32372833), China Agriculture Research System of MOF and MARA (CARS-37), Science and Technology Innovation Project of the Chinese Academy of Agricultural Sciences (ASTIP-2021-IAS-03) and the Program of Anhui Provincial Key Laboratory of Livestock and Poultry Product Safety Engineering (XM2405). We are grateful to all members of the Cattle Breeding Innovative Research Team for sample collection and data statistics as well as all staff at the cattle experimental unit in Shayang for animal fattening.

## Declaration of AI and AI-assisted technologies in the writing process

No AI or AI-assisted technologies were used in the writing or editing of this manuscript.

