## Supplementary figures and images for "A Cattle BodyMap of Transcriptome across 52 Tissues and 3 Developmental Stages Reveals New Genetic Insights into Beef Production Traits"

### FigureS1.tif

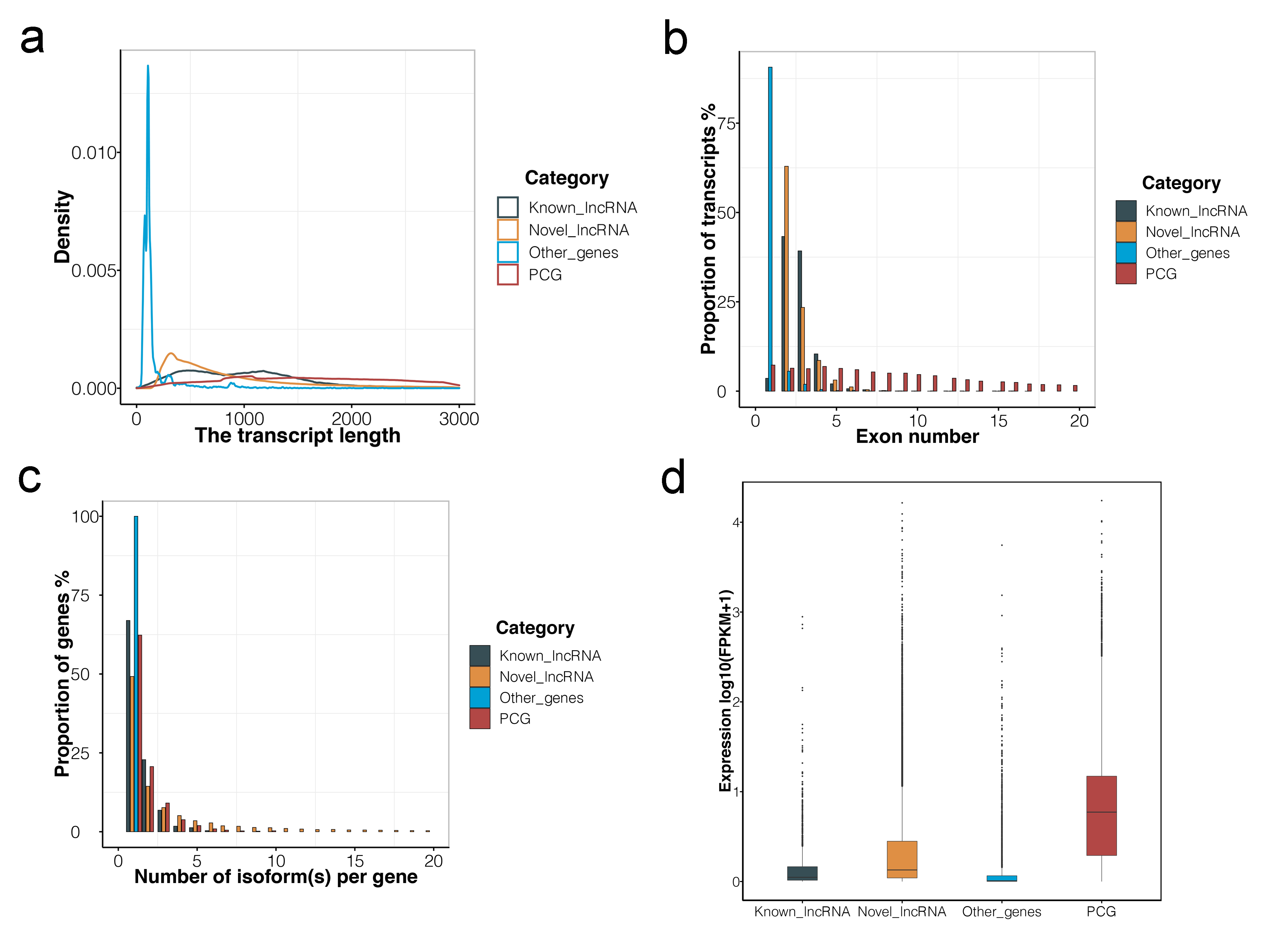

### FigureS2.tif

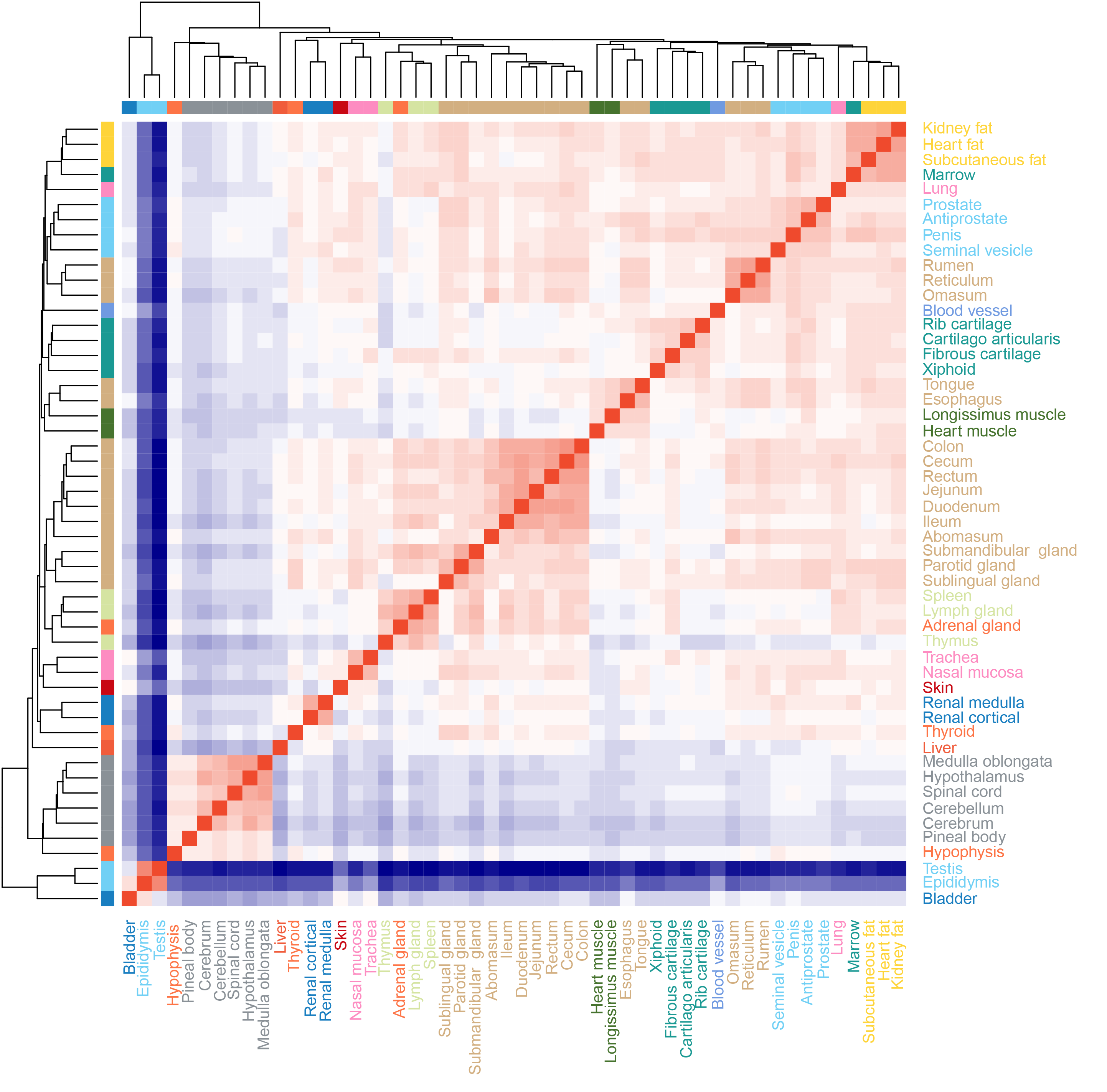

### FigureS3.tif

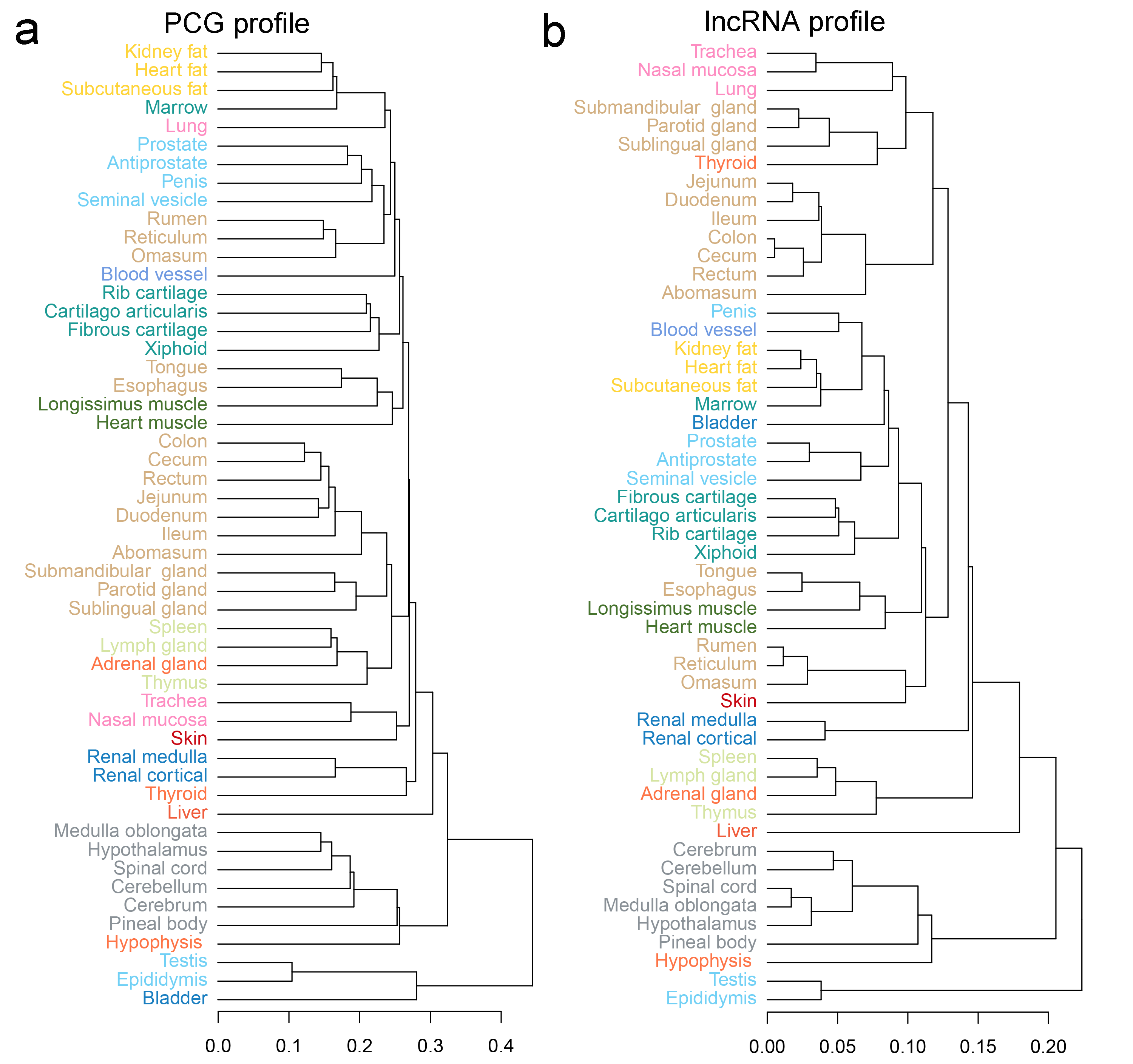

### FigureS4.tiff

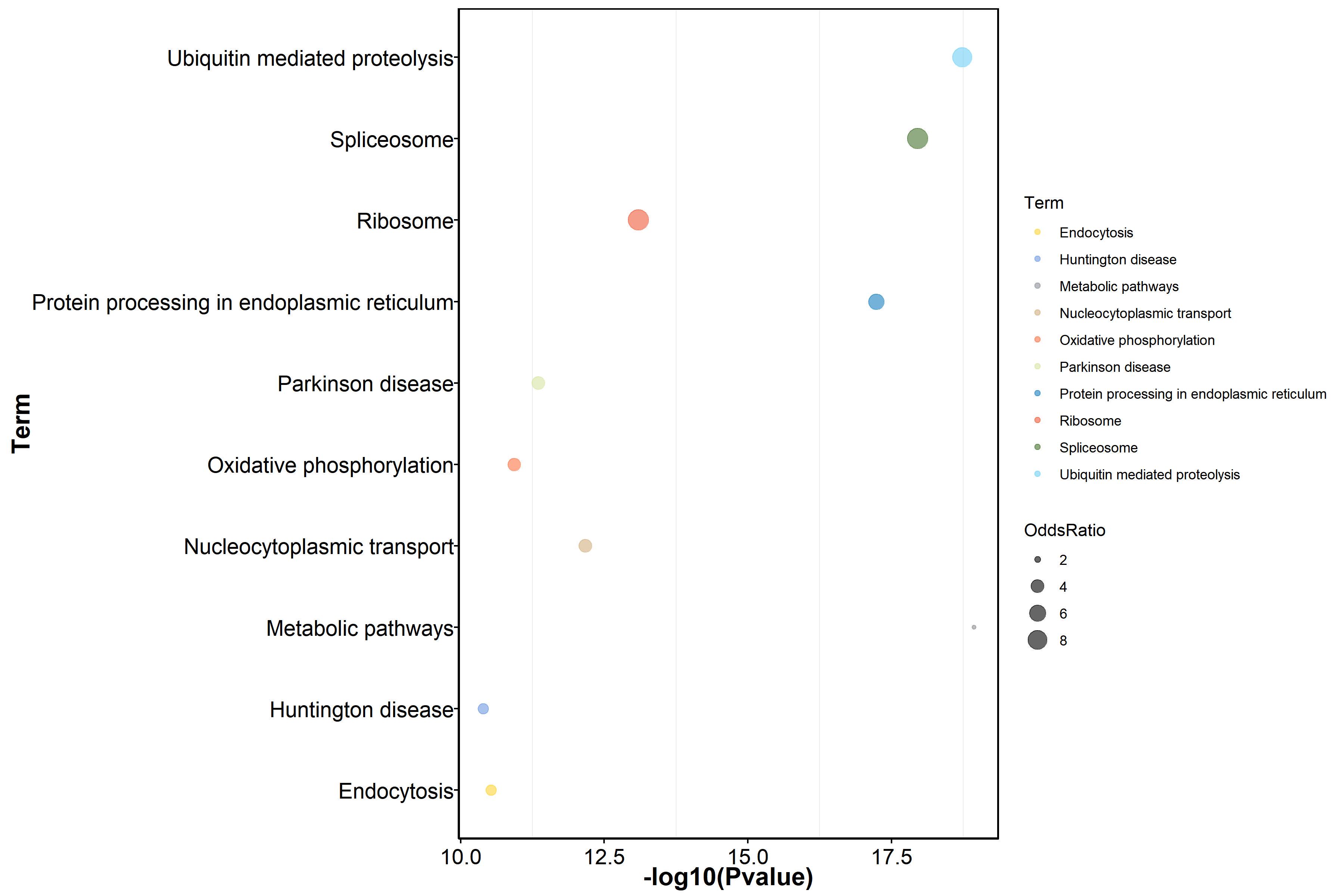

### FigureS5.tif

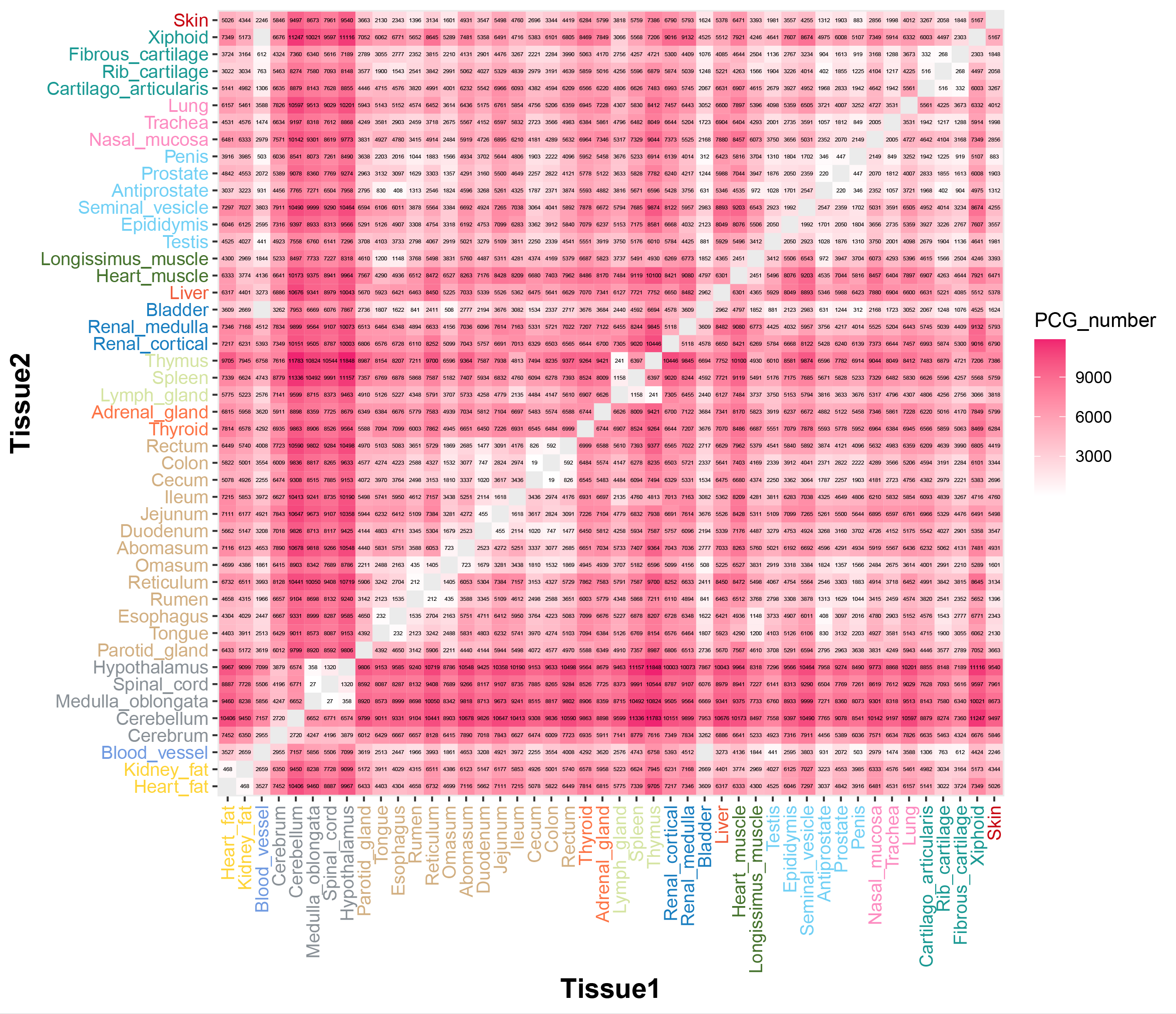

### FigureS6.tif

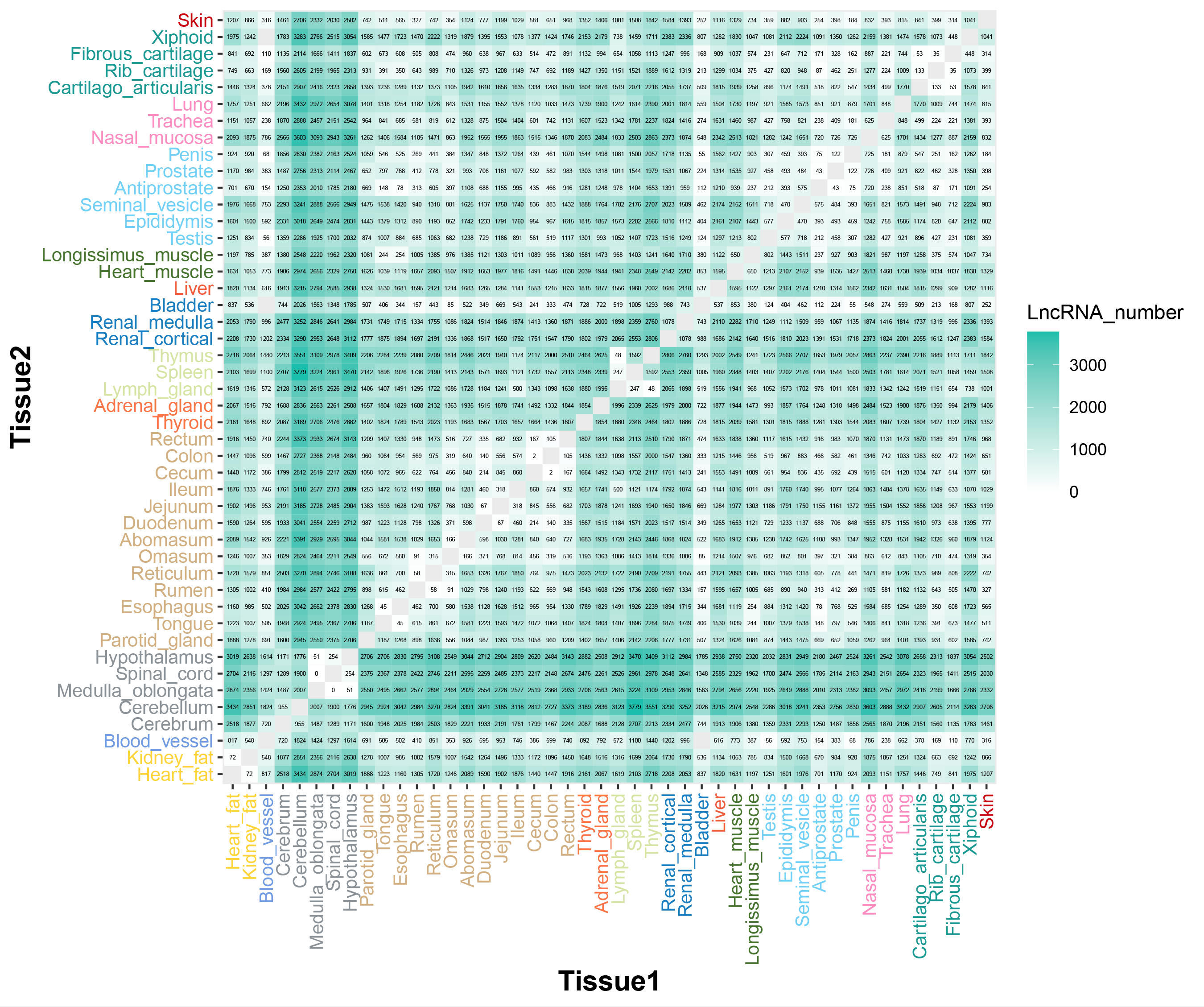

### FigureS7.tif

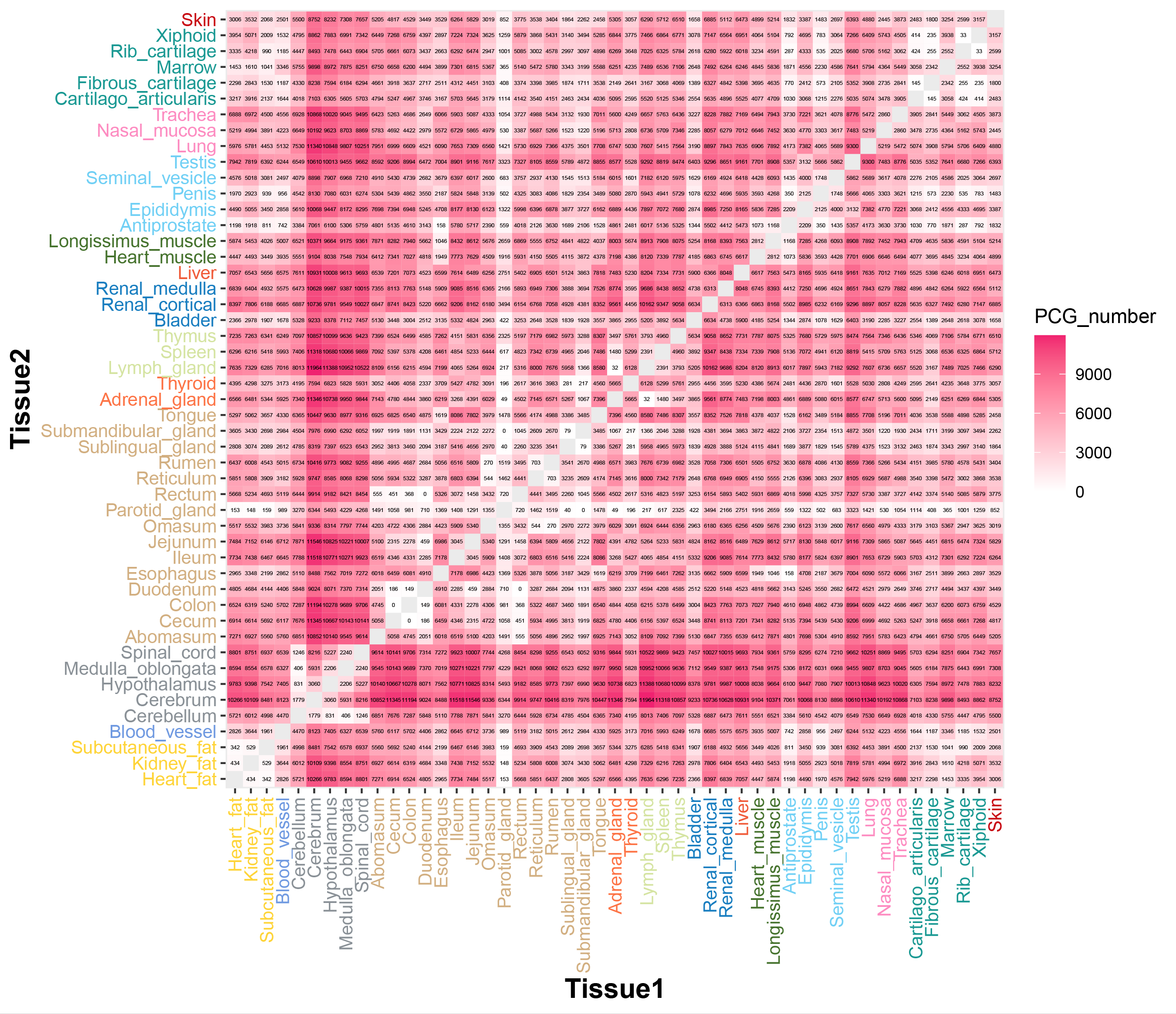

### FigureS8.tif

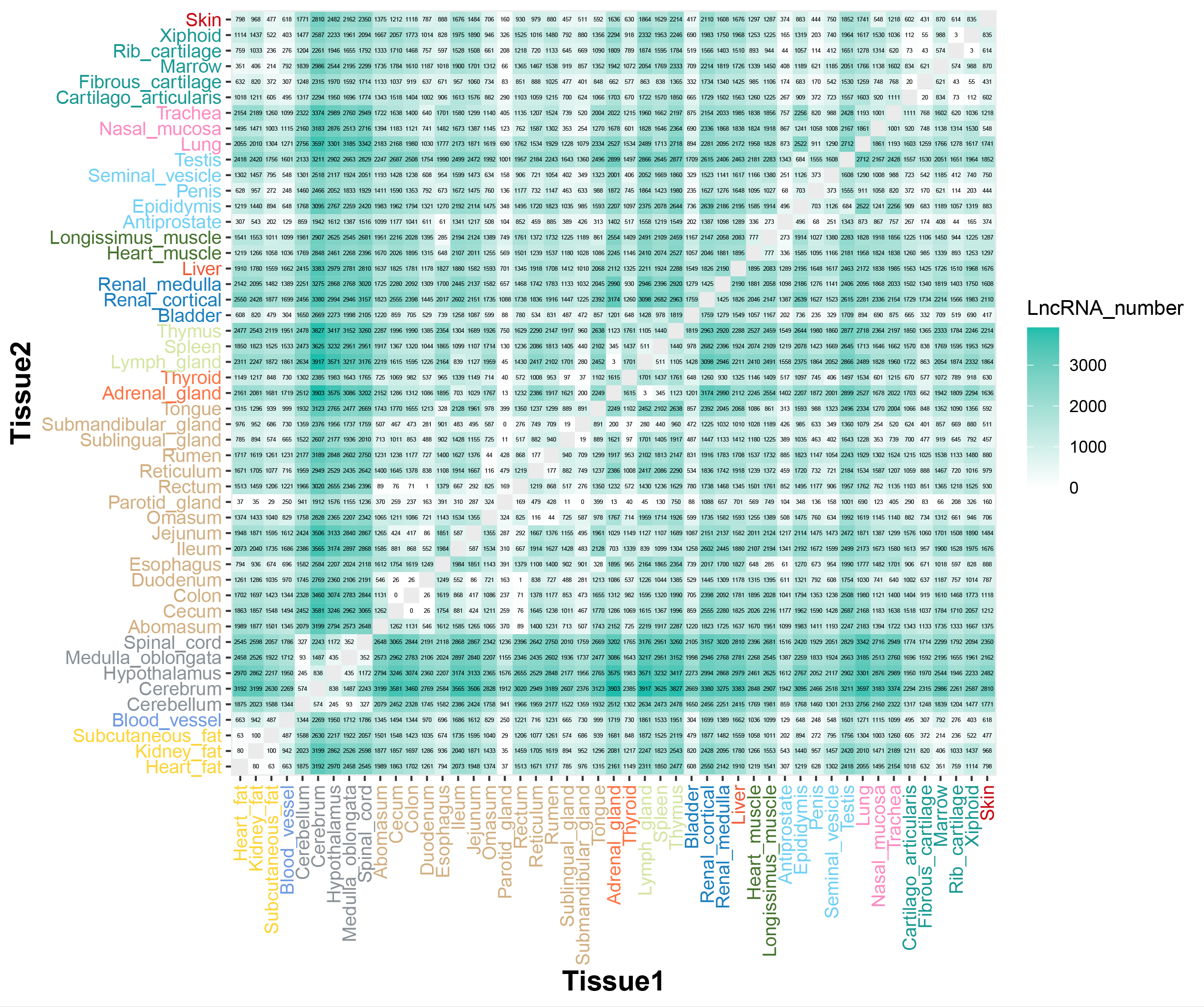

### FigureS9.tif

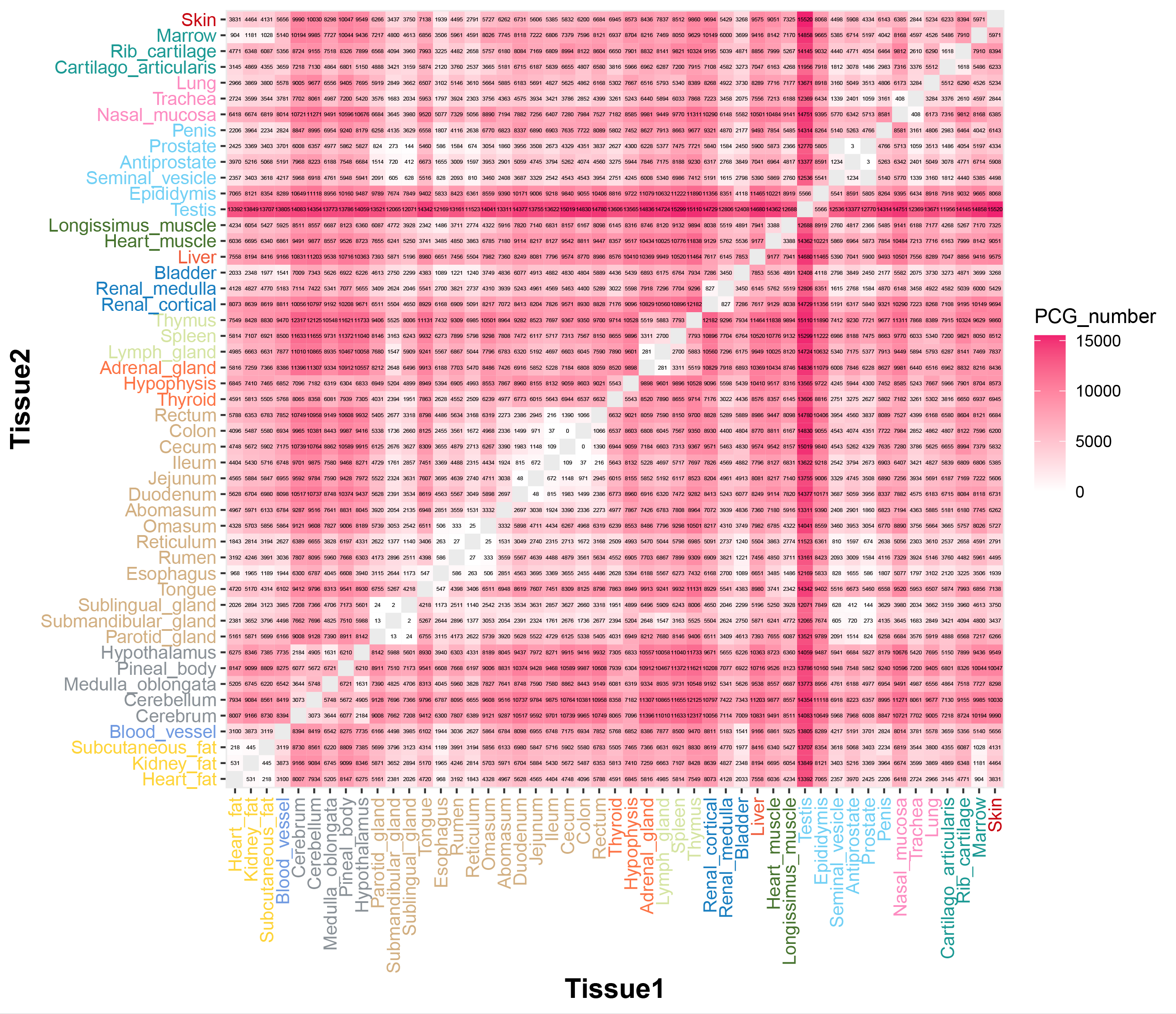

### FigureS10.tif

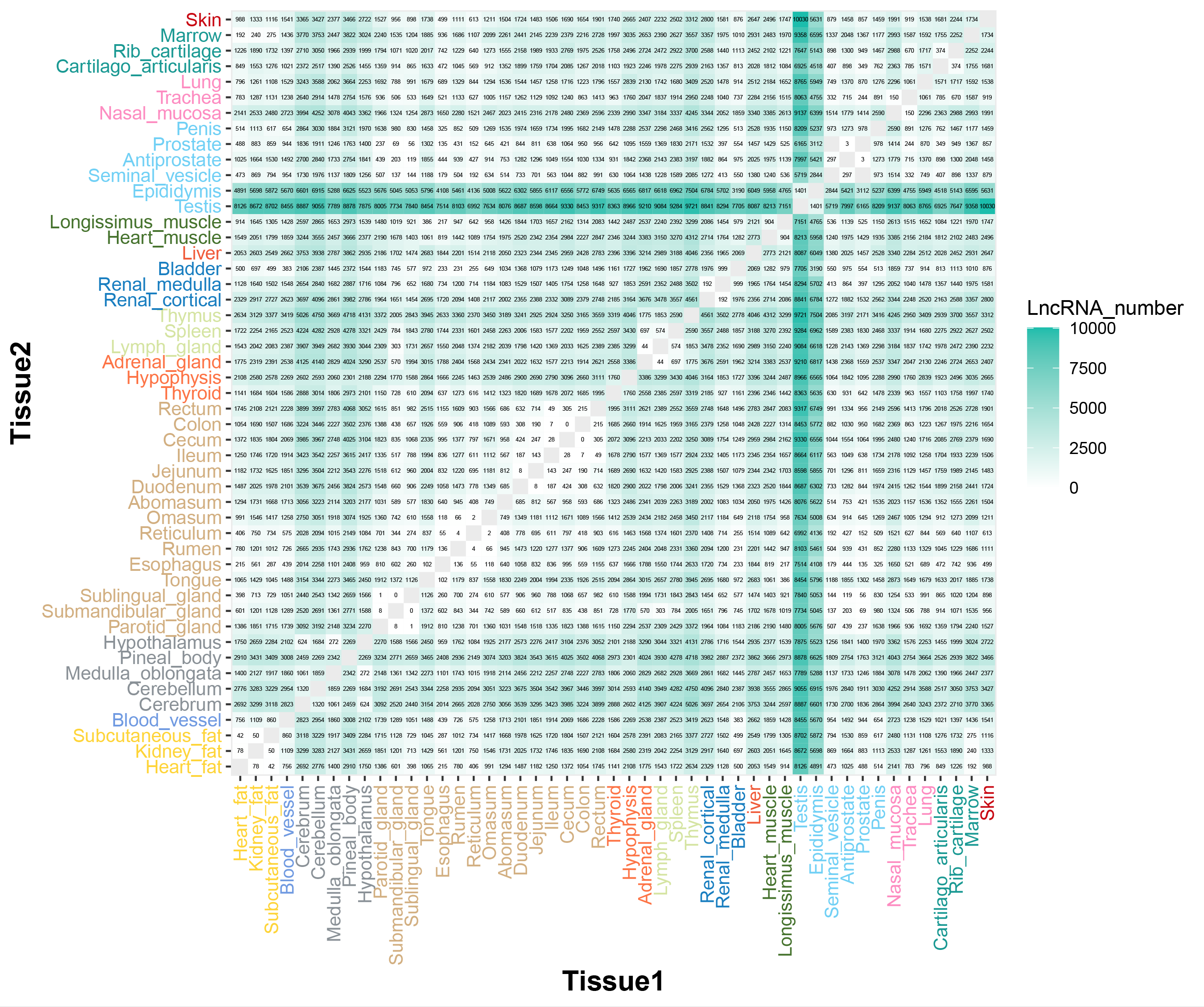

### FigureS11.tif

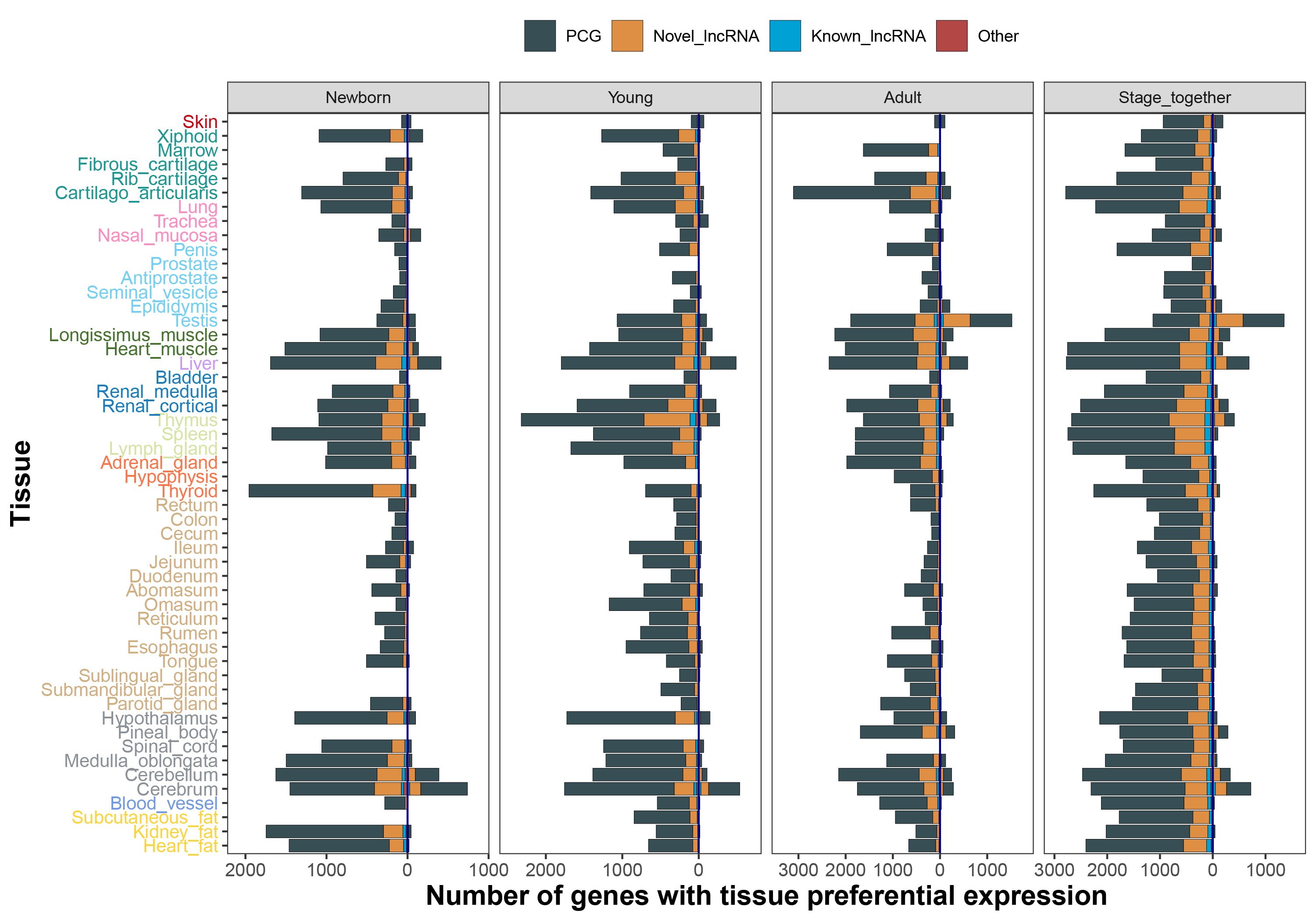

### FigureS12.tif

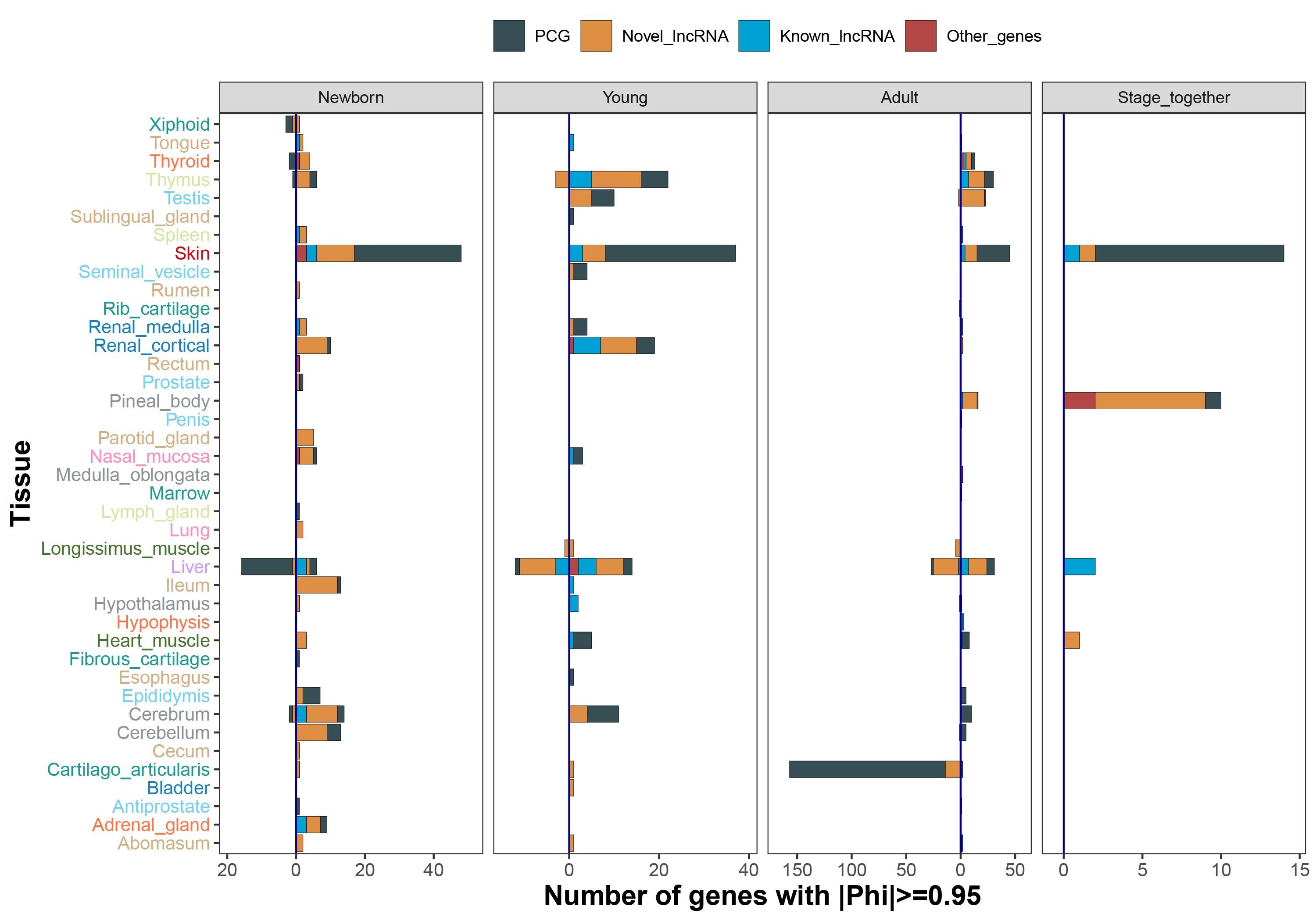

### FigureS13.tif

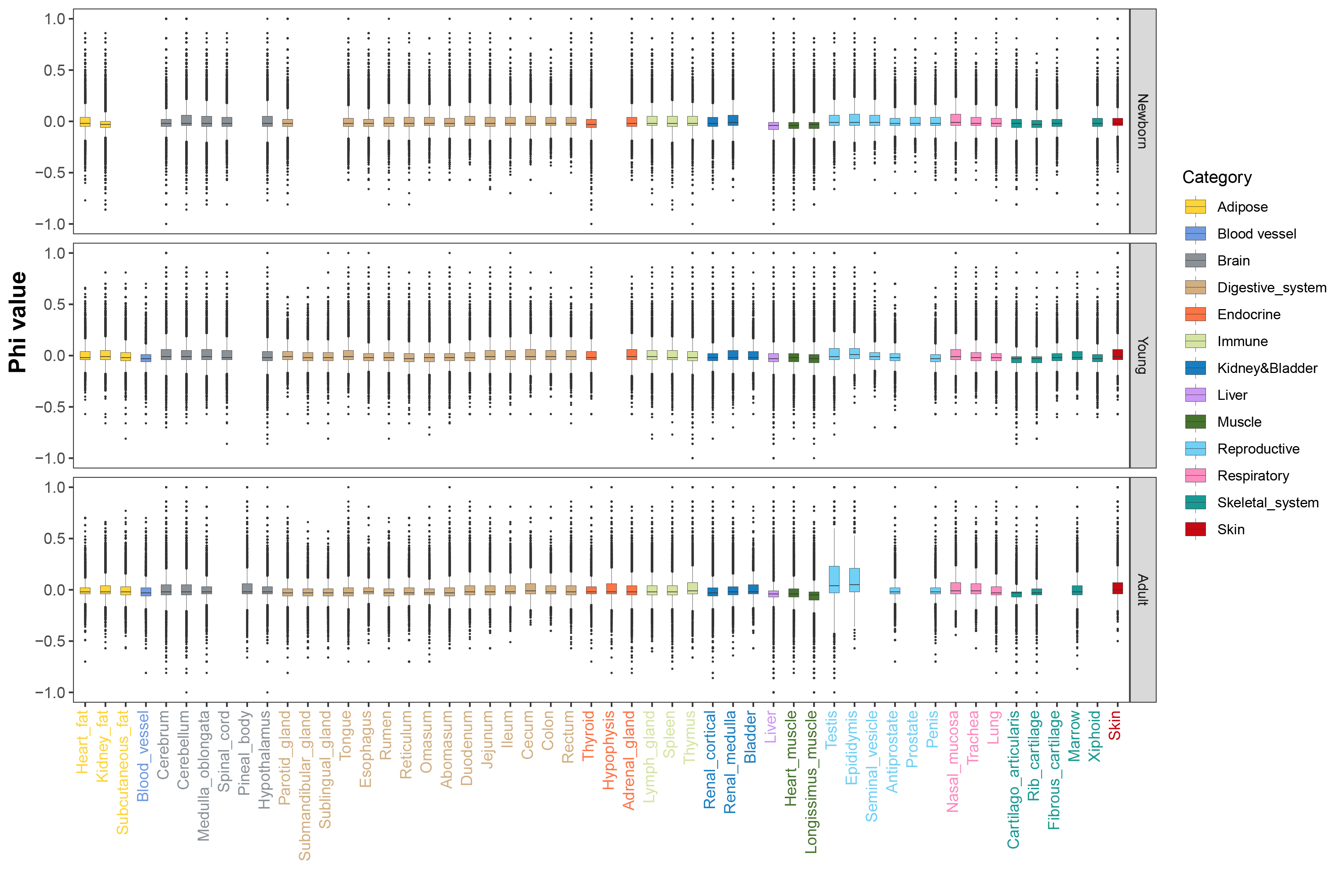

### FigureS14.tif

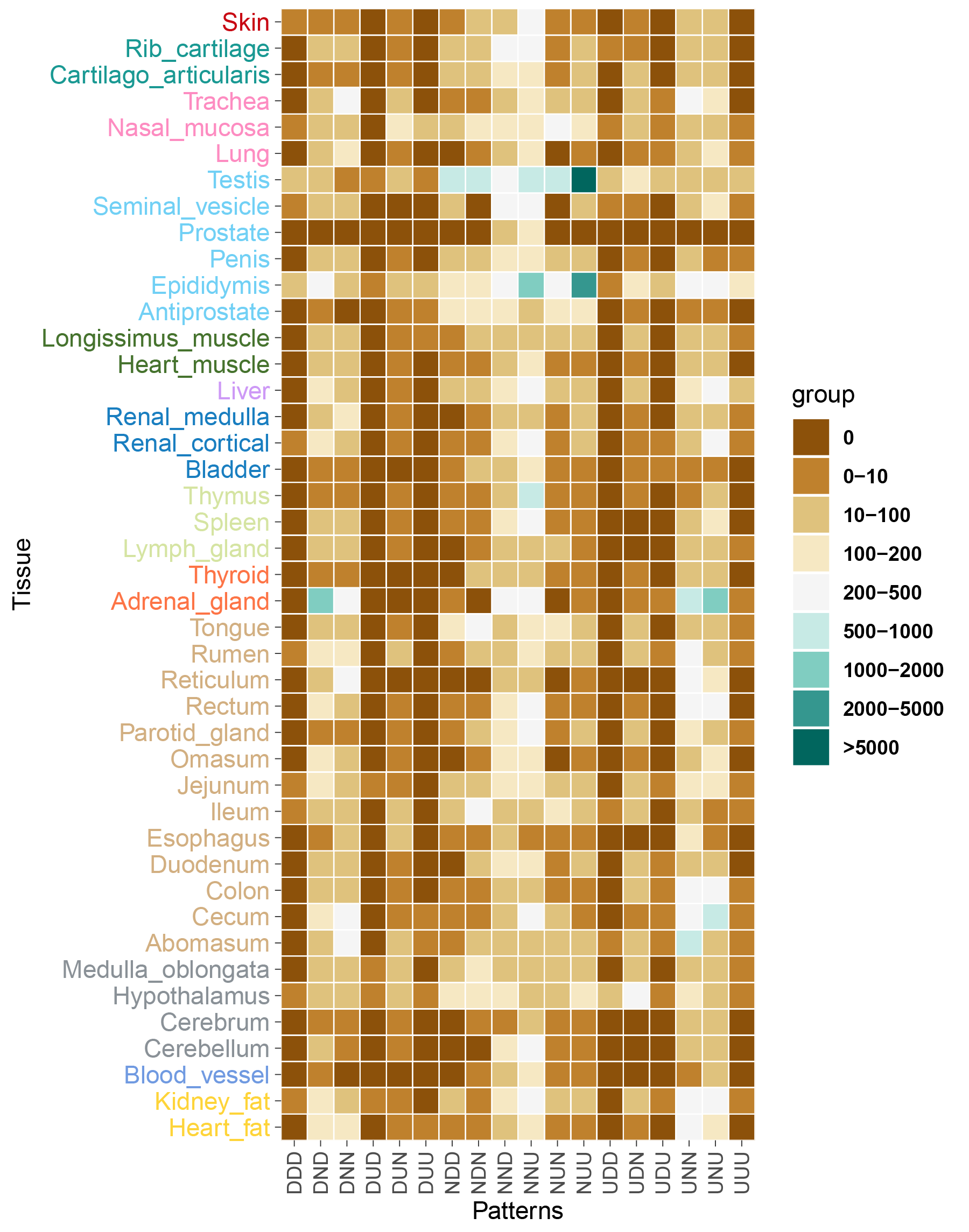

### FigureS15.tif

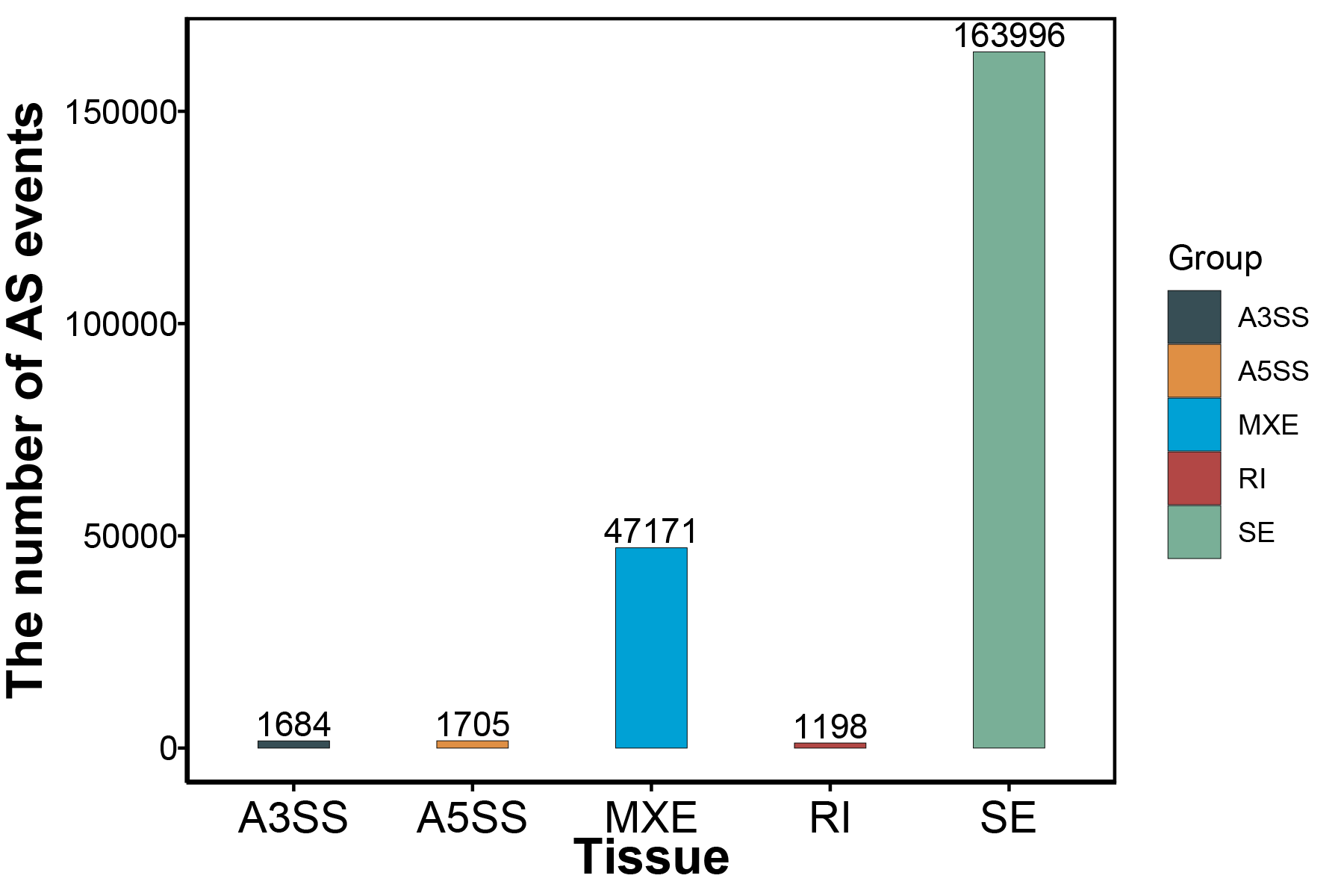

### FigureS16.tif

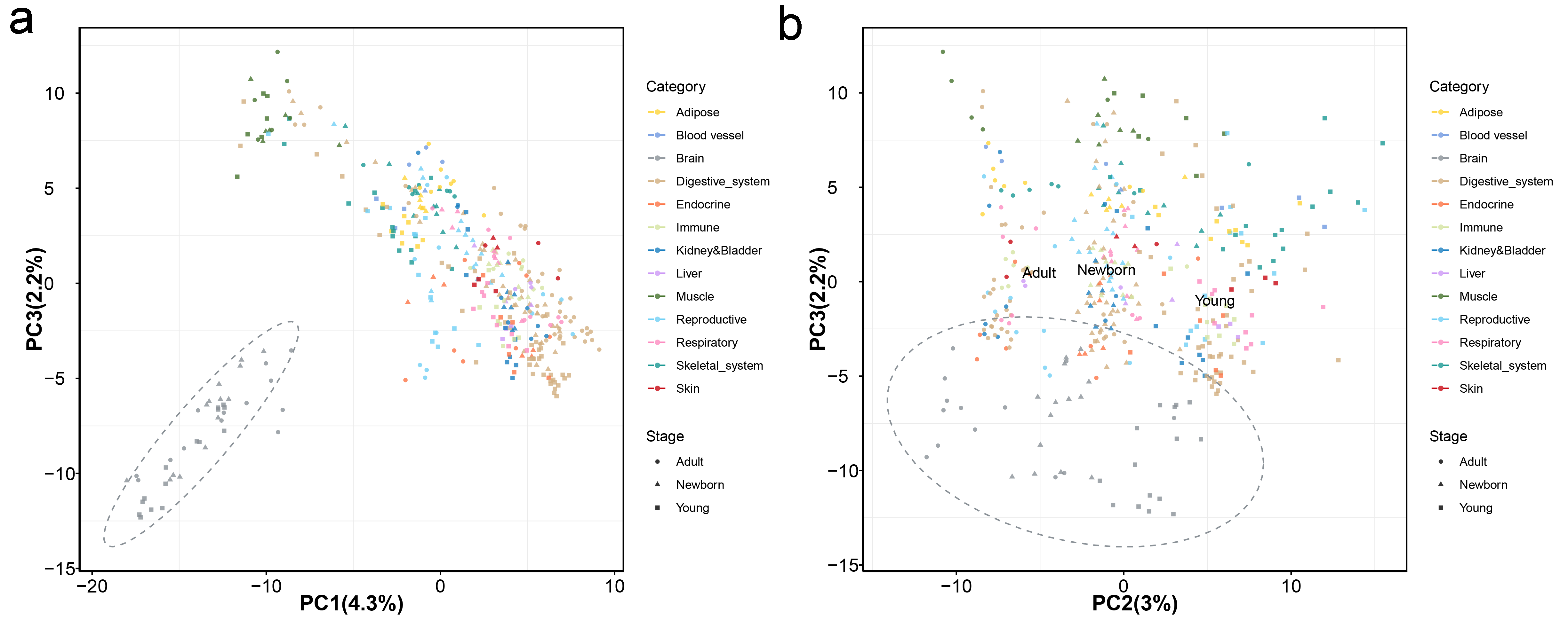

### FigureS17.tif

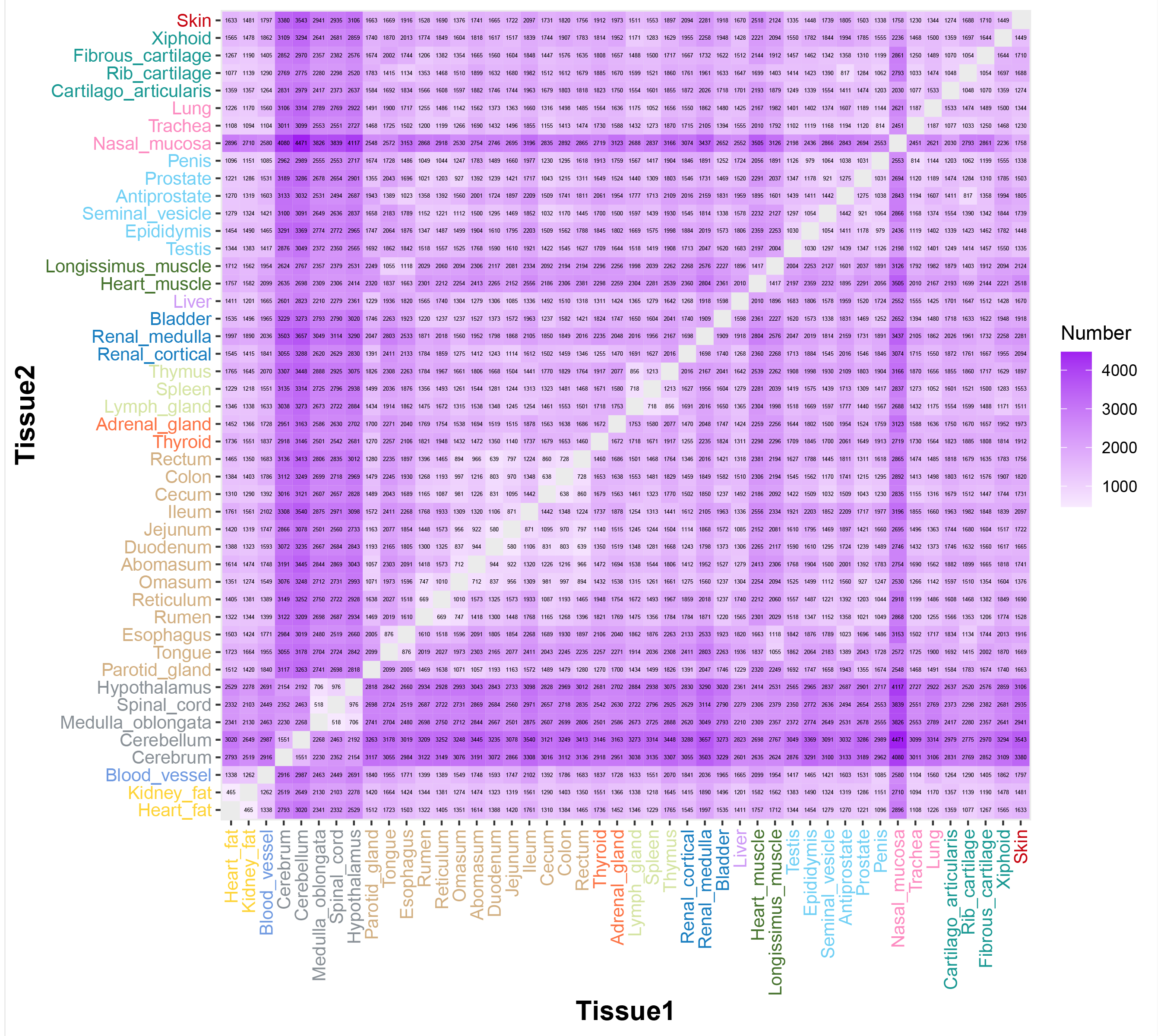

### FigureS18.tif

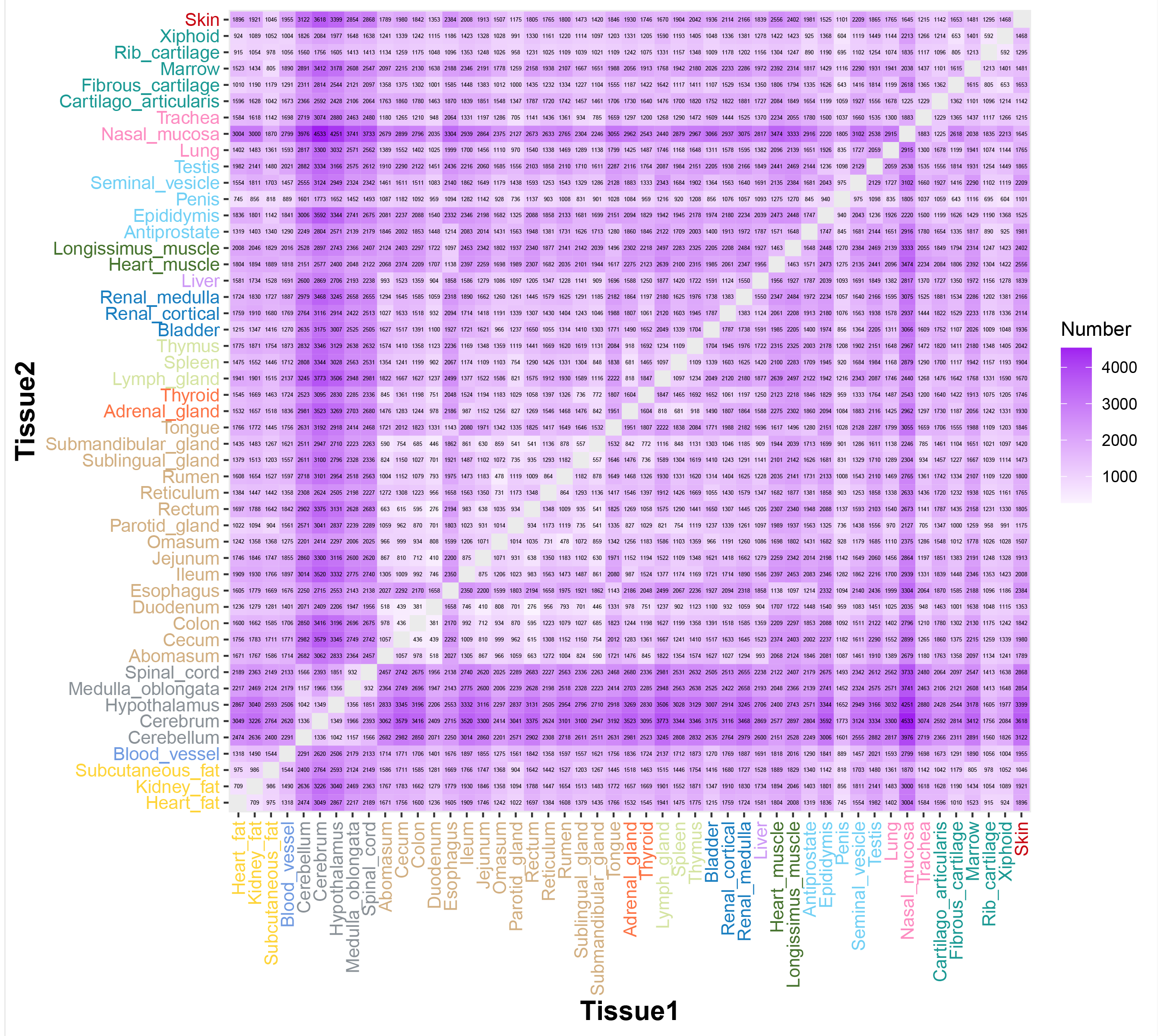

### FigureS19.tif

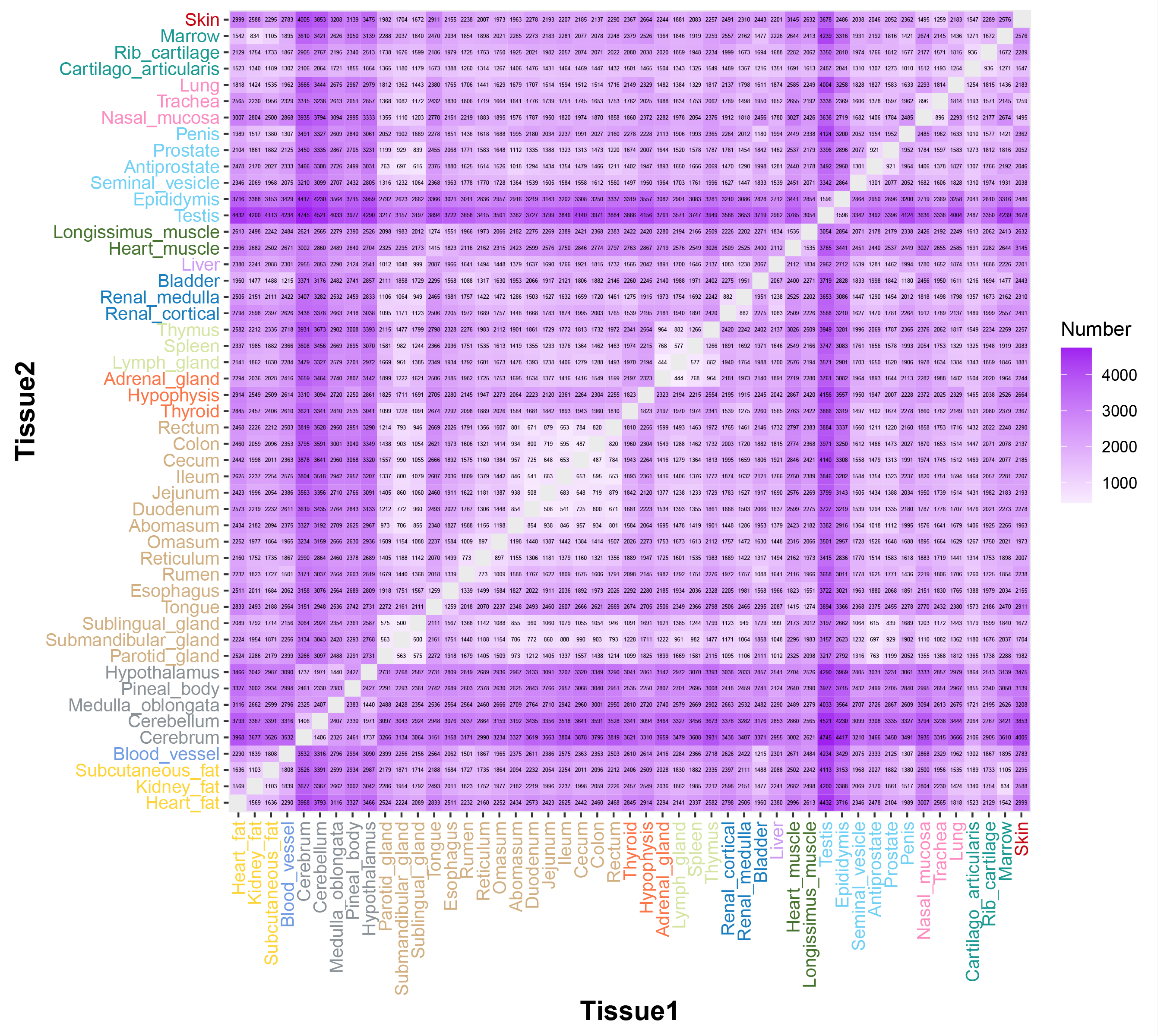

### FigureS20.tif

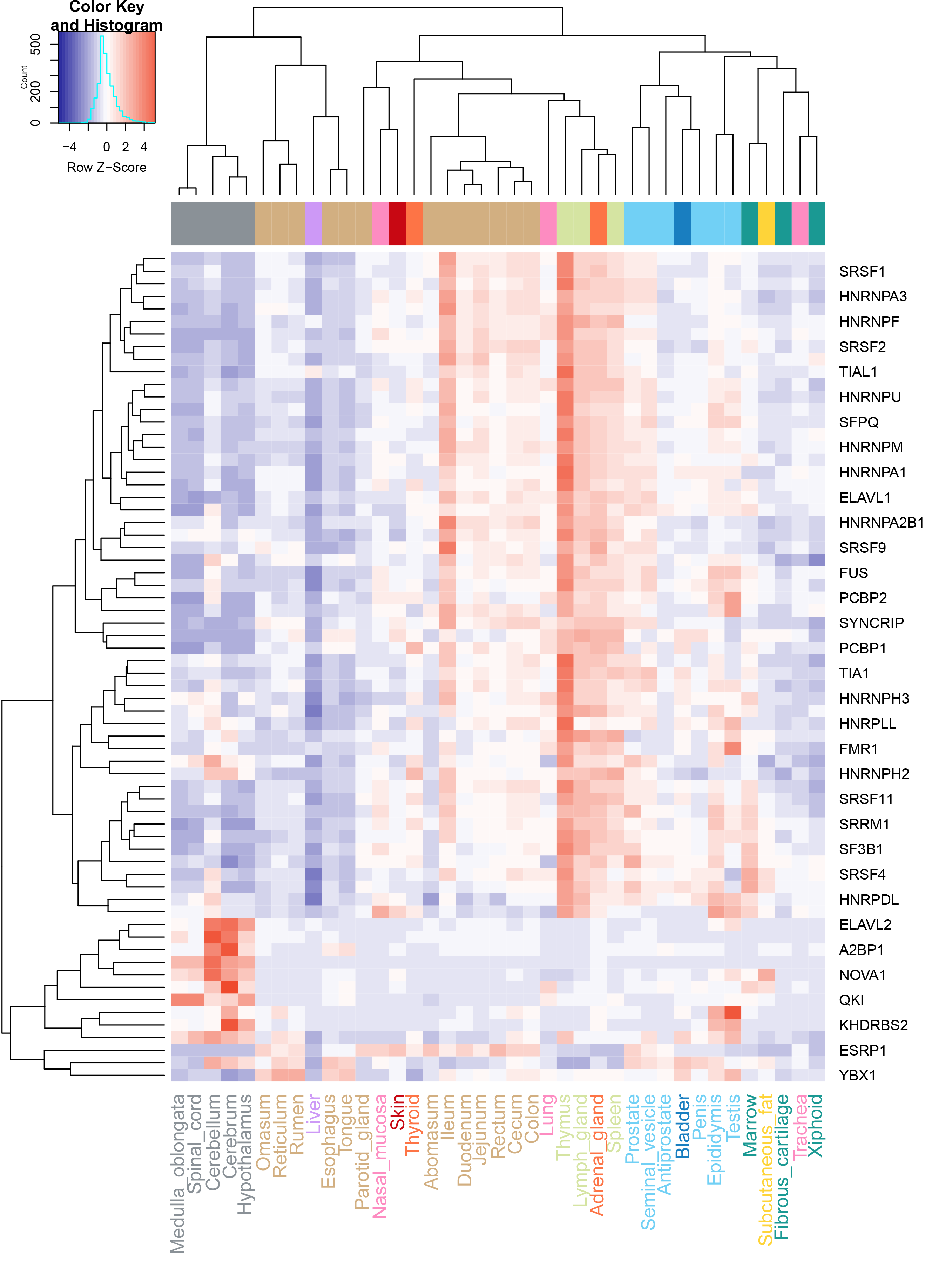

### FigureS21.tif

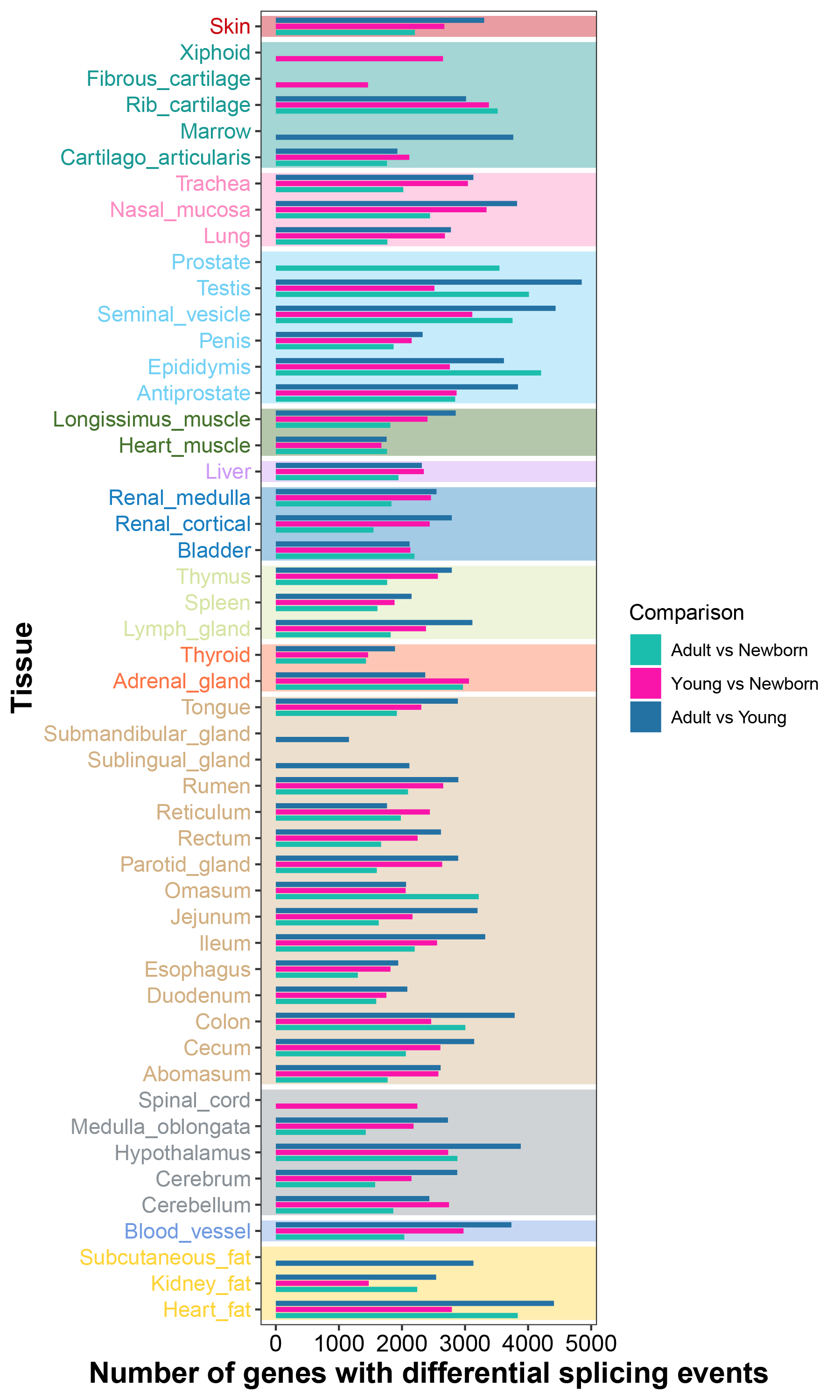

### FigureS22.tif

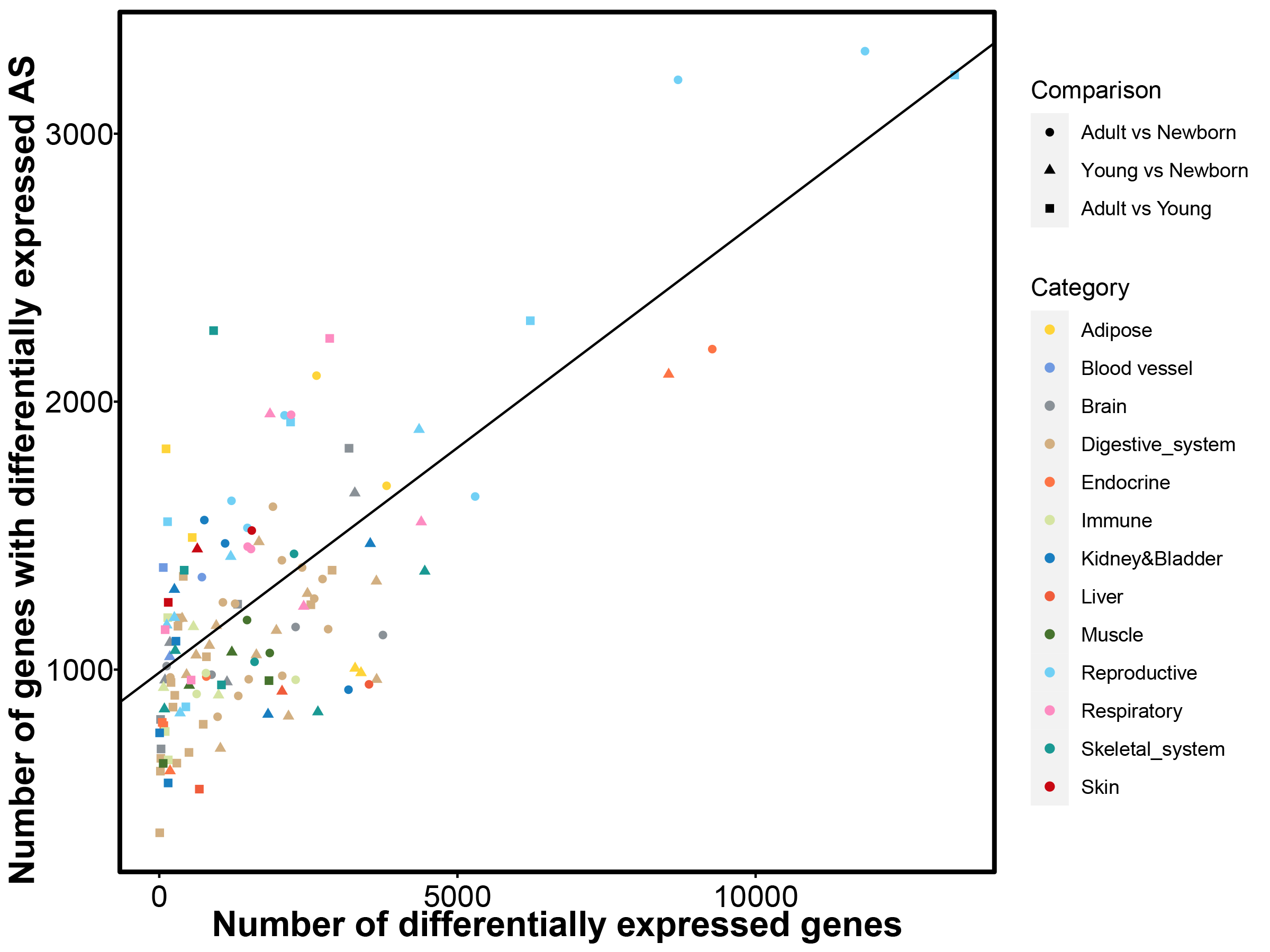

### FigureS23.tif

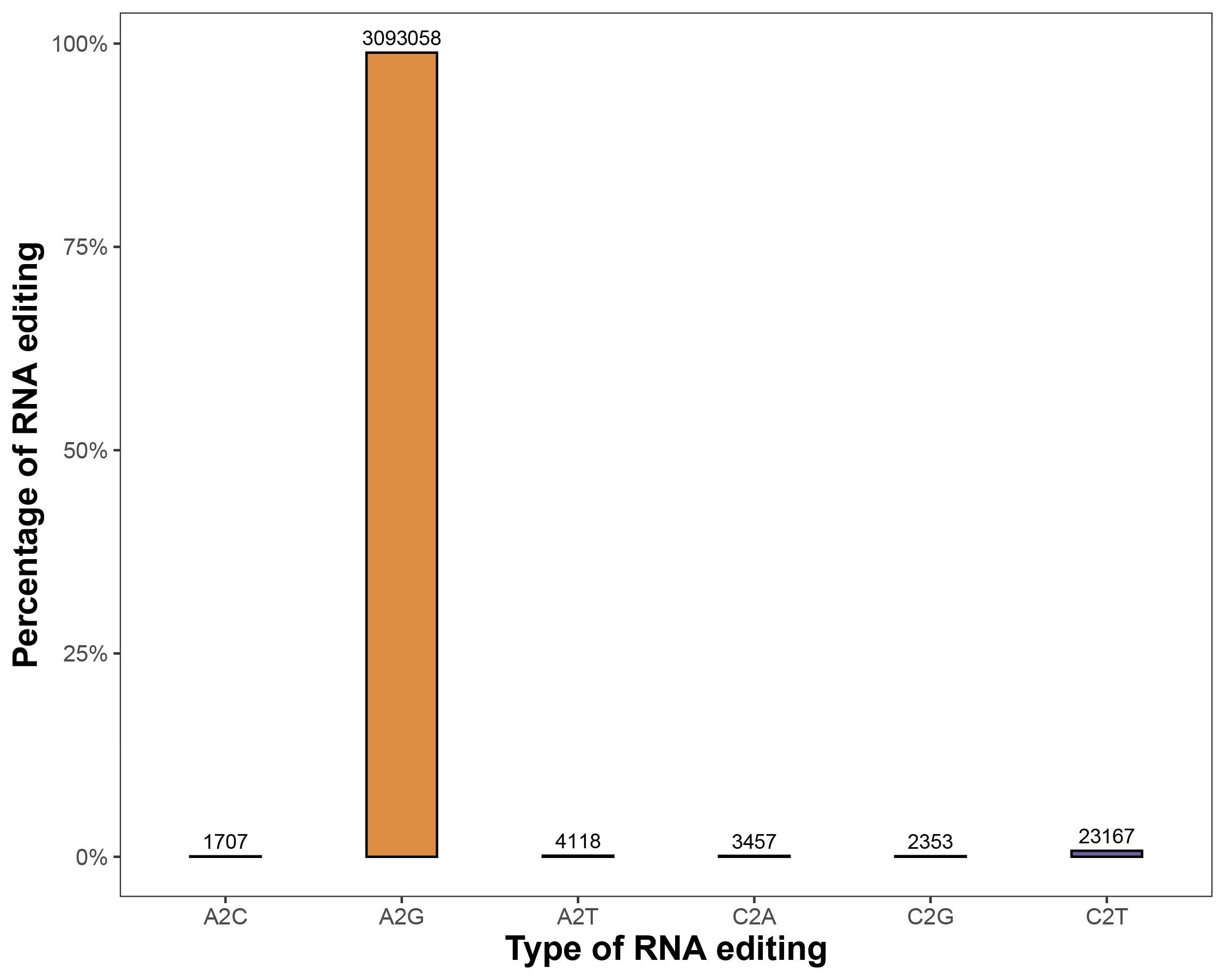

### FigureS24.tif

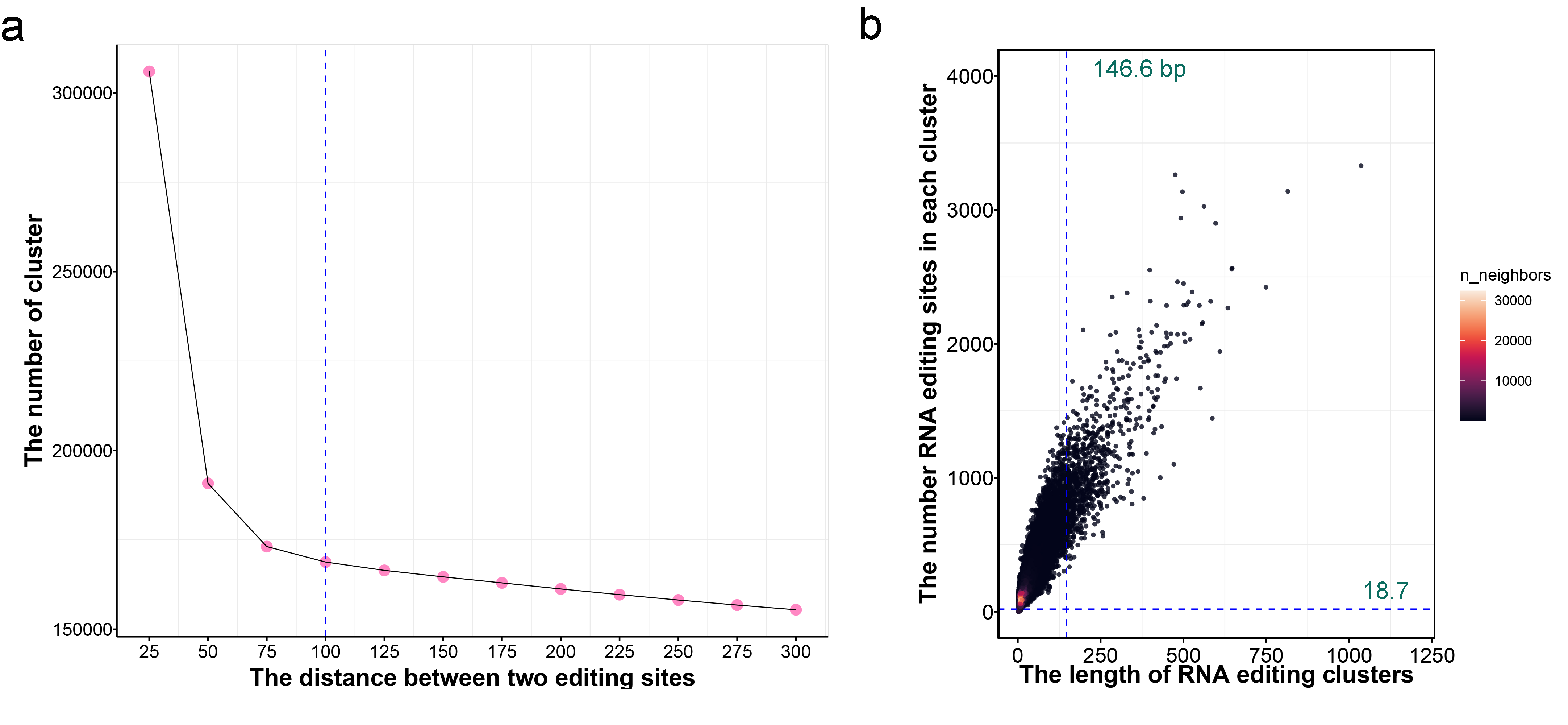

### FigureS25.tif

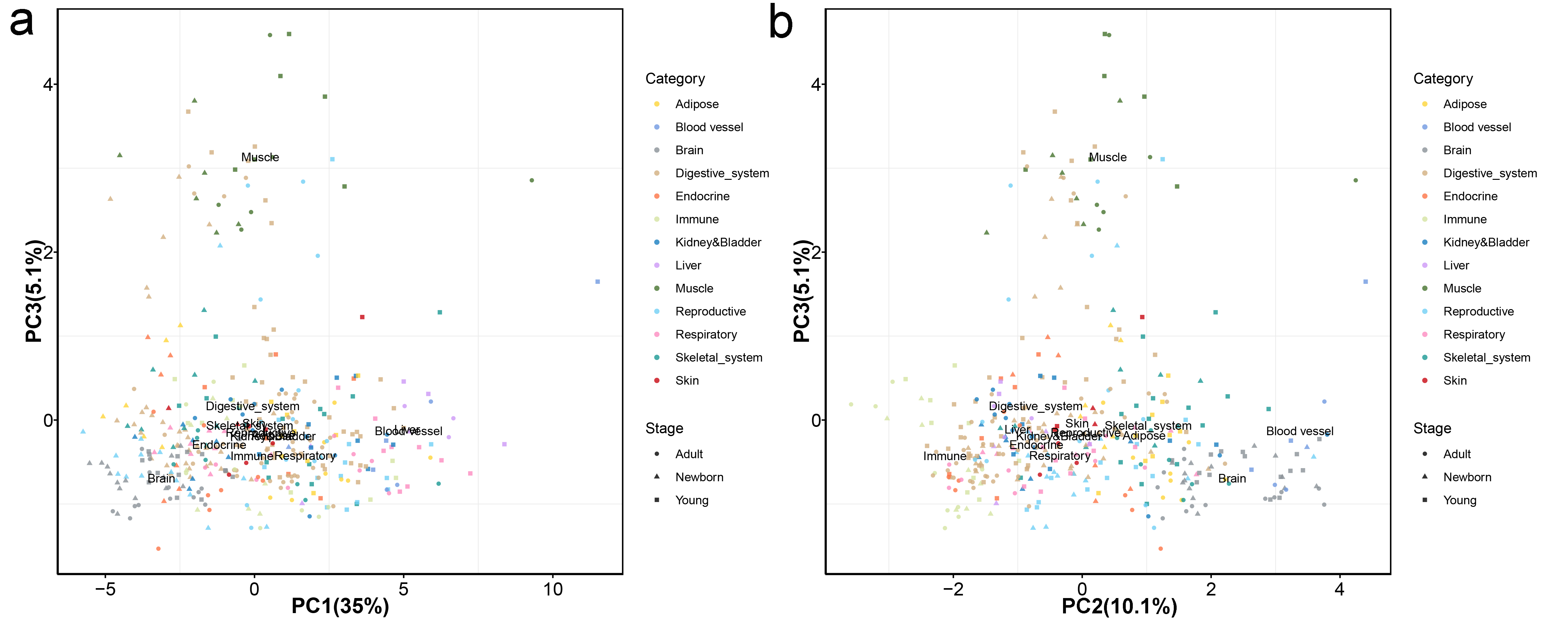

### FigureS26.tif

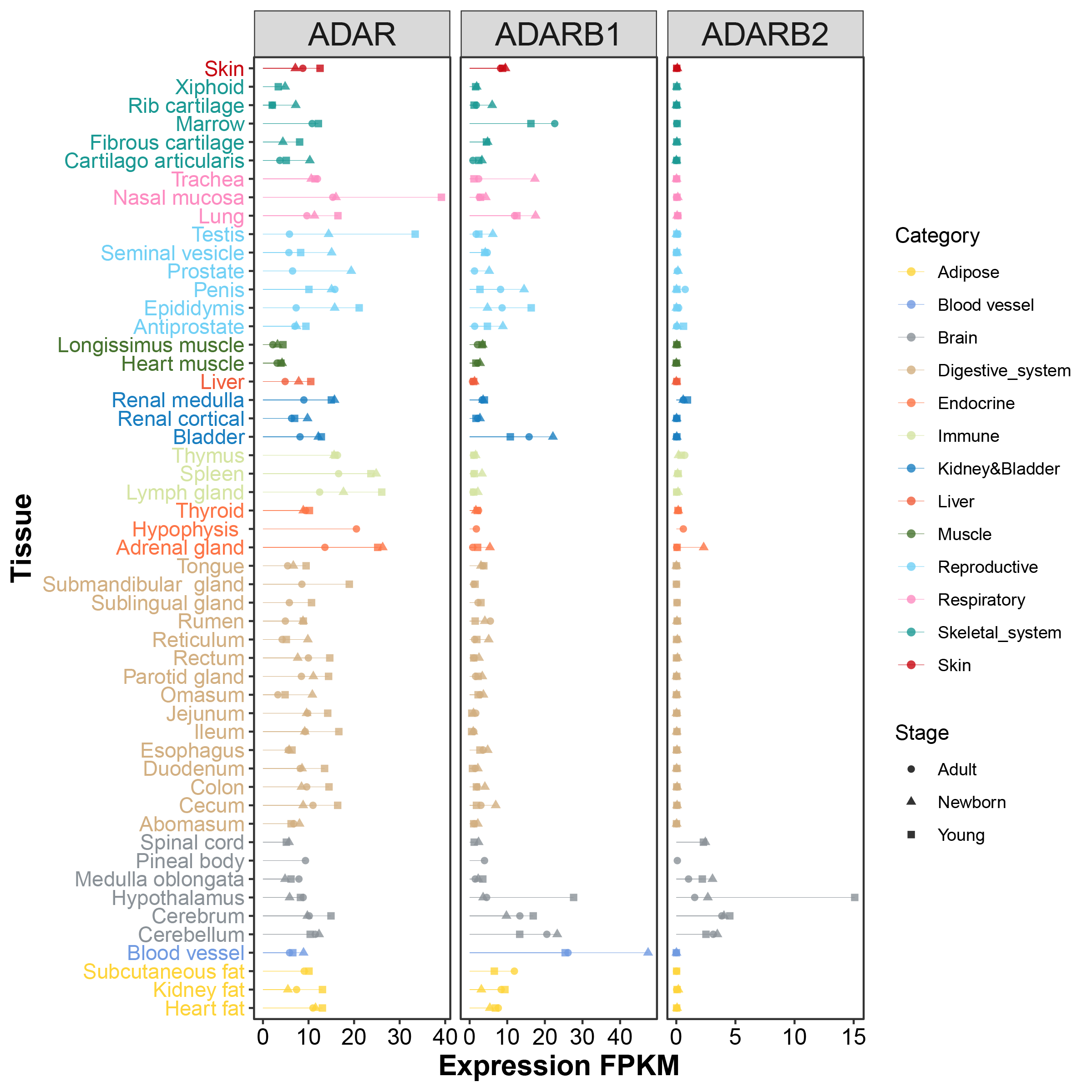

### FigureS27.tif

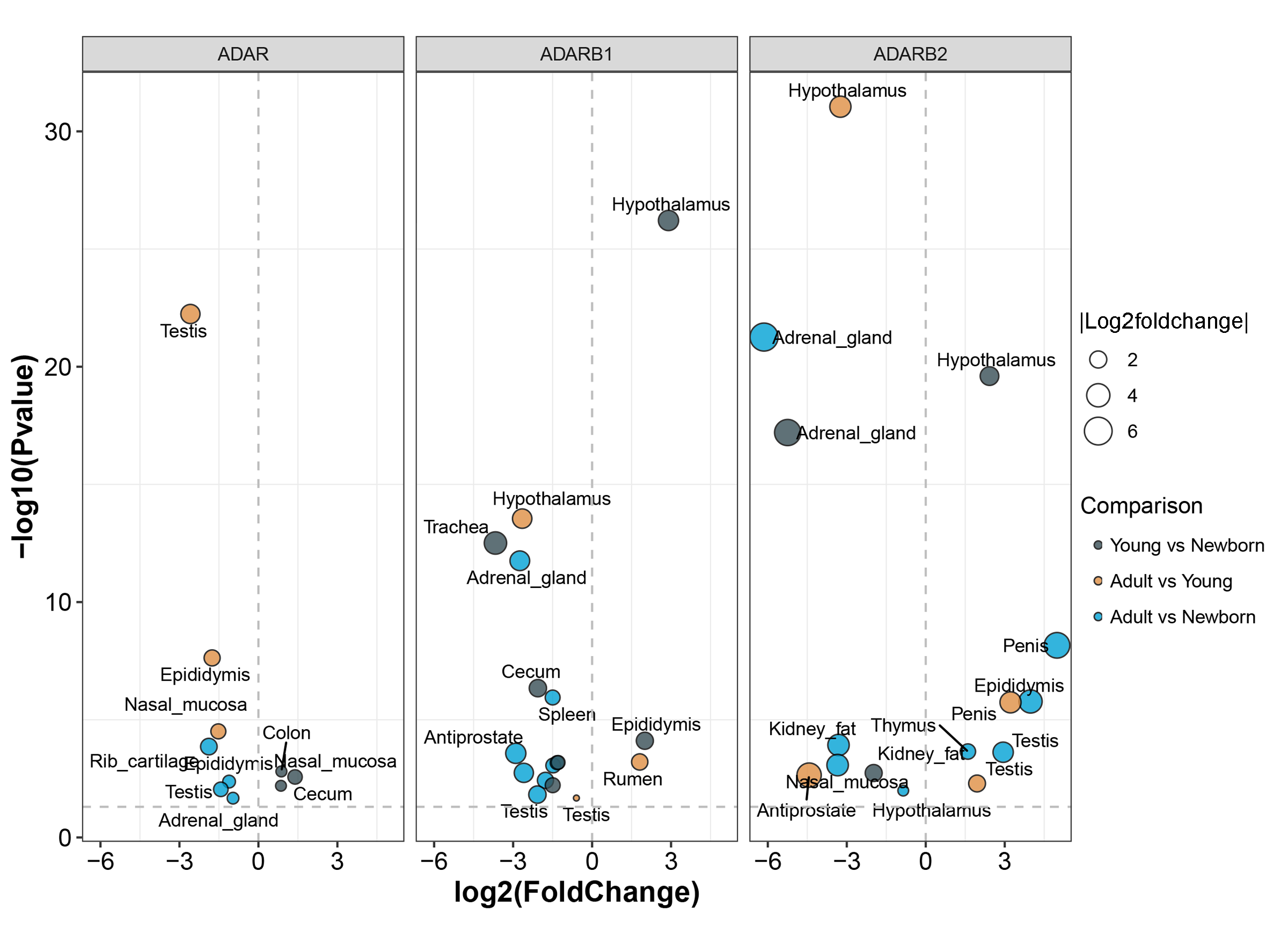

### FigureS28.tif

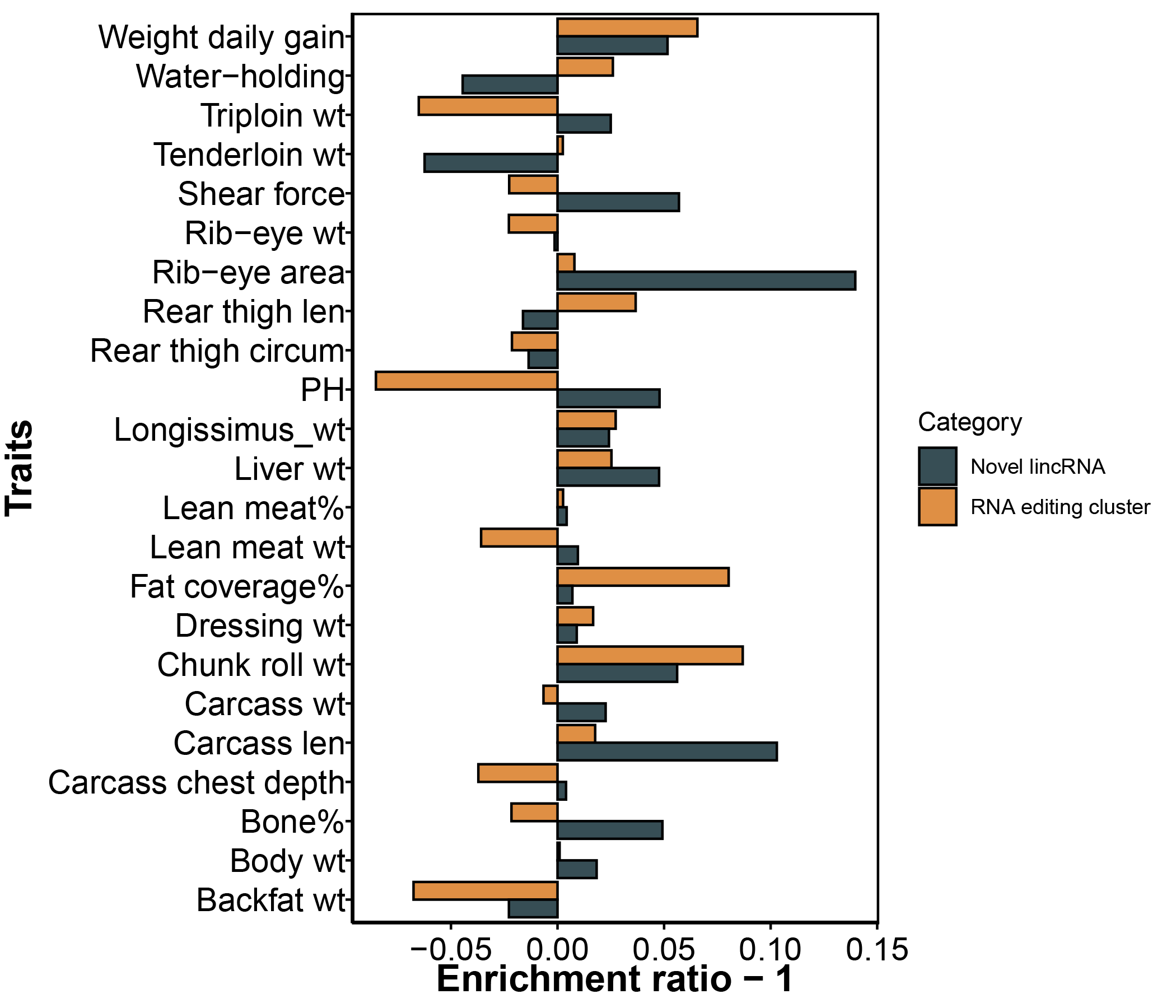

### FigureS29.tif

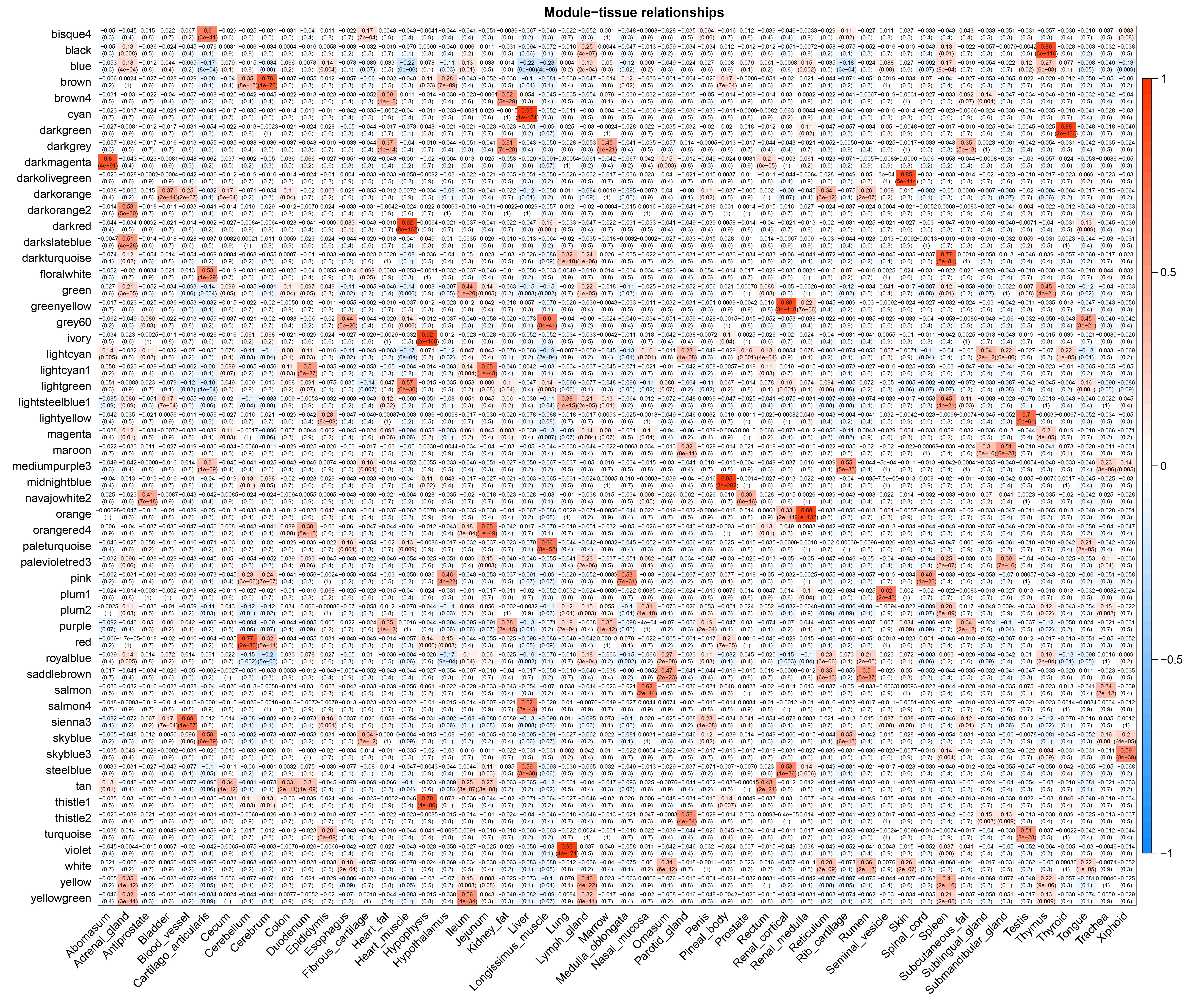

### FigureS30.tif

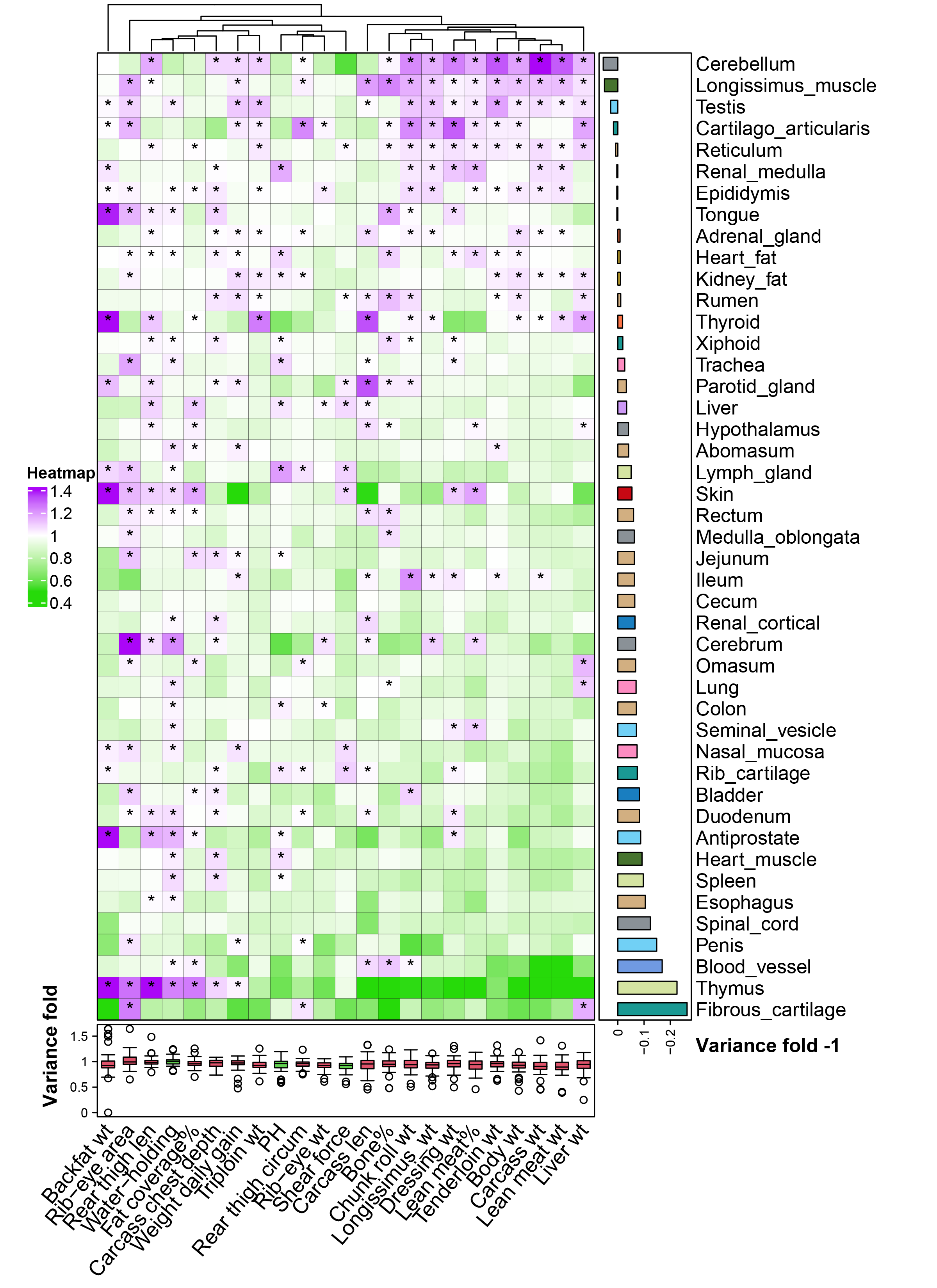

### FigureS31.tif

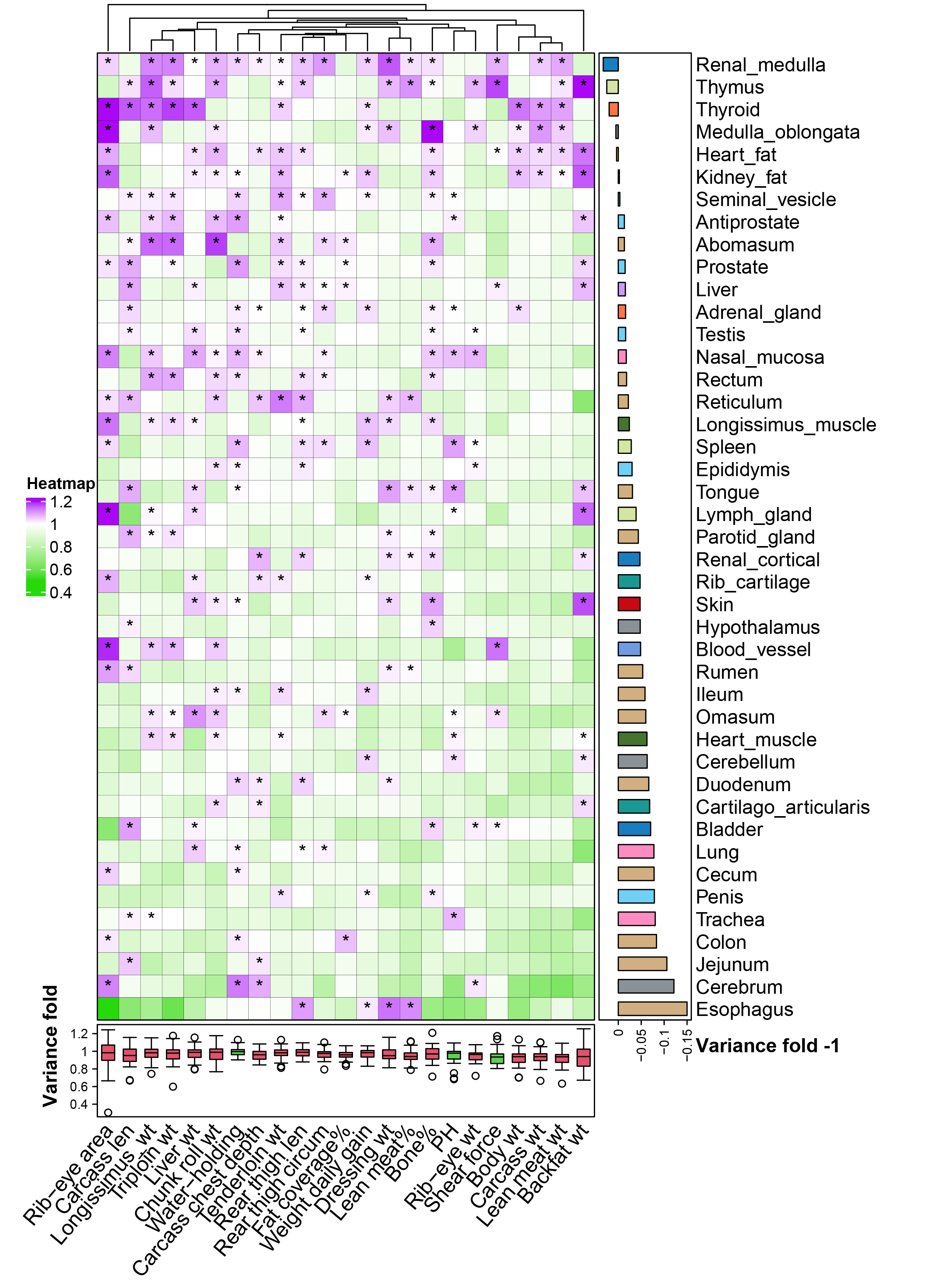

### FigureS32.tif

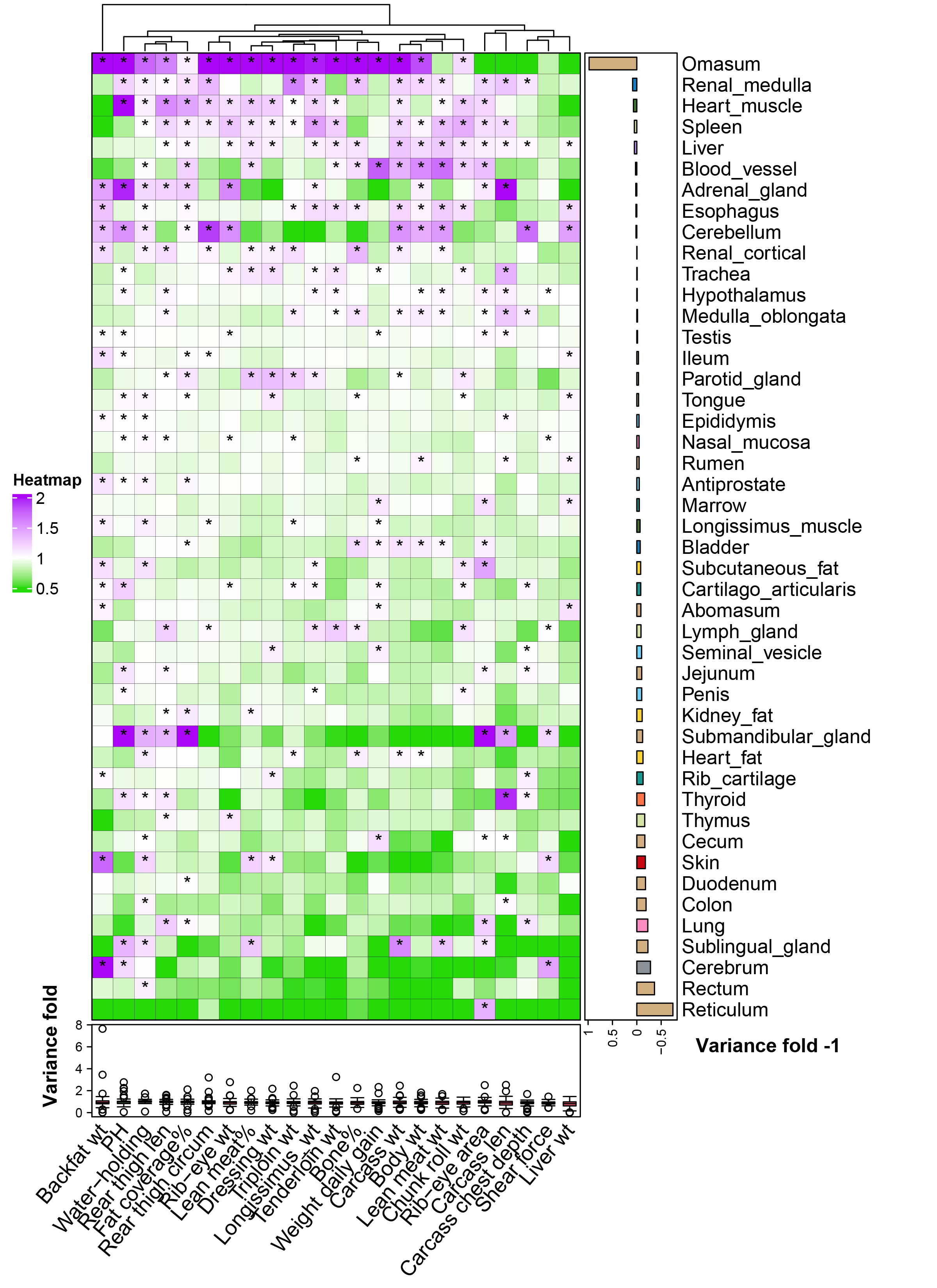

### FigureS33.tif

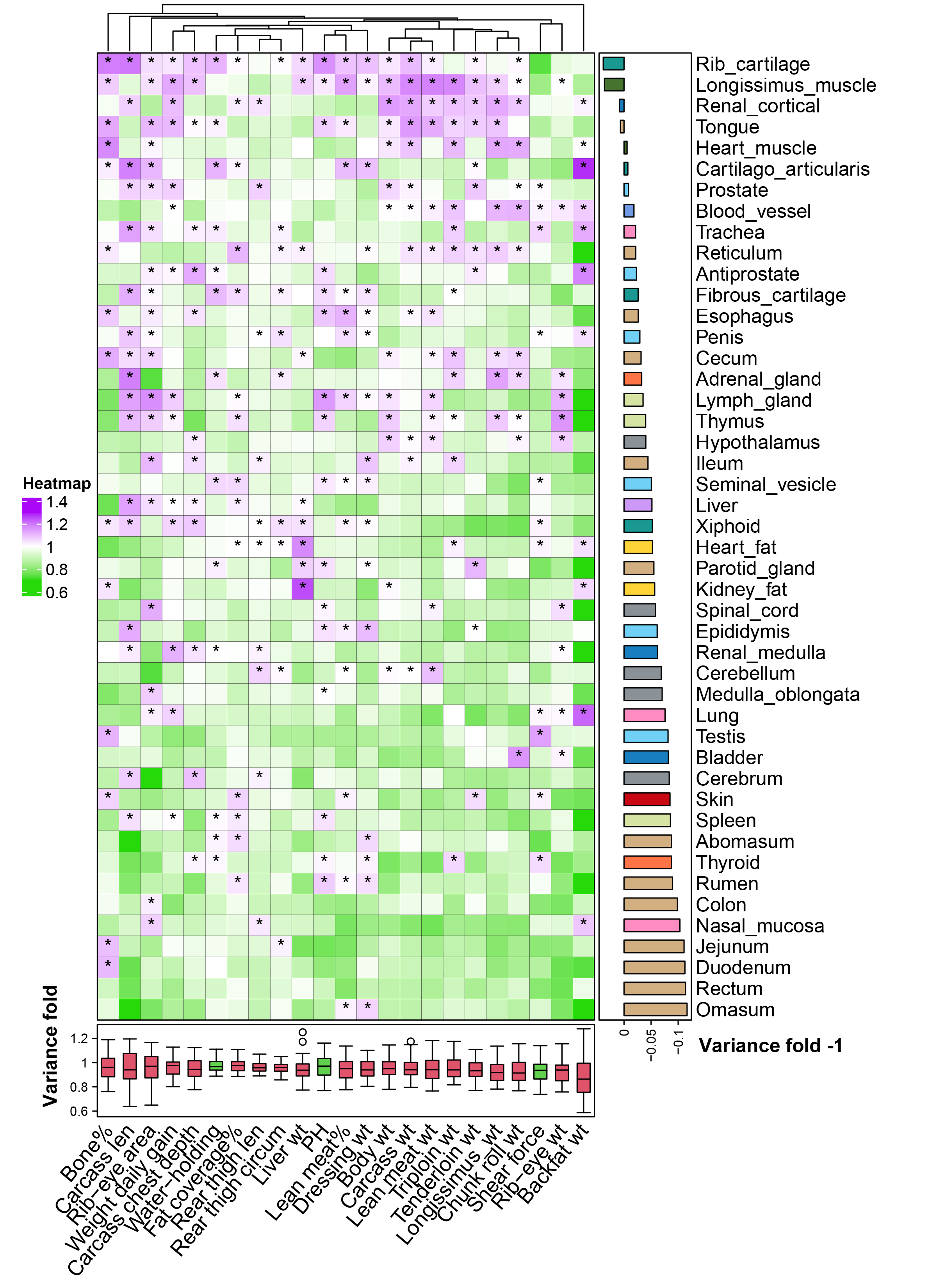

### FigureS34.tif

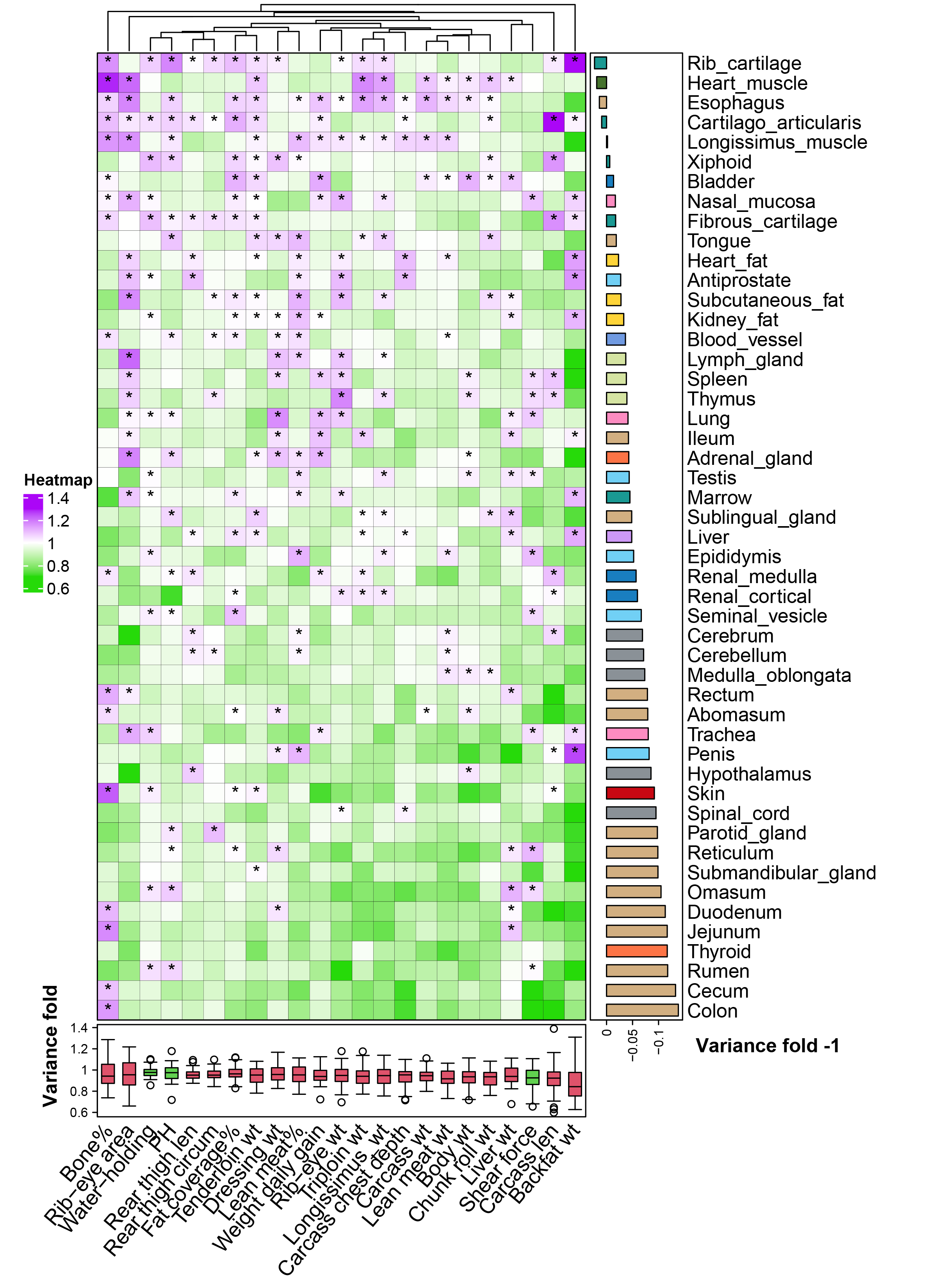

### FigureS35.tif

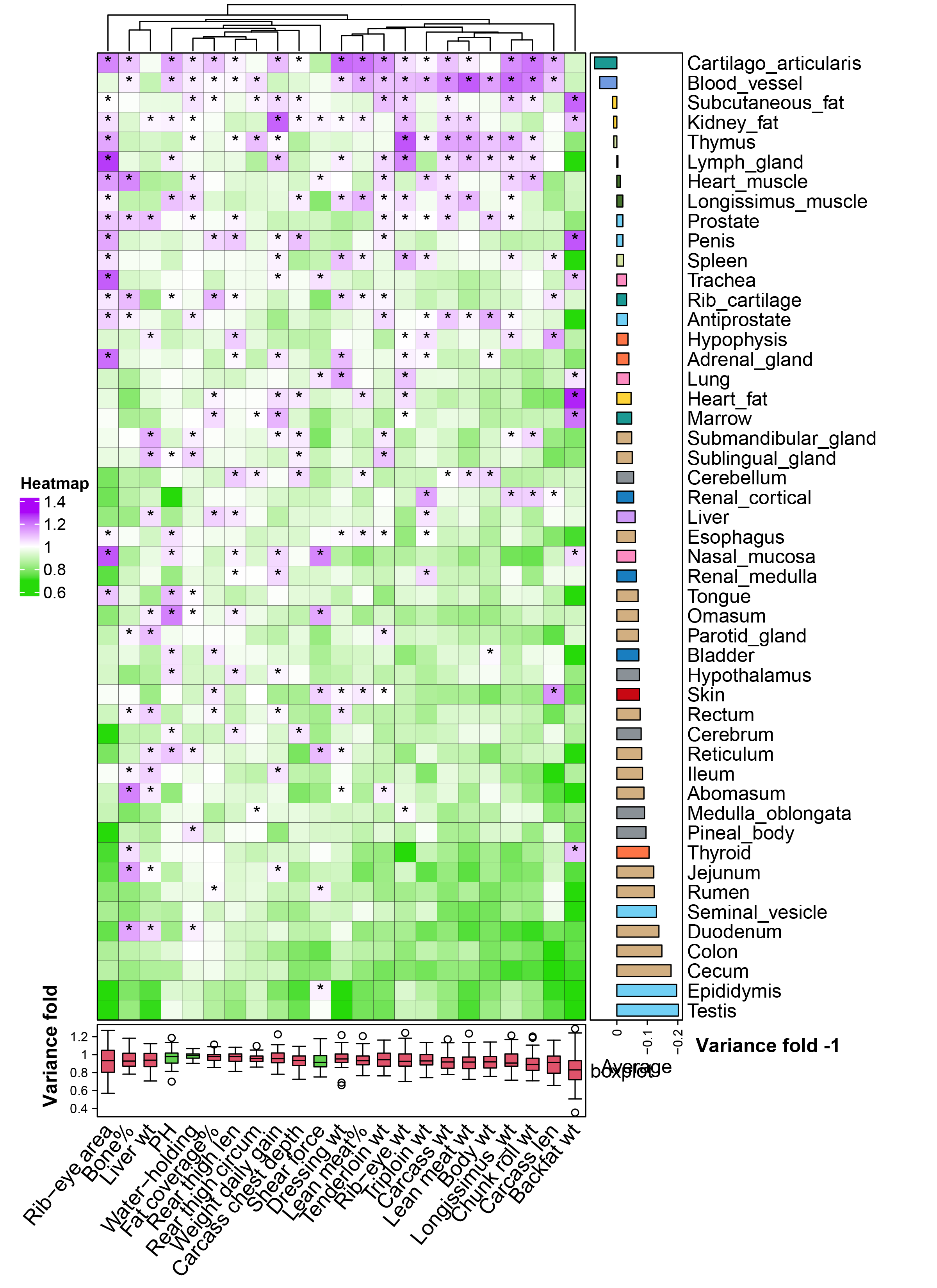

### FigureS36.tif

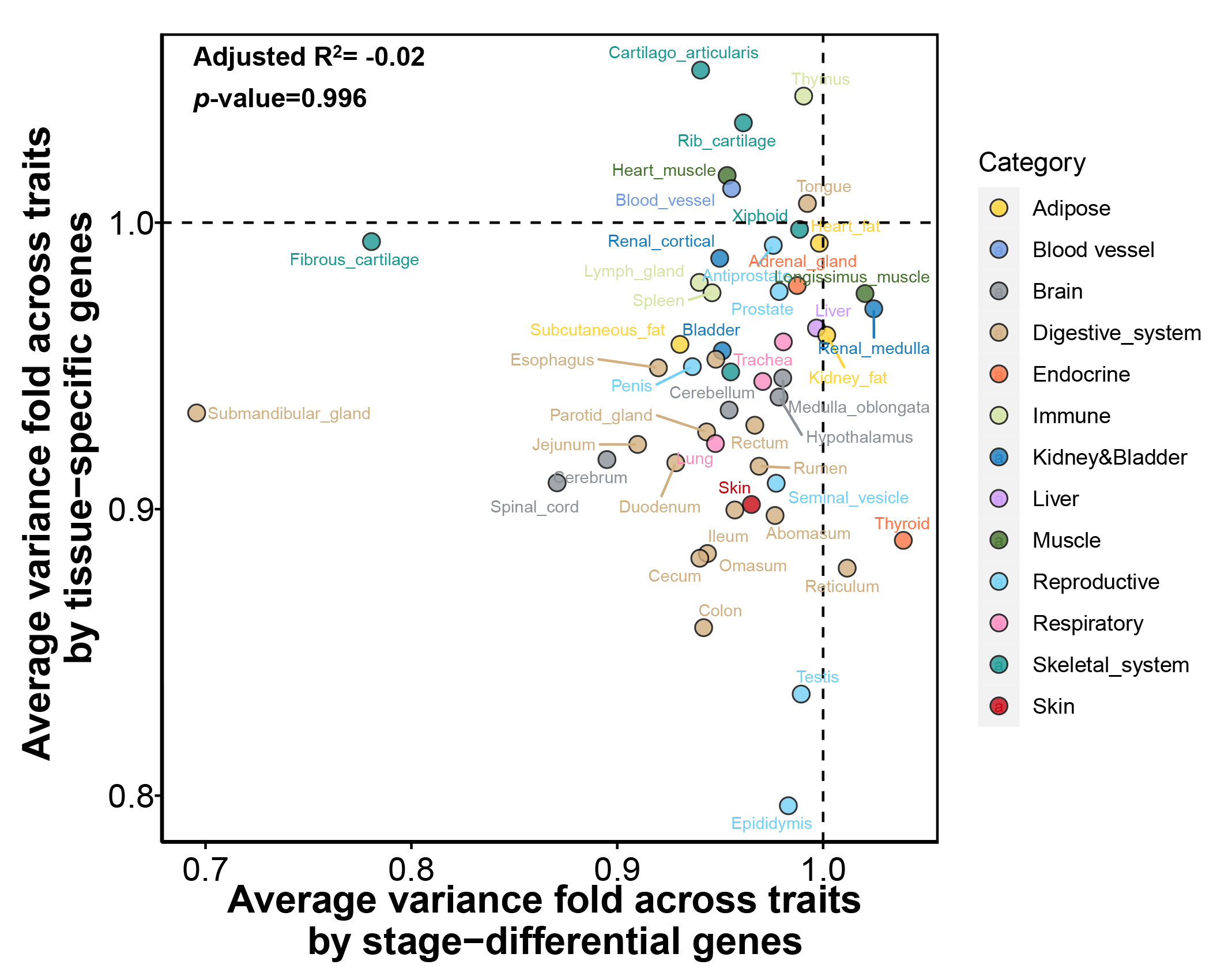
